# Aberrant expression prediction across human tissues

**DOI:** 10.1101/2023.12.04.569414

**Authors:** Florian R. Hölzlwimmer, Jonas Lindner, Georgios Tsitsiridis, Nils Wagner, Francesco Paolo Casale, Vicente A. Yépez, Julien Gagneur

## Abstract

Despite the frequent implication of aberrant gene expression in diseases, algorithms predicting aberrantly expressed genes of an individual are lacking. To address this need, we compiled an aberrant expression prediction benchmark covering 8.2 million rare variants from 633 individuals across 49 tissues. While not geared toward aberrant expression, the deleteriousness score CADD and the loss-of-function predictor LOFTEE showed mild predictive ability (1-1.6% average precision). Leveraging these and further variant annotations, we next trained AbExp, a model that yielded 12% average precision by combining in a tissue-specific fashion expression variability with variant effects on isoforms and on aberrant splicing. Integrating expression measurements from clinically accessible tissues led to another two-fold improvement. Furthermore, we show on UK Biobank blood traits that performing rare variant association testing using the continuous and tissue-specific AbExp variant scores instead of LOFTEE variant burden increases gene discovery sensitivity and enables improved phenotype predictions.

## Main

Aberrant gene expression, gene expression levels outside the physiological range, is a frequent cause of diseases. Aberrant underexpression of tumor suppressor genes and aberrant overexpression of oncogenes are hallmarks of oncogenesis^1,2^. Moreover, aberrant gene expression is a frequent cause of rare inheritable disorders^3–8^ and contributes to risks for common disease-associated traits^9^.

Statistical methods to call expression outliers from RNA-seq data^10–13^ applied to large cohorts have enabled investigating the genetic basis of aberrant expression. Rare variants have been found to be associated with expression outliers^14^. Specifically, rare variants including rare structural variants and rare variants likely triggering nonsense-mediated decay such as premature stop codons, frameshift, and splice-disrupting variants are enriched among underexpression outliers^6,14,15^. Moreover, rare structural variants, notably gene duplication, were found enriched among overexpression outliers^14,15^.

Building on these findings, algorithms have been developed to prioritize which variants may be the genetic cause of an expression outlier identified in an RNA-seq sample given the matched genome^14,15^. However, there is no algorithm predicting aberrant expression caused by genetic variants. A model predicting aberrant expression in multiple tissues and generalizing to unseen variants could improve our ability to identify high-impact rare variants in large genomic cohorts which in turn would aid in the identification of disease-associated genes and pinpointing disease-causal variants.

To address this unmet need, we here establish a benchmark for rare-variant based prediction of aberrant gene expression across human tissues (Fig. 1). While underexpression outliers can be assumed to have strongly impaired or entirely lost function, the functional consequence of an overexpression outlier is less clear, as it could as well result in a gain of function. We therefore primarily focussed in this study on underexpression outliers and developed AbExp, a machine learning model predicting aberrant underexpression in multiple tissues from an individual’s rare genetic variants. By integrating various variant annotations with tissue-specific isoform proportions and expression variability, AbExp significantly outperforms other loss-of-function annotators that were not explicitly developed for aberrant underexpression prediction. We demonstrate how AbExp can be used in rare-variant association testing as well as phenotype prediction on the UK Biobank dataset using 40 blood traits. Finally, we show that when gene expression measurements from clinically accessible tissues are available, AbExp scores can be integrated to improve the prediction of aberrant expression in non-accessible tissues.

**Figure 1:**
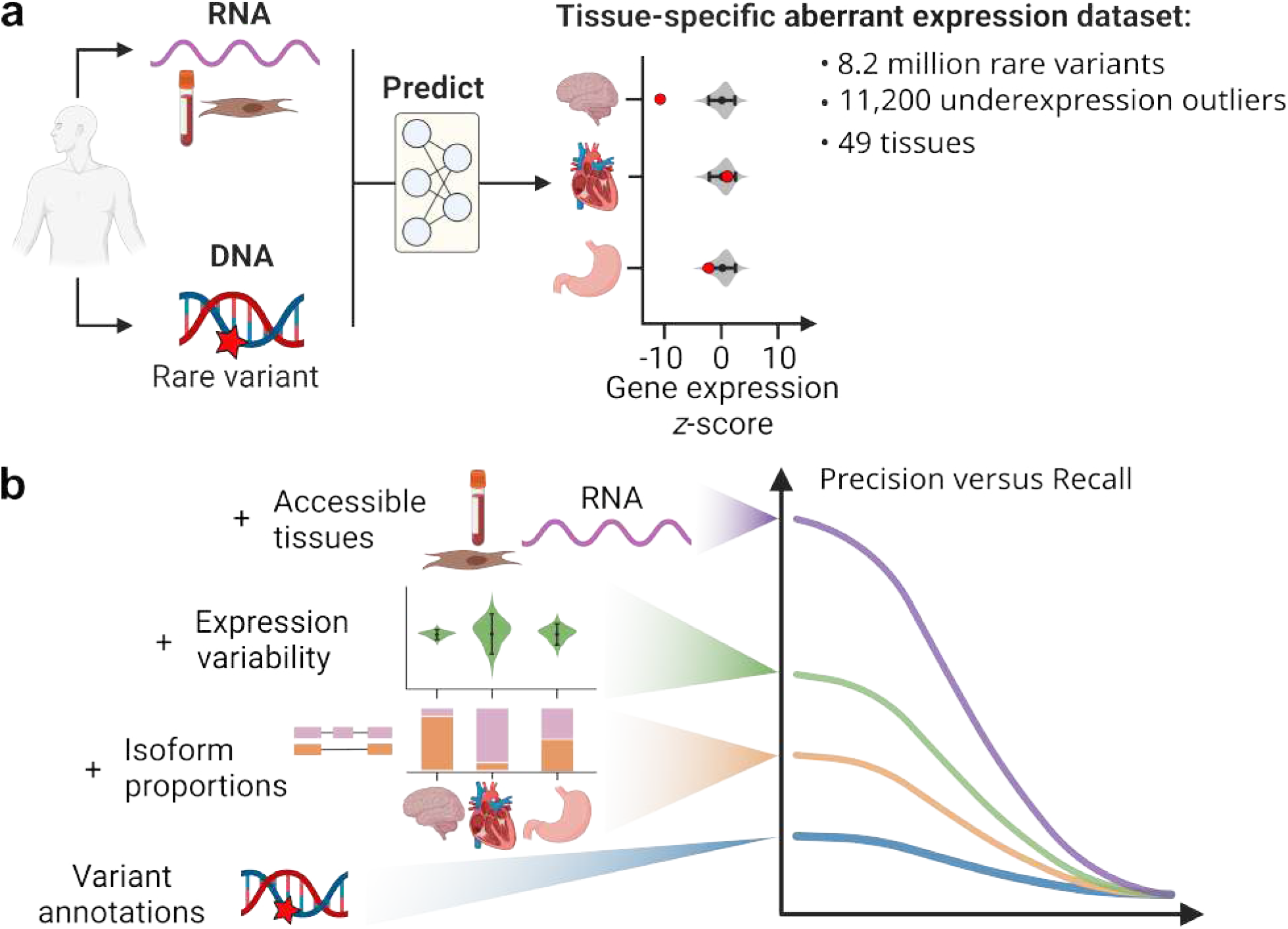
Study design. **(a)** We aimed to predict whether protein-coding genes are aberrantly underexpressed across 49 human tissues based on DNA and, optionally, RNA-seq data of clinically accessible tissues. Therefore, we created a benchmark for aberrant underexpression prediction by processing 11,215 RNA-seq samples from 633 individuals across 49 tissues from GTEx. This yielded 11,200 underexpression outliers out of 99 million assessable gene-sample pairs (0.01%). **(b)** Assessing various variant and tissue annotations, we found that predictions could be significantly improved by weighting variant effects with tissue-specific isoform proportions and incorporating the expression variability of a gene. Further integration of expression measurements from clinically accessible tissues led to another two-fold improvement.

## Results

### A benchmark dataset of underexpression outliers across 49 human tissues

We first set out to predict which protein-coding genes are aberrantly underexpressed in which tissues given an individual’s genome. We focussed on protein-coding genes because they represent the vast majority of transcripts implicated in human disease. Moreover, the degradation mechanisms of messenger RNAs, which are strongly coupled with translation notably via nonsense-mediated decay, substantially differ from the degradation mechanisms of non-coding RNAs, a very heterogenous class of transcripts ^16^ whose degradation involves a large variety of mechanisms^17–19^. To this end, we created a benchmark dataset using the aberrant expression caller OUTRIDER^11^ on 11,215 RNA-seq samples with paired whole- genome sequencing data of the Genotype-Tissue Expression dataset^20^ (GTEx v8), spanning 49 tissues and 633 individuals. For the whole-genome sequencing data, we selected GTEx version 7 since, unlike for version 8, structural variant calls were available. Although the choice against GTEx version 8 reduced the number of underexpression outliers by 20%, it allowed us to consider structural variants, which are important determinants of aberrant expression^15^. In each tissue, we restricted the analysis to all protein-coding genes with an average read-pair count of at least 450, an estimated minimal coverage required to detect 50%-reduction outliers^6^. Overall, 18,171 protein-coding genes out of 18,563 (97.9%) showed sufficient average coverage in at least one tissue.

We defined a gene to be aberrantly underexpressed if OUTRIDER reported a false discovery rate (FDR) lower than 0.05 and the expression was lower than expected (Methods). We further removed samples with more than 20 outliers since such a high number of outliers could reflect an unreliable fit of OUTRIDER due to technical reasons, for instance low sequencing depth^11^, or biological reasons, for instance an ancestry underrepresented in the cohort^6^. This exact cutoff combination on FDR and on the number of outliers per sample was selected among 12 possible combinations based on the generalisation performance of simple outlier predictors trained on established outlier-associated features^15^ (suppl. Fig. S1). Application of these cutoffs led to a benchmark dataset of 11,200 underexpression outliers occurring in 3,240 genes and 10,999 samples (suppl. Fig. S2), amounting to nearly one underexpression outlier per sample on average. With 99 million non-outliers in the dataset (gene-sample pairs assessed by OUTRIDER but not identified as outliers), the proportion of underexpression outliers is extremely low at just 0.01%. This creates a highly imbalanced prediction task.. A detailed overview of dataset statistics including how many samples and genes remained after each filtering step can be seen in suppl. table T1.

### Integrating rare variant annotations to predict underexpression outliers across tissues

We considered predicting aberrant expression for all 49 tissues for any protein-coding gene from rare variants, here defined as variants occurring in at most two GTEx individuals and with a gnomAD minor allele frequency of less than 0.1%. We reasoned that above 0.1% minor allele frequency, variants are unlikely to cause an expression outlier since the average frequency across samples of underexpression outliers among genes with sufficient RNA-seq coverage is about 0.01%. Following observations from Ferraro and colleagues^15^, we focussed on single- nucleotide variants and short insertions and deletions located within the gene and up to 5,000 bp around the gene to fully cover promoter and transcription termination regions. Moreover, we considered structural variants up to 1 megabase 5’ of the gene^15^. We do not exclude that some single nucleotide variants and short indels could also have strong effects when located further than 5,000 bp of a gene. However, we did not consider this class of variants because sequence- based models capturing distal enhancer effects are lacking^21^.

We first investigated whether existing variant annotation tools, that were not developed for predicting aberrant underexpression, showed informative signals. Among the variant consequences annotated by Ensembl VEP, we found frameshifts, variants affecting start and stop codons, and splicing variants to be strongly enriched among underexpression outliers, consistent with previous reports^6,14,15^ (Fig. 2a, suppl. Fig. S3). Variants including frameshifts, splice-variants, and stop-gains, which introduce premature stop codons are known to be strongly enriched within gene underexpression outliers as these often trigger nonsense- mediated decay (NMD)^14^. LOFTEE is a tool that predicts a high-confidence subset of loss-of- function associated variants, notably variants likely to trigger NMD, by implementing filters such as removing stop-gained and frameshift variants that are within 50 bp of the end of the transcript, or variants that affect splicing only in UTRs^22^. In GTEx more than 23% of the aberrantly underexpressed genes had a LOFTEE variant, compared to non-outliers that had a LOFTEE variant in less than 0.1% of the cases (Fig. 2a).

**Figure 2:**
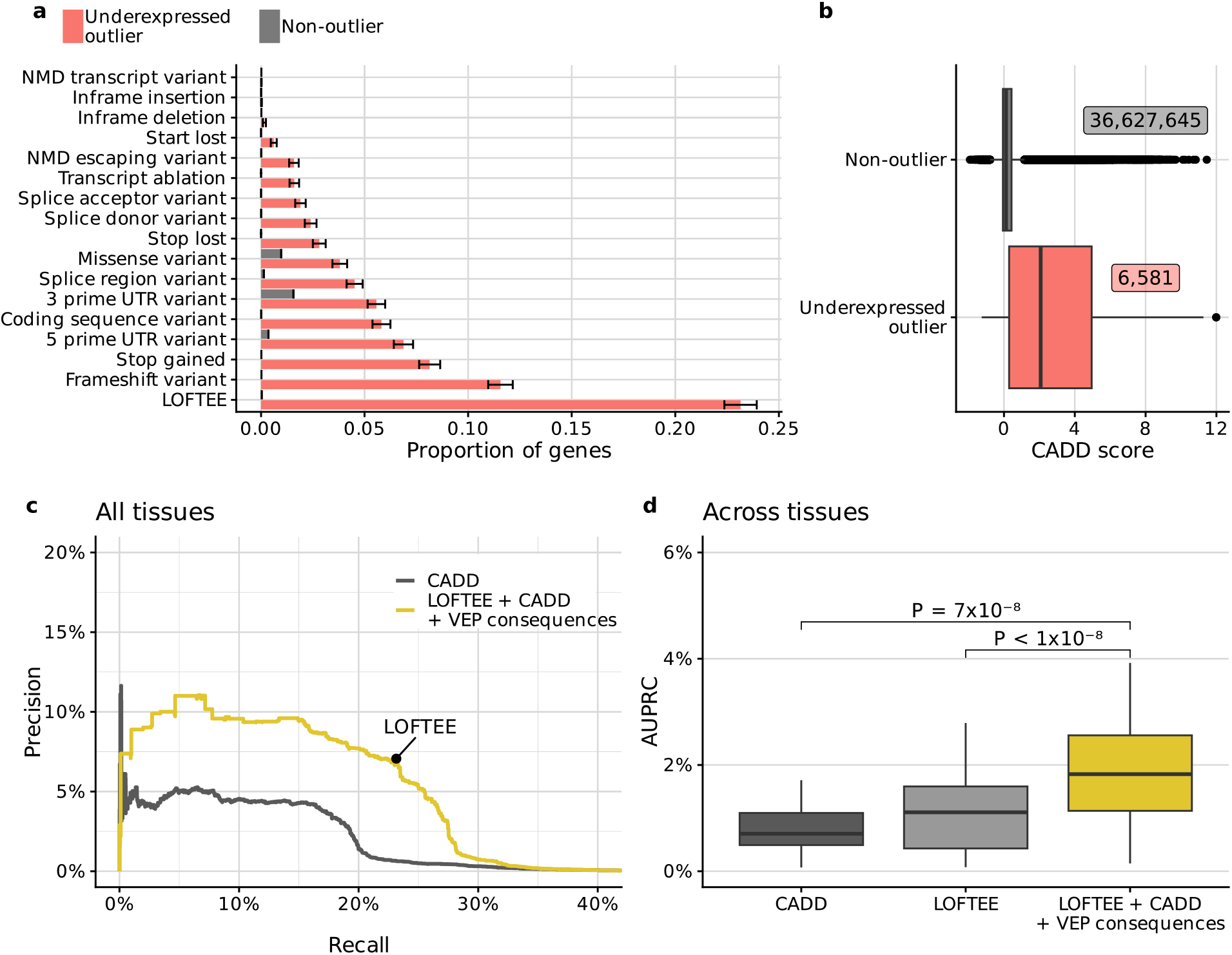
Integrating rare variant annotations to predict underexpression outliers across tissues. **(a)** Proportion of underexpressed outliers (red), and non-outliers (gray) with a rare variant of a given annotation (rows, Methods). Error bars mark 95% binomial confidence intervals. Notably, rare variants likely to trigger nonsense- mediated decay are strongly represented in underexpressed genes. **(b)** Distribution of CADD scores in different outlier classes. The numbers of data points are labeled for each box. **(c)** Precision-recall curve for all tissues combined. LOFTEE shows up as a single point because it is a binary filter. **(d)** Distribution of average precision (AUPRC) across 27 GTEx tissue types (Methods). *P*-values were obtained using a paired Wilcoxon test. A non-linear model based on LOFTEE, CADD and VEP annotations significantly outperforms existing methods. **For all boxplots:** Center line, median; box limits, first and third quartiles; whiskers span all data within 1.5 interquartile ranges of the lower and upper quartiles.

Moreover, we hypothesized that the deleteriousness score CADD^23^ could also be predictive of aberrant gene expression. The advantage of CADD, which was trained to distinguish between simulated de novo variants and variants that have arisen and become fixed in human populations, is that it provides a score for any variant. We found that the median CADD score of rare GTEx variants was ∼17 times higher among underexpressed genes than among non- outliers (Fig. 2b).

Despite these enrichments, LOFTEE and CADD by themselves showed limited predictive value. Using the sole LOFTEE-positive variants recalled 23.2% of the underexpression outliers at a precision of 7.1% (Fig. 2c). Besides a small spike to 11.6% precision at 0.2% recall, CADD never reached the same precision nor the same recall as LOFTEE (Fig. 2c). Next, we trained a non-linear model that integrated all the above-mentioned features to quantitatively predict the OUTRIDER *z*-score (Methods). Predicting the underlying quantitative z-scores turned out to lead to better classifiers than directly predicting the binary classes of outliers and non-outliers. Moreover, ranking based on the predicted *z*-scores uniformly outperformed ranking based on CADD scores on held-out data (Fig. 2c). The integrative model reached the same precision at the same recall as filtering for LOFTEE variants with the added value of providing a continuous score, thereby allowing for applying more stringent cutoffs to yield a higher precision (up to 11%, Fig. 2c). Lastly, the advantage of the integrative model over CADD and LOFTEE was observed in aggregate (Fig. 2c) as well as across individual tissues according to the average precision (measured by the area under the precision-recall curve here and elsewhere, AUPRC, Fig. 2d).

Additionally, we adapted Watershed, a method that was originally designed to identify the most likely variant responsible for an observed expression outlier, to fit our task^15^. Watershed is a Bayesian approach that relies on strong modeling assumptions. In particular, Watershed represents the outlier status as a categorical variable and models variant features independently. Using the Watershed features, we found that predicting tissue-specific *z*-scores with a non-linear model largely outperforms the Watershed-derived outlier predictor (suppl. Fig. S4). Altogether, these investigations demonstrate that outlier prediction benefits from a direct and quantitative treatment and from flexible, non-linear machine learning models.

### Accounting for tissue-specific isoform expression improves predictions

By construction, the predictions of this first model were independent of the tissue as neither the variant annotations nor the considered transcript isoforms were tissue-specific. However, since the transcript isoforms of a gene are often expressed at different proportions across tissues, variants can have tissue-dependent effects^24^. For example, ENST00000358514, the canonical transcript of *PSMB10*^25^, was estimated to generate only about 4% of *PSMB10* total gene expression in putamen (Methods). The vast majority (91%) of *PSMB10* gene expression in putamen was attributed to another transcript, ENST00000570985. Conversely, in fibroblasts, the canonical transcript contributed to nearly 48% of the total gene expression. Exon 4 is not included in the transcript ENST00000570985 but is included in the canonical transcript ENST00000358514, explaining why a frameshift variant in exon 4 was associated with a high impact on gene expression in cultured fibroblasts but showed a limited effect in putamen (Fig. 3a,b).

**Figure 3:**
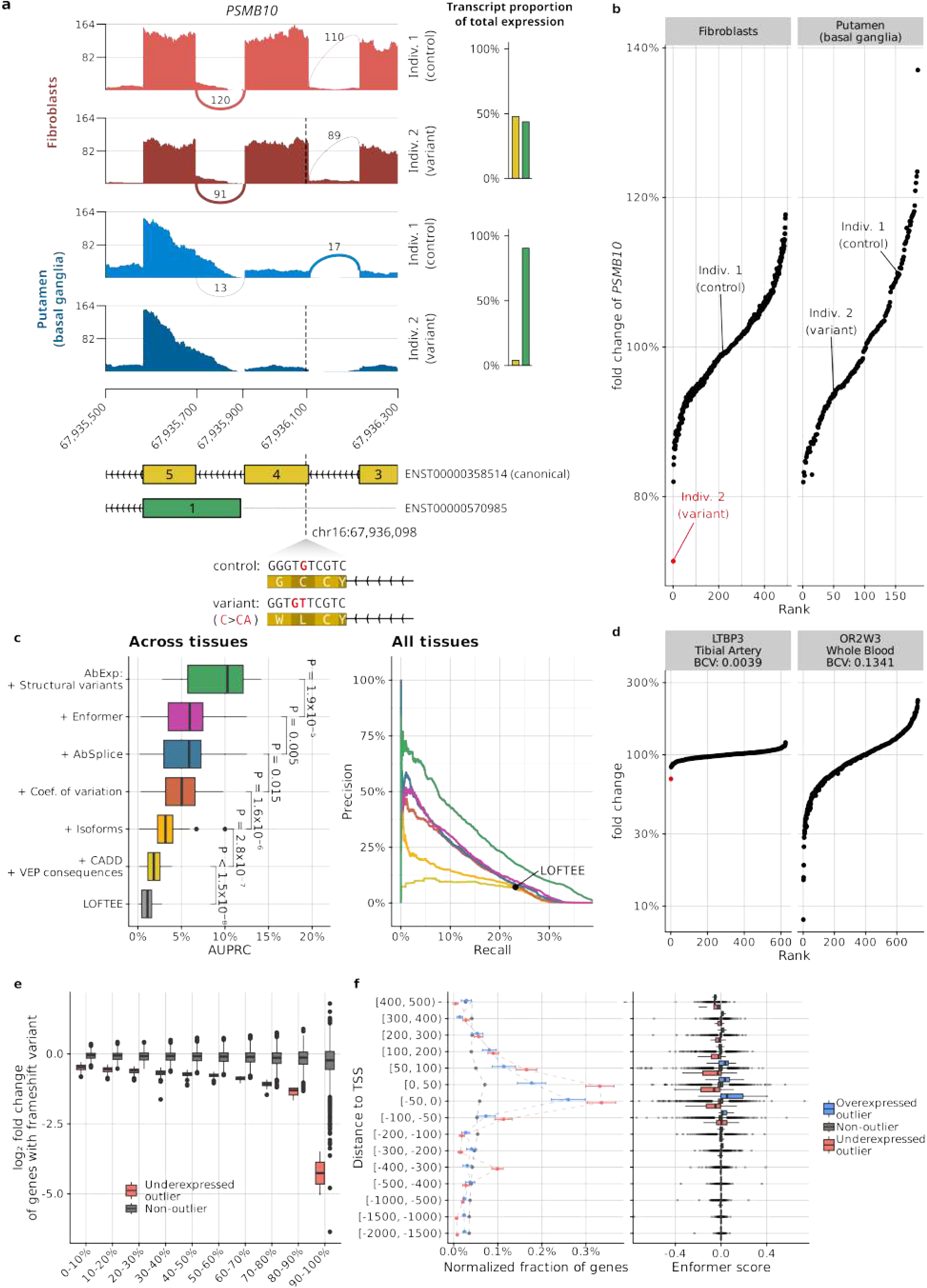
Tissue-specific features improve the model prediction. **(a)** Sashimi plot of *PSMB10* for two individuals, one carrying no rare variant in this region (control, upper tracks), and one carrying a heterozygous frameshift variant (dashed line and lower tracks), in cultured fibroblasts (top) and putamen (bottom). The frameshift variant is located on exon 4 which is included on the canonical transcript (ENST00000358514) but not on transcript ENST00000570985. On the right, the bar plots show the transcript expression proportions on each tissue on average across GTEx. In fibroblasts, the rare variant is associated with an approximately 25% reduction of RNA-seq coverage in this window whereas in putamen no major RNA-seq coverage change is observed. **(b)** Fold change of gene expression against normalized gene expression rank for *PSMB10* in fibroblasts and putamen (basal ganglia) brain tissues. *PSMB10* is an expression outlier (red) in individual 2 in fibroblast but not in putamen, consistent with the rare variant triggering nonsense-mediated decay and leading to a strong gene expression reduction in the tissue for which the exon 4- containing transcript is the major isoform. **(c)** Left: Distribution of average precision (AUPRC) across 27 GTEx tissue types. *P*-values were obtained using the paired Wilcoxon test. AbExp combines various variant and tissue annotations to predict aberrant gene expression and outperforms LOFTEE by about sevenfold. Right: Precision-recall curve for all tissues combined. **(d)** Fold change of gene expression against normalized gene expression rank for *LTBP3* in tibial artery, an autosomal recessive gene whose defect can lead to dental anomalies and short stature ^26^, and for *OR2W3* in blood, an olfactory gene whose defect should not impair the viability of an individual ^27^. Expression outliers are highlighted in red. *LTBP3* is tightly regulated with a fold change range of ± 20% among non-outliers. The individual marked in red carries a heterozygous frameshift variant that associates with 30% reduction and which is detected as an outlier. In contrast, *OR2W3* shows very large variations where individuals with 30% reductions are not outliers. **(e)** Distribution of gene expression fold changes for underexpression outliers (red) and non-outliers (grey) harbouring a frameshift variant per decile of expression variability (biological coefficient of variation). **(f)** Left: Proportion of underexpressed outliers (red; n=2,051), non-outliers (gray; n=2,749,843), and overexpressed outliers (blue; n=1,177) with a rare variant within a certain distance of the transcription start site (TSS), normalized by the length of each distance interval. Error bars mark 95% binomial confidence intervals. Right: Distribution of Enformer scores in different outlier classes among rare variants within a certain distance of the TSS. **For all boxplots:** Center line, median; box limits, first and third quartiles; whiskers span all data within 1.5 interquartile ranges of the lower and upper quartiles.

Generally, we found that only 30% of the canonical transcripts contributed more than 90% of the total expression for their respective genes and that as much as 18% of the canonical transcripts contributed to less than 10% of the total expression for their respective genes (suppl. Fig. S5). Therefore, relevant information is lost when considering the variant consequence assigned to a single isoform, even if it is annotated as the canonical one. To address this issue, we calculated the isoform composition in every tissue and weighted the VEP consequences and LOFTEE classification of each variant by the proportion of affected transcripts per gene and tissue (Methods). Training the model using these tissue-specific weighted annotations increased the average precision by 58% to reach 3.2% in median across tissues (Fig. 3c).

### Incorporating the tissue-specific gene expression variability further improves predictions

Similar to other statistical models for RNA-seq data, OUTRIDER includes a measure of gene expression variability called the biological coefficient of variation^11,28^. The biological coefficient of variation is used instead of the plain coefficient of variation to account for sampling noise, which particularly affects low RNA-seq read counts^28^. In our benchmark dataset, the biological coefficient of variation captures the expression variability of genes per tissue across the GTEx population. We reasoned that the same expression fold changes could cause an expression outlier for a gene with low expression variability but not for a gene with high expression variability. Indeed, we observed that the minimal fold-change among expression outliers decreased with the biological coefficient variation (suppl. Fig. S6). Therefore, a given relative reduction in gene expression can lead to aberrant expression in one gene or tissue but not necessarily in another. For instance, a 30% reduction of the gene *LTBP3* in tibial artery expression resulted in an outlier. In contrast, a 30% expression reduction in blood of *OR2W3* would not lead to an outlier as *OR2W3* shows a very large gene expression variation in blood ranging between 10% and 230% (Fig. 3d). It is not surprising that *OR2W3*, one of the over 800 human olfactory receptor genes^27^ for which dysfunction is likely benign, exhibits more expression variability than *LTBP3*, a gene whose dysfunction is associated with dental anomalies and short stature^26^.

Next, we aimed to improve our underexpression outlier predictor by adjusting for expression variability. We first considered modeling expression fold-changes from variants and then converting the predicted fold-changes into a *z*-score, under the assumption that variants affect gene expression fold-changes independently of the expression variability. Using variants likely triggering NMD to test this assumption, we noticed however that the same class of variants associated with lower fold-changes among genes with lower expression variability (Fig. 3e, suppl. Fig. S7), perhaps because genes with low expression variability are subject to regulatory buffering mechanisms^29,30^. Therefore, we opted for a more general modeling approach in which the biological coefficient of variation is provided as an input feature along with variant annotations to a non-linear model predicting the *z*-score. This model increased the performance by more than 63% to 5.0% average precision (median across tissues, Fig. 3c). Altogether, these results show that gene expression variability must be taken into account when predicting aberrantly expressed genes, and that predicting z-score rather than fold-change is more relevant for variant interpretation.

### Contribution of aberrant splicing variants, core promoter variants, and structural variation

Aberrant splicing isoforms often contain a premature termination codon and as a consequence, yet not always, are degraded by NMD^31^. Our model already included splice site information, as it was based on LOFTEE, CADD, and tissue-specific isoform weighting of VEP annotations. However, these features ignored splice sites that are not part of the genome annotations upon which these tools are built. We have recently developed a model called AbSplice that predicts aberrant splicing across tissues using a more comprehensive map of splice sites, including unannotated weak splice sites, and their tissue-specific usage^32^. We found that AbSplice scores were in median 10 times higher among underexpression outliers than among non-outliers (suppl. Fig. S8a). Integrating AbSplice scores significantly increased the performance to 5.9% average precision in median across tissues. In contrast, adding SpliceAI did not show any significant improvement (suppl. Fig. S8b), in agreement with AbSplice improving over SpliceAI for predicting tissue-specific aberrant splicing^32^.

To better capture variants affecting transcription, we considered Enformer^33^, a state-of-the-art deep learning model that predicts thousands of transcription-related genome-wide assays including 635 CAGE tracks of the ENCODE project, based on a 200-kb sequence context. Building on previous work^21^, we mapped Enformer-predicted CAGE tracks to gene expression for the 49 GTEx tissues with regularized linear regression models (Methods). We found that using elastic net models, rather than ridge regression, and modeling effects on canonical isoforms, rather than on mixtures of tissue-specific isoforms, resulted in the best median AUPRC across tissues (suppl. Fig. S9 a,b). This Enformer-based model was predictive for expression outliers in the correct direction but only in the close vicinity of the transcription start site (−50 bp to +200 bp, Fig. 3f and suppl. Fig. S9c,d). On the one hand, this observation may reflect the biology of genetic determinants of aberrant expression, whereby variants with extreme effects on gene expression may primarily lie in the core promoter, which extends to 50 bp on either side of the transcription start site in eukaryotes^34^. Consistently, we found a higher proportion of outlier-associated variants in the core promoter region, independently of Enformer predictions (Fig. 3f). On the other hand, the poor precision of Enformer beyond this narrow window around the transcription start site may reflect the difficulty that Enformer has in capturing long-range effects^21^. Despite the strong enrichment in underexpressed outliers, adding Enformer to the AbExp feature set increased the median AUPRC only minimally to 6.0% across tissues, albeit significantly.

We next considered the inclusion of structural variants overlapping transcripts (Methods). Transcript ablations, which could be inferred in GTEx from the structural variant deletion calls, were especially predictive of underexpression. Overall, 33 out of the 43 GTEx individuals harboring transcript ablations also showed an expression outlier in at least one tissue. Including structural variants into the model resulted in a large gain in precision among the top-ranked predictions and increased the average precision to 10.2% in median across tissues (Fig. 3c). We also considered structural variants up to 1 megabase 5’ of the transcription start site, which were enriched for expression outliers in agreement with Ferraro et al.^15^ (suppl. Fig. S10a). However, as the integration of these variants did not lead to a significant improvement (suppl. Fig. S10b) we did not consider the distant structural variants further.

In the following, we refer to the model which integrates all significant features mentioned so far as AbExp. AbExp takes as input a set of variants within 5,000 bp of any annotated transcript of a protein-coding gene and returns a predicted z-score for each of the 49 tissues. For user convenience, we furthermore suggest a high-confidence cutoff (AbExp < -3.84) corresponding to 50% precision and 7.8% recall on our benchmark data, and a low-confidence cutoff (AbExp < -1.64) corresponding to 20% precision and 21.5% recall. The high-confidence cutoff leads to about 1 positive prediction every 6 GTEx samples, whereas the low-confidence cutoff leads to about 1.2 positive predictions per GTEx sample, which is approximately the number of outliers per sample (1.02). AbExp performs especially well in predicting the impact of NMD-associated variants, outperforming other methods in these thanks to a more effective ranking (suppl. Fig. S11). However, AbExp cannot reliably predict the impact of variants in categories rarely associated with expression outliers in GTEx, such as missense and UTR variants. Larger datasets and complementary models will be needed to improve on these variant categories.

Although our primary focus was on predicting underexpression outliers, we leveraged the fact that AbExp quantitatively predicts z-scores to assess its performance for overexpression outliers (suppl. Fig. S12). We found that the performance for overexpression outlier prediction was substantially lower compared to underexpression outliers (median AUPRC 3.4% across tissues). By far, the major predictive features were structural variants, yet Enformer and isoform proportions also contributed significantly. Not surprisingly, LOFTEE, CADD, AbSplice, and other loss-of-function associated variant effects were not predictive for overexpression. These results align with the variant category enrichments reported by Ferraro et al., which showed that duplications and variants near the TSS were the only categories specifically enriched among overexpression outliers, but not among underexpression outliers^15^. Altogether, these results show that AbExp can also be employed for over-expression outlier prediction, yet with a much lower performance than for underexpression outliers.

### AbExp replicates on independent datasets

We next assessed how AbExp performance replicated on two independent datasets. The first dataset consisted of 295 individuals suspected to be affected by a mitochondrial disorder^6^ with whole-exome sequencing data paired with RNA-seq from fibroblasts. The second dataset consisted of 233 whole-genome sequencing samples with RNA-seq from iPSC-derived motor neurons from the AnswerALS research project^35^. Structural variant calls, and thus transcript ablation calls, were not available on either dataset. Also, we did not compute Enformer scores due to computational constraints and its limited added value for outlier prediction. Moreover, we observed that the recall for all methods was twice as low on the ALS dataset than in GTEx and in the mitochondrial disorder dataset (suppl. Fig. S13), perhaps because of poorer expression outlier calls, a stronger role of epigenetic and trans-regulatory effects, or combinations thereof. Taking these differences into account, our results on those two independent datasets were in agreement with the evaluation on GTEx. We found that AbExp without transcript ablation annotation significantly outperformed LOFTEE and CADD by two to three times larger average precision (suppl. Fig. S13). Here too, AbExp without structural variants and Enformer allowed for slightly better precision at the same recall than LOFTEE filtering, while offering a continuous score allowing for reaching much higher precisions.

One concern of having used the biological coefficient of variations in the GTEx benchmark is that they were computed on the very same RNA-seq data as those used to compute the ground truth expression outliers. Nevertheless, including the biological coefficient of variation computed on GTEx significantly improved the predictions on the mitochondrial disorder dataset and was on par on the ALS dataset (suppl. Fig. S13), showing that the biological coefficient of variation contributes with independent information to outlier prediction.

### AbExp prioritizes deleterious variants

Having established and independently validated AbExp as an aberrant expression predictor, we next assessed its potential for identifying deleterious variants. To this end, we first considered the capacity of AbExp at distinguishing pathogenic from benign and likely benign variants as reported in the ClinVar database (Methods). For each variant, we retained the minimum AbExp score, i.e. the most impactful underexpression prediction across all tissues. We found that AbExp improved over CADD, LOFTEE, and AbExp without biological coefficient of variation in the high precision range with precision exceeding 99% up to 47% recall (Fig. 4a). However, at lower precision and higher recall both CADD and LOFTEE outperformed AbExp.

**Fig 4:**
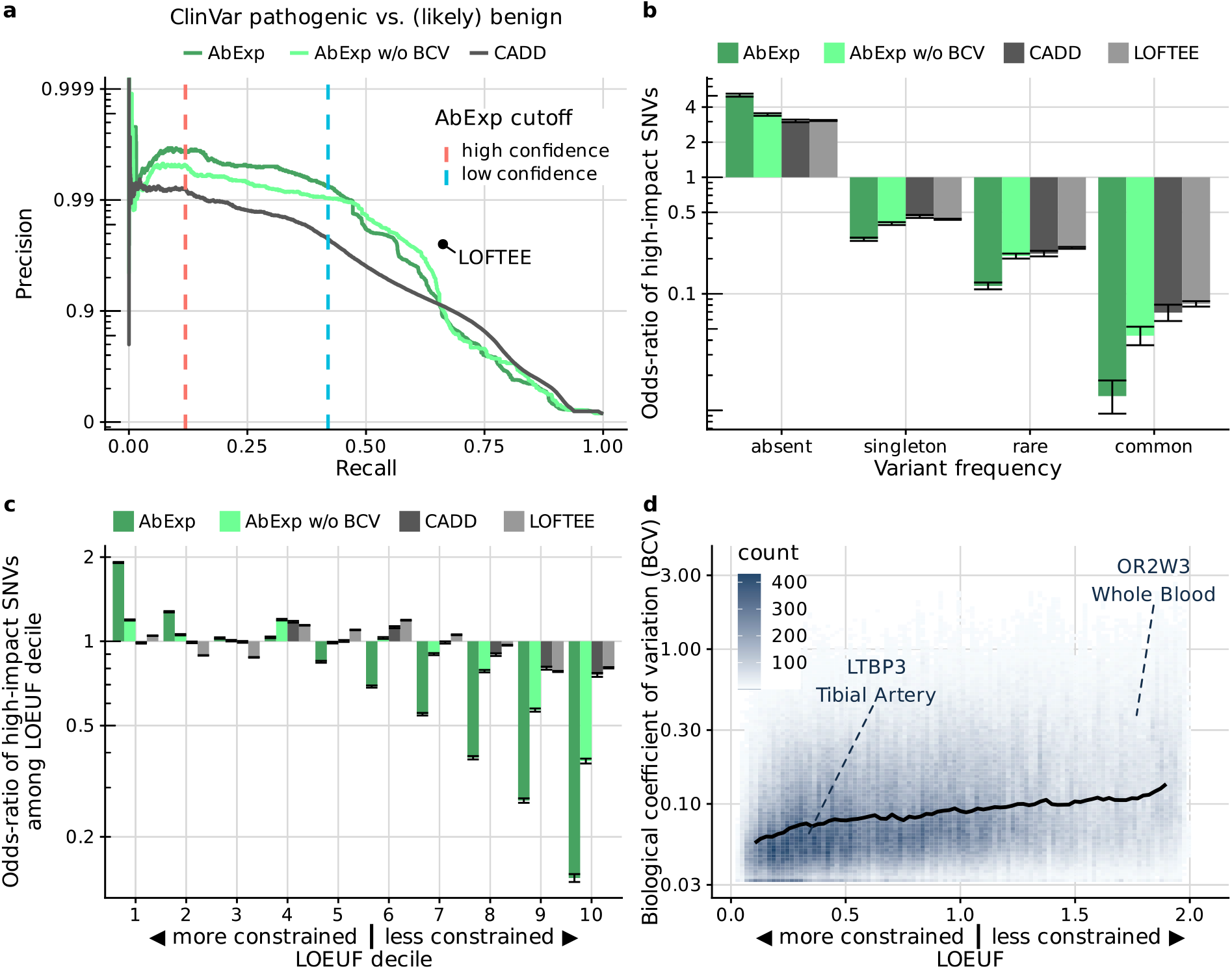
AbExp prioritizes deleterious variants. **(a)** Precision-recall curve of AbExp (trained with and without using the biological coefficient of variation), CADD, and LOFTEE on distinguishing pathogenic from benign and likely begin variants in ClinVar. LOFTEE as a binary predictor is shown as a single point. The dashed vertical bars denote the high and low confidence cutoffs of AbExp. **(b)** Odds-ratio of high-impact variants among absent, singleton, rare (MAF < 0.1%) and common SNVs in gnomAD for the models shown in **a**. The analysis is restricted to variants within 5 kb of protein-coding genes. Error bars show Wald 95% confidence intervals from logistic regression fits. The high-impact cutoffs for CADD and AbExp without BCV were set to match the quantile of the high-impact cutoff of AbExp. **(c)** As in **b** among SNVs absent in gnomAD as a function of gene LOEUF decile^22^. Genes with a high LOEUF are more tolerant to loss of function. Error bars as in **b. (d)** BCV versus LOEUF across all genes and tissues. The black line shows a running median between LOEUF and BCV highlighting the two genes from Fig. 2d. The autosomal recessive gene *LTBP3* has a low LOEUF, denoting a low loss-of-function tolerance. In contrast, the olfactory gene OR2W3 has a high LOEUF, denoting a large loss-of-function tolerance.

One limitation of this benchmark is that ClinVar submitters are using established bioinformatics tools including LOFTEE and CADD to identify pathogenic variants, introducing a bias in favor of those tools. To circumvent this issue, we next examined a dataset devoid of human annotations that could indicate that AbExp allows prioritizing deleterious variants. Reasoning that more frequent variants in the human population are less likely to be deleterious, we stratified all possible variants within 5 kb of protein-coding genes into four frequency categories: absent in gnomAD, singletons (found only once), rare (more than once but less than 0.1%) or common (more than 0.1%). AbExp was significantly and more strongly associated with these variant categories than LOFTEE, CADD, and AbExp without biological coefficient of variation (Fig. 4b). Moreover, focussing on variants absent in gnomAD, AbExp high-impact variants were more significantly enriched than alternative methods among highly genetically constrained genes^22^, i.e. for those genes for which one can expect the strongest phenotypic impact (first LOEUF deciles, Fig. 4c). Notably, when the biological coefficient of variation was not accounted for in AbExp, the relative enrichment on constrained genes was reduced (Fig. 4c). This is consistent with our observation that more genetically constrained genes showed lower expression variability (Fig. 4d), aligning with previous findings in primates^36^. This interpretation was also consistent when performing the enrichment analyses against LOEUF for further allele frequency categories. At the extreme, common variants classified as high-impact by AbExp were depleted in the first LOEUF decile and enriched in the last LOEUF decile (suppl. Fig. S14). Intermediate behaviors were observed for singletons (variants observed only once in gnomAD) and rare variants (non-singleton gnomAD variants with MAF <0.1%). Altogether, these results indicate that AbExp can be used as an informative variant prioritization algorithm.

### AbExp improves rare variant association testing and phenotype prediction

The rise of large exome-sequencing and genome-sequencing cohorts empowers rare variant association testing (RVAT) which helps pinpoint causal genes for traits^37,38^ and enables improved phenotype predictions, particularly among individuals showing extreme phenotypes^39^. RVAT consists of identifying genes within which the occurrence of likely high-impact variants are associated with a trait^40,41^. To this end, accurate prediction of high-impact variants can be advantageous, suggesting that AbExp may have the potential to improve RVAT. To test this hypothesis, we considered 40 continuous blood traits including high-density lipoprotein cholesterol, glucose, and urate levels (suppl. table T2) from the UK Biobank 200k exome release^42^. We chose blood traits for this proof-of-concept investigation since they are well- studied and frequently measured to diagnose and monitor chronic disease conditions. Moreover, RVAT methods are typically better calibrated on continuous traits as opposed to binary traits^43,44^.

To ease comparisons between variant annotations, we used linear regression as a common framework for rare variant association testing (RVAT). As a realistic baseline, we considered RVAT based on LOFTEE variants, similar to the Genebass study, a phenome-wide study leveraging UK Biobank data^38^. For this baseline model, gene-trait association was tested by regressing the trait against the number of LOFTEE variants. The second model leveraged that AbExp is both quantitative and tissue-specific. To this end, we considered the lowest AbExp *z*- score across all rare variants for each of the 49 tissues. The gene-trait association was tested by regressing the trait against the resulting 49 values. Two further models were considered by regressing against the minimum and against the median of the 49 values. To adjust for other relevant factors and effects due to common genetic variation, all four models included as covariates sex, age, the first 20 genetic principal components, a polygenic risk score predicting the trait, and common variants reported to be associated with the trait and located 250,000 bp around the gene (Methods). We fitted every model on two thirds of the dataset for gene-trait association discovery. Phenotype permutation analysis indicated that all models were calibrated (suppl. Fig. S15).

Using AbExp predictions for the 49 tissues, we identified in total 30% more gene-trait associations compared to the LOFTEE-based model (Fig. 5a), showing that AbExp can significantly (*P* = 5.2×10^−4^) improve RVAT-based gene discovery. Notably, association testing using the tissue-specific predictions outperformed aggregated forms of the AbExp score in most cases by finding more gene-trait associations (suppl. Fig. S16). In some instances, we could rationalize which tissues showed the most significant associations. This was the case for the gene encoding Apolipoprotein B, a major constituent of triglyceride-rich lipoproteins synthesized in the liver and whose predicted aberrant underexpression in the liver was found to be negatively associated with blood triglyceride levels (suppl. Table T3). However, the high correlation between the AbExp scores per gene across tissues can make the estimation of each individual tissue-specific coefficient unstable, and, therefore, their interpretation should be done with caution.

**Figure 5:**
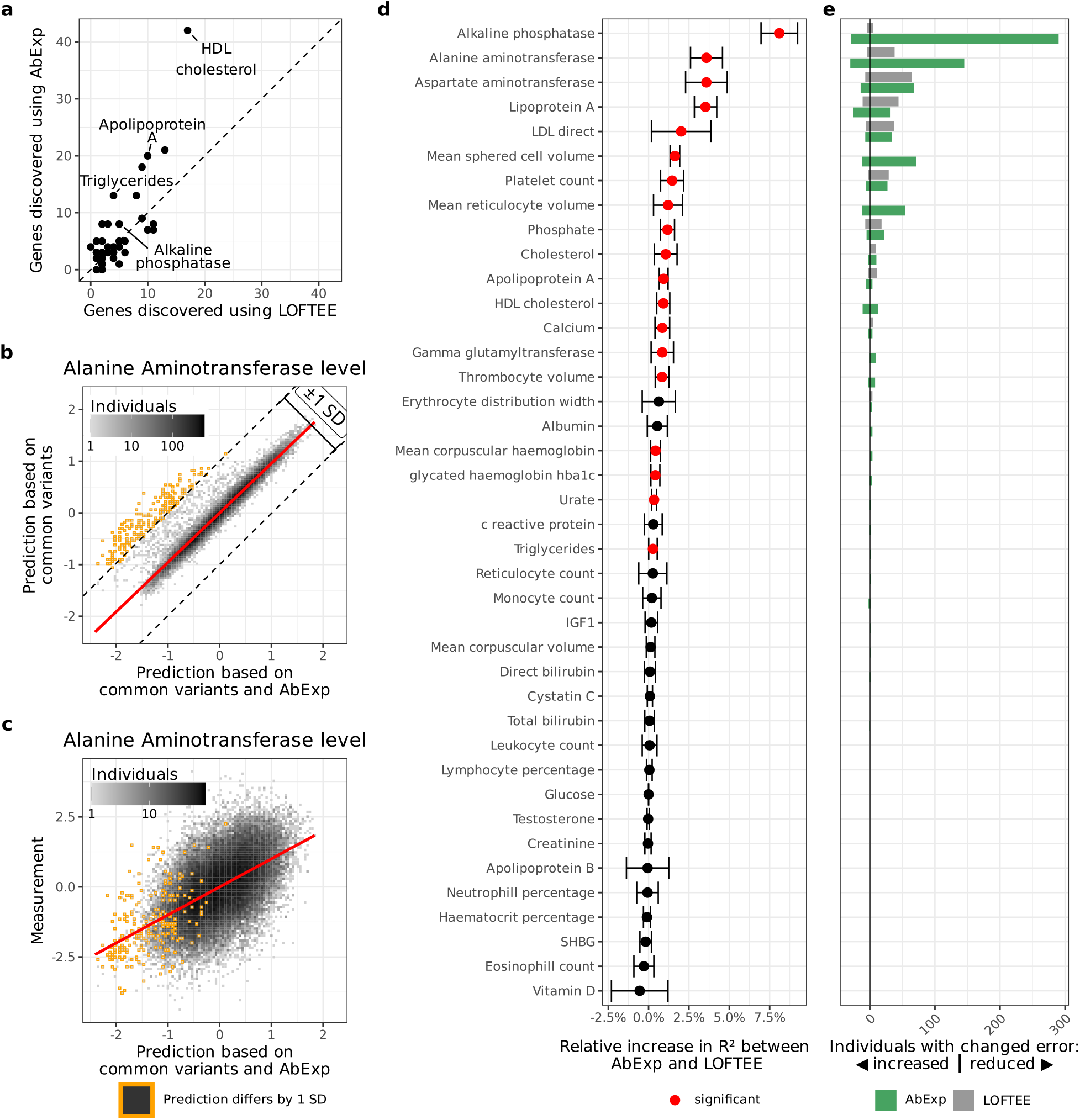
AbExp improves rare variant association testing and phenotype prediction. **(a)** Number of genes associating with different traits using a model based on LOFTEE or AbExp. **(b)** Alanine aminotransferase level predicted using a model solely based on common variants (y-axis) against predictions using a model based on common variants and AbExp scores (x-axis). Individuals whose predictions differed by more than 1 standard deviation of the population trait distribution are marked in orange. **(c)** Alanine aminotransferase measurements against predictions based on common variants and AbExp scores. Orange data points as in b). **(d)** Relative R^2^ increase between AbExp-based and LOFTEE-based predictions across traits. Traits with a significant difference between both models are marked red (two-sided paired t-test, nominal *P* < 0.05). Error bars show the standard deviation among 5 cross-validation folds. **(e)** Positive bars show the number of individuals with an error reduced by at least one standard deviation in the trait scale and therefore improved prediction, negative bars show the number of individuals with an error increased by at least one standard deviation in the trait scale and therefore worse prediction of the AbExp-based model (green) and the LOFTEE-based model (grey). All data presented in b-e) are computed on held-out folds of a 5-fold cross-validation within a third of the UKBB data not used for the gene discovery shown in a).

Having shown that AbExp can improve the gene-discovery sensitivity of RVAT, we next assessed its utility in phenotype prediction. To this end, we used the remaining third of the dataset which was not used for gene-trait association discovery. Specifically, we fitted gradient boosted trees models predicting the traits given the AbExp scores on the one hand or the number of LOFTEE variants on the other hand, of the genes discovered on the first two thirds of the data. Compared to linear regression models, gradient boosted trees can capture non-linear relationships through an ensemble of decision trees. These models were controlled for sex, age, the first 20 genetic principal components, and a polygenic risk score predicting the trait (Methods). The predictions rarely differed between the common variant-based model and the model further including AbExp scores of rare variants, as exemplified for the Alanine aminotransferase blood levels (Fig. 5b). For this trait, predictions for less than 0.5% of all individuals differed between the two models by more than 1 standard deviation of the population trait distribution (Fig. 5b). Remarkably, the trait values of those differing individuals tended to deviate largely from the population average, suggesting that the model integrating AbExp scores especially improves the predictions of individuals with extreme phenotypes that common variants cannot explain (Fig. 5c).

On held-out data, the phenotype prediction model based on AbExp scores significantly increased the amount of explained variation (R^2^) over the model based on LOFTEE in 45% of the traits and never significantly decreased R^2^ (Fig. 5d). Considering the number of individuals differing by more than 1 standard deviation to the common-variant based model, the AbExp based model improved the prediction of 816 individuals across the 40 blood traits, while the LOFTEE-based model only improved the prediction of 264 individuals (Fig. 5e). Moreover, the advantage of using AbExp scores were similarly observed when predicting phenotypes with regularized linear regression instead of gradient boosted trees (suppl. Fig. S17a,b), yet yielding less accurate phenotype predictions (suppl. Fig. S17c). These results using regularized linear regression, a method which like gradient boosted tree allows for robust fitting compared to standard linear regression but that does not model non-linearities, further confirm the added value of the tissue-specific predictions of AbExp for rare variant association testing.

Altogether, these results show that AbExp provides useful variant annotation for gene-trait association discovery by rare variant association testing and for building improved genetic risk scores.

### Incorporating RNA-seq from clinically accessible tissues (CATs) boosts prediction performance

RNA sequencing is becoming increasingly popular for rare disease diagnostics as a complementary assay to genome or exome sequencing as it allows the direct measurement of aberrant gene regulation^4–6,45–47^. However, many rare disorders are suspected to originate from tissues that can only be very invasively sampled such as the brain or the heart. We and others ^6,48^ have shown that clinically accessible tissues (CATs), notably skin fibroblasts and to a lesser extent whole blood, share a substantial fraction of expressed genes with non-CATs and, therefore, are likely to capture aberrant expression occurring in non-CATs. The GTEx dataset – a dataset of post-mortem samples – offers a unique opportunity to test the validity of this assumption as it provides matched samples for CAT and non-CAT tissues. We found that the mere ranking of genes according to their OUTRIDER *z*-score in fibroblast RNA-seq samples led to an average precision in predicting underexpression outliers in non-CATs of 20.1% (median across tissues), significantly larger than the genome-based predictor AbExp (10.1%, *P* = 1.6×10^− 6^, Fig. 6a). Next, we developed a model taking as input AbExp, whether the gene is expressed in a CAT and, if so, its OUTRIDER z-score. Using RNA-seq from skin fibroblasts to predict aberrant underexpression in all other tissues, this model reached an average precision of 23.8% in median across tissues (Fig. 6a). Consistent with previous work based on shared expressed genes^6,48^ and our work on aberrant splicing prediction^32^, fibroblasts turned out be more informative than whole blood (median average precision 9.3% using RNA-seq only and 17.5% when integrating AbExp, Fig. 6b), owing to blood expressing less genes than fibroblasts (See e.g. ref. ^6^). Altogether, we have established a method integrating direct measurements of aberrant expression from RNA-seq data in a CAT along with genomic variant annotations to predict aberrant underexpression in other tissues. Doing so, we showed that integrating RNA- seq from fibroblasts yields a substantial improvement as it doubles the average precision over using genomic variants alone.

**Figure 6:**
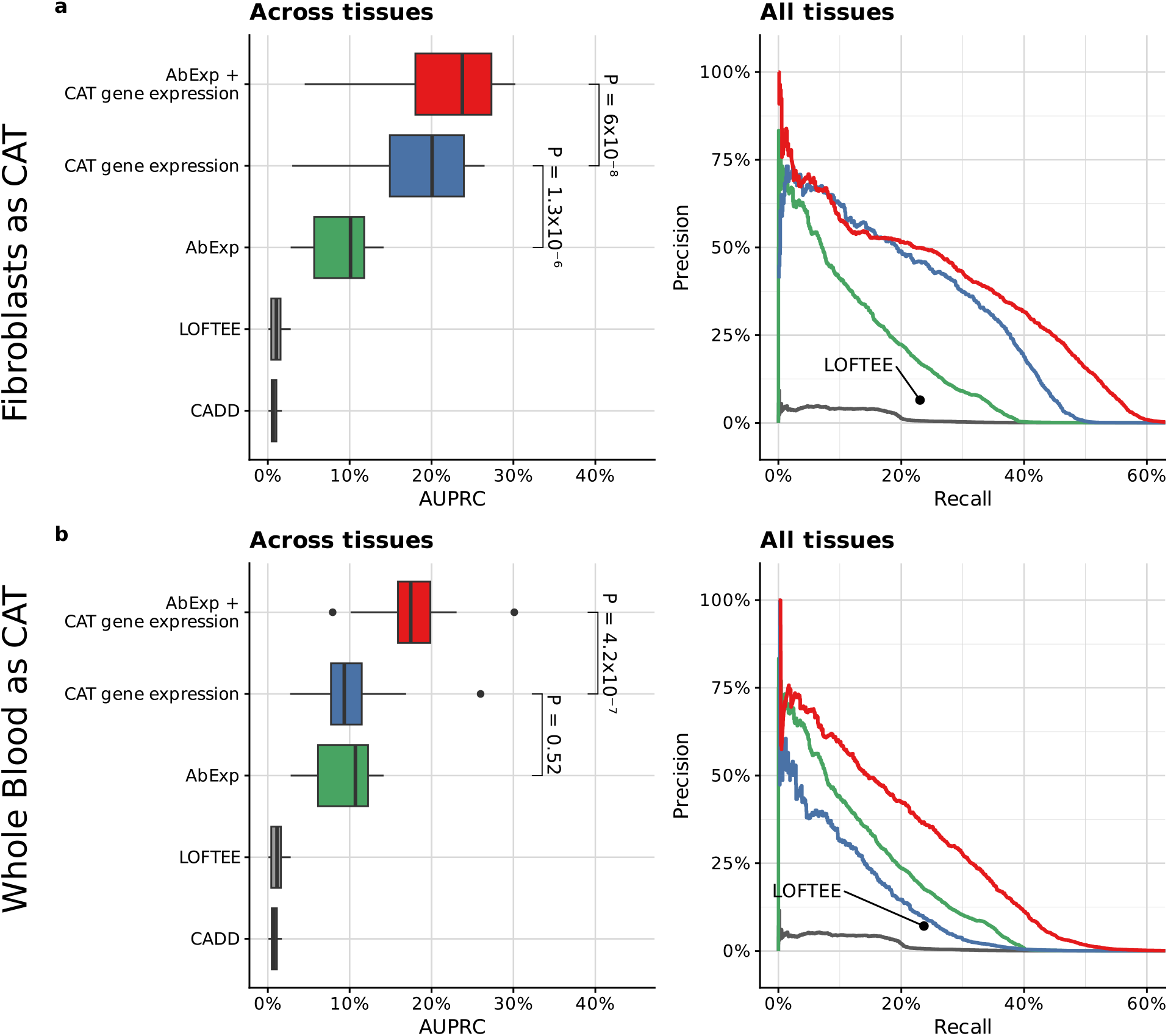
Combining RNA-seq measurements from clinically accessible tissues with AbExp further improves the prediction performance. **(a)** Left: Distribution per predictor (rows) of average precision (AUPRC) across 26 tissue types excluding skin tissues (Methods). Center line, median; box limits, first and third quartiles; whiskers span all data within 1.5 interquartile ranges of the lower and upper quartiles. *P*-values were obtained using the paired Wilcoxon test. The “Gene expression (CAT)” predictor ranks genes according to their OUTRIDER *z*-score in fibroblasts RNA-seq data. Right: Precision-recall curve aggregated across the same GTEx tissues as in the left panel. LOFTEE as a binary predictor is shown as a single point. **(b)** as in (a) using Whole blood as CAT and all other tissues as non-CAT.

## Discussion

Altogether, we established a benchmark dataset for aberrant gene underexpression prediction in 49 human tissues, addressing an unmet need in the area of high-impact variant effect prediction. We developed AbExp, a machine learning model predicting aberrant underexpression across tissues by integrating existing variant annotations with tissue-specific gene expression variability and transcript isoform composition. AbExp outperformed existing variant annotation tools by up to 7-fold in average precision. Enhanced enrichments for pathogenic variants and for variants not yet observed in gnomAD, particularly among mutationally constrained genes, indicate that AbExp is a promising tool for variant prioritization in clinical diagnostics. Using UK Biobank blood traits, we demonstrated that the continuous and tissue-specific AbExp scores provide added information over the state-of-the-art putative loss of function classifier LOFTEE for rare variant gene association testing as well as for phenotype prediction. Finally, we showed that AbExp scores can be combined with gene expression measurements from clinically accessible tissues to predict aberrant expression in other tissues yielding an increased prediction performance by 2-fold over AbExp.

Refining the predictor, while our primary objective, also shed light into the biology of underexpression outliers. We found that gene expression variability plays a dual role in this context. On the one hand, altering expression of a gene is more likely to result in an outlier if the expression of the gene varies little in the population than if it varies largely. On the other hand, each variant category was associated with milder fold-changes for genes with lower expression variability, indicating the involvement of regulatory mechanisms that confer robustness to genetic perturbations. We modeled the outcome of these two counteracting phenomena with a non-linear model trained from the data. Future biophysical investigations unraveling these buffering mechanisms could help improving the predictions and more generally improve variant interpretation. Our work also confirmed the importance of nonsense-mediated decay, which underpins a substantial proportion of the outliers, and the need to take tissue-specific transcript isoform into account when interpreting splice-affecting and nonsense variants as pioneered by Cummings and colleagues^24^.

This study has limitations. A basic assumption of AbExp is that an underexpression outlier is caused by a rare variant. However, it is possible that a rare combination of frequent genetic variants causes an expression outlier. Also, damage caused by one variant might be recovered by another variant, e.g. a second frameshift variant recovering the frame after a first frameshift variant. AbExp does not evaluate combinations of variants, which would require more complex modeling in particular by taking phasing into account. Moreover, AbExp covers the variants up to 5 kb away from transcript boundaries, missing middle-range and long-range enhancers. Integrating structural variants located further 5’ of the transcription start did not lead to significant performance improvement. Future work could expand to distal variants, for instance if sequence-based models of gene expression improved at this task^21^. Also, this work was focused on cis-acting regulation by considering only variants located within or near the genes. The effect of trans-acting gene regulation, which would need a very different modeling paradigm than investigating here in order to capture regulatory networks, remains to be addressed. The current recall remains modest. However, this can be an underestimation of the true recall since it is unclear how many of the outliers that we aim to predict are artifacts, due to mosaicism, non- genetic causes, or because they are measurement errors. Moreover, it is unclear whether a gene expression outlier is caused by a change of transcript length or transcript abundance. Indeed, our outlier classifications are based on read counts across annotated exons, which may include shorter transcripts among down-regulated outliers^6^. Given that GTEx data relies on short-read sequencing, accurately determining transcript length is challenging. Perhaps the advance of long-read RNA-sequencing will allow to disambiguate these cases in the future. Finally, we have used here an outlier calling method that was applied for each tissue separately and that could not leverage statistical evidence across tissues. Future aberrant expression predictors may be improved with better outlier calls.

Predicting gene expression from sequence is a long-standing goal of computational biology that is still far from completion. While existing sequence-based models of cis-regulation are trained across the whole range of expression levels^33,49,50^, we have proposed here to focus on extreme expression variations. Extremes may not well be captured by models trained to globally predict gene expression, as evident from the limited contribution of Enformer^33^ to our model. Also, the biological mechanisms underpinning extreme expression variations may differ from those governing moderate expression variations. However, the relevance of extreme expression for clinical diagnostics and research is high. We hope that the benchmark and algorithms we have developed will foster further research in this direction and aid in the development and validation of methods predicting the impact of large-effect variants on the human transcriptome.

## Methods

### Underexpression outlier benchmark dataset

#### GTEx dataset

We downloaded the GTEx RNA-seq read alignment files in the BAM format from dbGaP (phs000424.v8.p2). We excluded tissues with less than 60 RNA-seq samples due to insufficient statistical power^11^. This filter discarded bladder, endocervix, ectocervix, fallopian tube, and kidney medulla, leaving 49 tissues.

We obtained SNPs and small indels from the GTEx hg19 variant calls from the file GTEx_Analysis_2016-01- 15_v7_WholeGenomeSeq_635Ind_PASS_AB02_GQ20_HETX_MISS15_PLINKQC.vcf.gz from the dbGap entry phg000830.p1. Structural variants were obtained from Ferraro and colleagues^15^.

#### Expression outliers

Gene expression outlier analysis was performed following the aberrant expression module of DROP v1.1.0^51^ based on OUTRIDER^11^. To this end, we used as reference genome the GRCh38 primary assembly release 34 of the GENCODE project^52^. A fragment (reads pair) was assigned to a gene if and only if both reads were entirely aligned within the gene, allowing for fragments to be assigned to more than one gene. On each tissue separately, genes with an FPKM less than 1 in 95% or more of the samples were considered to be not sufficiently expressed in the tissue and filtered out, as previously described^11^. Since the variants were aligned to the hg19 reference genome, whereas the RNA-seq data was aligned to hg38, we lifted the gene expression counts by their Ensembl gene IDs over to hg19, relying on Gencode^52^ gene mappings. Overall, the liftover had a minimal impact with only 76 out of 19,682 genes and 294 out of 87,748 transcripts that could not be mapped.

OUTRIDER is an expression outlier caller that uses an autoencoder to model RNA-seq fragment count expectations with a negative binomial distribution. Specifically, OUTRIDER models the probability of the observed fragment count *x*_*s,g*_ for every gene *g* in a sample *s* as:

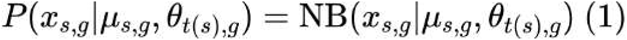

where:

- *µ*_*s,g*_ is the expected fragment count
- *θ*_*t(s),g*_ is the dispersion parameter for the gene *g* in the tissue of sample *s t(s)*

OUTRIDER further outputs:

- the biological coefficient of variation: 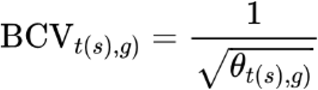
- the log_2_-transformed fold-change of the observed fragment count compared to the expected fragment count: 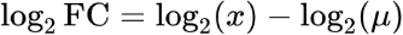
- the nominal *p*-value
- the False Discovery Rate using the Benjamini-Yekutieli method^53^

The resulting table was subsetted to individuals with an available whole genome sequencing and to protein-coding genes. Some of the data points could not be detected as outliers due to lack of statistical power. To reduce the proportion of these insufficiently powered data points in our benchmark, we discarded observations with an expected fragment count *µ*_*s,g*_ less than 450, a minimal value that was empirically estimated to allow recovering half of the two-fold reduction outliers transcriptome-wide upon a FDR cutoff of 5%^6^. We labeled as gene expression outliers all observations with an FDR less than 5%. Lastly, RNA-seq samples that contained more than 20 outliers were discarded because samples with numerous outliers may be samples for which OUTRIDER could not adequately fit the data or for which gene expression is globally affected, resulting in widespread expression aberrations throughout the genome that cannot be predicted from local sequence variation.

#### OUTRIDER *z*-score computation

We quantile-mapped the OUTRIDER-fitted negative binomial distributions (Eq. 1) to the standard normal distribution as follows:

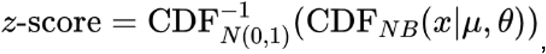

where 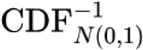 is the inverse cumulative distribution function of the standard normal distribution and CDF_*NB*_(*x*|*μ, θ*) the negative-binomial cumulative distribution function.

#### Precision-recall

We evaluated models using precision-recall curves due to the small proportion of outliers and summarized them with the area under the precision-recall curve (AUPRC) as implemented in ‘average_precision_score’ function of the scikit-learn package v1.3.2:

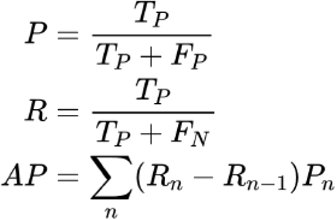

where P_n_ and R_n_ are the precision and recall at the n^th^ top prediction. The AUPRC is the average precision for each cutoff weighted by the recall difference.

The average precision was computed on all held-out data together on the one hand, and on held-out data for each of the 27 GTEx tissue types on the other hand. The GTEx tissue types group together highly similar tissues, notably many regions of the brain. By calculating performance within tissue types instead of reporting highly similar tissues separately, we avoid biasing our evaluation for overrepresented tissue types.

### Variant filtering, annotation, and gene-level aggregation

#### Filtering for rare high-quality variants

We considered a variant to be rare if it had a minor allele frequency in the general population ≤0.001 based on the Genome Aggregation Database (gnomAD v2.1.1) and was found in at most 2 individuals within GTEx. Variants had to be supported by at least 10 reads and had to pass the conservative genotype-quality filter of GQ > 30. For structural variants, we only filtered for the number of occurrences in the GTEx dataset (less than in 2 individuals).

#### Variant annotation with VEP and AbSplice

For all rare variants, we used Ensembl VEP^54^ v108 to calculate consequences, LOFTEE loss-of- function annotation, as well as CADD v1.6 scores. Tissue-specific aberrant splicing predictions were generated with AbSplice^32^.

#### Definition of gene-level features

For each combination of gene, individual, and tissue, we required a set of features to predict the underexpression outlier label. Therefore, we constructed a set of features starting from the annotations of the underlying rare variants. CADD predictions were max-aggregated per gene across the rare variants. For AbSplice, we kept the maximum absolute score in the corresponding gene and tissue.

Isoform-specific variant annotations, namely LOFTEE and VEP consequences, were aggregated differently depending on whether isoform proportions should be taken into account. When disregarding isoform proportions (i.e. the model “LOFTEE+CADD+consequences”), we only incorporated variants affecting the canonical transcript (canonical according to VEP). The gene was assumed to have an annotation (LOFTEE classification and each VEP consequence, e.g. stop-gained) when any of the variants in the gene had this annotation. Since the VEP canonical transcript does not differ between tissues, all tissues end up with the same set of gene-level features for these annotations.

For all models incorporating isoform proportions, isoform-specific variant annotations were first weighted by the total proportion of isoforms *i* in tissue *t* that they affect:

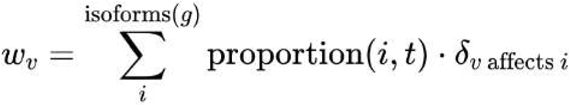

where *δ* _*υ affects i*_ is 1 if the variant affects the isoform, otherwise 0, the proportion of an isoform *i* in a tissue *t* was estimated as the median TPM proportion across individuals among all isoform of the same gene *g*:

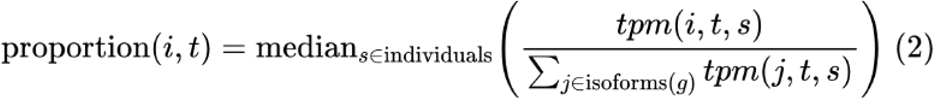

and where tpm(*i, t, s*) is the transcript-level TPM obtained from GTEx v8 (dbGaP Accession phs000424.v8.p2). All resulting variant annotations were then max-aggregated per gene and tissue across variants.

Enformer variant effect prediction

We used Enformer^33^ to evaluate the ability of deep-learning models to predict rare variant effects in promoters. Given a transcript, we first extracted three 393,216 nt DNA sequences, one centered at the transcription start site (TSS) and two others shifted by 43 nt upstream and downstream of the TSS. Subsequently, Enformer was applied to each sequence, and we extracted predictions for various subsets out of the 5,313 human assay tracks on the three central bins from each sequence. We denote *p* as the number of selected tracks. To ensure that regulatory elements falling on bin boundaries were accounted for, we averaged the predictions over the sequence shifts *k* and central bins *l*:

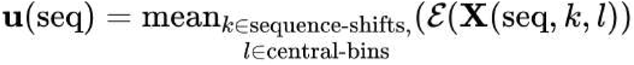

Here, the *p*-dimensional vector contains the aggregated Enformer scores over the central bins and shifts for all the *p* tracks, the matrix **X**(seq, *k, l*) is the one-hot encoding of the shifted sequence *k* centered on bin *l*, and ℰ is the Enformer model. Since Enformer does not directly predict RNA-seq coverage for GTEx tissues, we trained different linear regression models (e.g. ElasticNet, Ridge) taking the aggregated Enformer scores **u**(seq) defined above as input to predict the expression level for each GTEx tissue averaged across all GTEx individuals, as previously^21^.

Specifically, we introduced *y*_*i,t*_ ≔ log_10_(1 + mean_*s*∈individual_ (tpm(*i, t, s*))), where is the tissue, *i* is the transcript, tpm(*i, t, s*) is the transcript-level TPM from GTEx v8 of individual, and aimed at predicting *y*_*i,t*_ linearly from the aggregate scores. Hence, we modeled 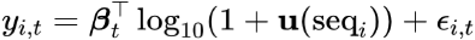, and estimated the coefficients with different regularized linear regression schemes (e.g. ElasticNet, Ridge).

Having fitted this model, we now have a function 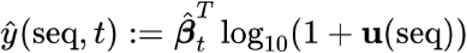 that predicts gene expression from sequence for each tissue. We employed two different methods to obtain a variant effect score for a given gene. First, we estimated the effect of the variant on the canonical transcript of the gene, if the variant was in close proximity to its promoter, as follows:

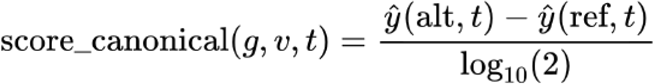

Here, *υ* is the variant of interest, *t* is the tissue, *g* is the gene of interest, and and are the alternative and reference sequences of the canonical transcript respectively. Second, we weighted all isoforms of the gene by their isoform proportions, using the following score:

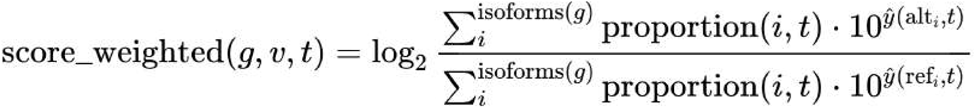

Here, υ is the variant of interest, *t* is the tissue, *g* is the gene of interest, alt_*i*_ and ref_*i*_ are the alternative and reference sequences of transcript *i* respectively, and isoforms (*g*) returns the isoforms of *g* that are located in close proximity to the variant.

For each GTEx individual, tissue, and gene combination, we took the signed maximum absolute variant effect across all relevant variants as the variant effect score. Finally, we performed cross-validation to optimize (1) the subset of Enformer tracks, (2) the linear model parameters (e.g. L1 and L2 regularization), (3) the variant-effect function (i.e. weighted or canonical isoforms), and (4) the shortest interval around the TSS, for which Enformer is predictive (suppl. Fig. S9).

### Variant type enrichment in underexpression outliers

To obtain the proportion of underexpression outliers explained by different classes of variants, we first identified the variant consequences each gene-individual-tissue combination is affected by. We then assigned each gene-individual-tissue combination to be caused by the most impactful consequence among the following consequences: Downstream variant, Inframe CDS (inframe deletion, inframe insertion, missense variant), Intron, NMD-like (NMD escaping, Frameshift, Start lost, Stop gained, Stop lost), Promoter (Upstream gene variant), Splicing disruption (Splice acceptor/donor/region variant), Structural ablation (Transcript ablation), Synonymous, UTR (3’ UTR, 5’ UTR). The order of assignment was determined by performing a Fisher’s exact test for each variant consequence against the underexpression outlier class and ordering the classes by decreasing significance. Afterward, the proportion of underexpression outliers explainable by each variant type was calculated (suppl. Fig. S3).

### Model training

All 633 individuals were split into six cross-validation groups with approximately equal numbers of underexpression outliers and tissues.

The DNA-based models were trained to predict the OUTRIDER *z*-scores. To this end, we used gradient-boosted trees^55^ from the LightGBM^56^ framework with default parameters: boosting_type: gbdt, learning_rate: 0.1, max_depth: -1, min_child_samples: 20, min_child_weight: 0.001, min_split_gain: 0, n_estimators: 100, num_leaves: 31, reg_alpha: 0, reg_lambda: 0, subsample: 1, subsample_for_bin: 200000, subsample_freq: 0.

For training the model integrating CAT RNA-seq data, the same cross-validation scheme was used as for the DNA-based models while excluding the CAT and highly related tissues thereof from the predicted tissues. When using fibroblasts as CAT, we excluded the non sun-exposed suprapubic skin, sun-exposed lower leg skin, and cultured fibroblasts. When using whole blood as CAT, we excluded whole blood and EBV-transformed lymphocytes from the predicted tissues. Unlike the DNA-based models, this model was trained to predict underexpressed outliers using a logistic regression taking as input i) a binary variable indicating whether the gene is expressed in the CAT, ii) the OUTRIDER *z*-score of the gene in the CAT, iii) the AbExp prediction for the target tissue, and all 3 interaction terms between those 3 variables.

To compare Watershed with other machine learning methods, we trained Watershed models using the following features: CADD, LOFTEE (high-confidence), LOFTEE (low-confidence), 5’ UTR variant, 3’ UTR variant, downstream gene variant, intron variant, missense variant, non- coding transcript exon variant, splice acceptor variant, splice donor variant, splice region variant, stop gained, synonymous variant, upstream gene variant, sift score, polyphen score. As the dataset was too large to be fitted in total with Watershed, we subsampled the non-outliers to ten times the number of outliers. This change in class balance affects the posterior by constant but not their ranking. Afterwards, we extracted the conditional probabilities P(Z|G) instead of the standard posterior of P(Z|G,E) to predict the likelihood of an expression outlier on the validation folds. Z is the latent variable, G is the vector of genomic annotations, and E is the outlier status.

### Replication in independent datasets

In the two replication datasets, variants were filtered for genotype quality ≥ 30 and read depth ≥ 10 reads. Moreover, rare variants were subsetted based on the gnomAD population with minor allele frequency ≤ 0.001. Those high-quality and rare variants were used as candidates for outlier prediction. Gene expression outliers were obtained with OUTRIDER and filtered for a sufficiently large expected number of fragments (*µ* > 450). Samples with more than 20 outliers were removed.

The mitochondrial disease dataset^6^ consisted of 311 whole-exome sequencing samples paired with RNA-seq from fibroblasts. After filtering, this dataset contained 808 underexpression outliers across 295 samples. A detailed overview of how many samples, genes, etc. remained after each filtering step can be seen in suppl. table T4.

For the amyotrophic lateral sclerosis (ALS) dataset, we downloaded 253 transcriptomes with matched whole-genome sequencing data from https://dataportal.answerals.org^35^. RNA-seq measurements were obtained from iPSC-derived spinal motor neurons. After filtering, the dataset contained 653 underexpression outliers across 233 samples (195 cases and 38 controls). The corresponding overview of how many samples, genes, etc. remained after each filtering step can be seen in suppl. table T5.

### Prediction of ClinVar pathogenic variants

To assess the efficacy of AbExp in distinguishing pathogenic from benign variants, we annotated known pathogenic, likely benign, and benign variants from the ClinVar database with both AbExp and CADD. Importantly, we used the official CADD pipeline to annotate all variants, including those with missing pre-computed scores. Variants classified as both likely benign and benign were retained solely as likely benign within the dataset. Furthermore, for each variant, we retained only the minimum predicted AbExp score across all tissues.

### GnomAD variant enrichments

For all possible SNVs within 5 kb of 19,018 protein-coding genes, we computed AbExp scores with and without using the BCV, CADD scores, and LOFTEE high-impact loss of function annotations. High-impact predictions of CADD and AbExp without BCV were identified by matching score cutoffs to the quantile of the high-impact cutoff of AbExp. Using logistic regression fits, we then computed the odds-ratio of high-impact variants among absent, singleton, rare (MAF < 0.1%), and common SNVs in gnomAD for these models. Moreover, we calculated odds-ratios for these variant types and methods as a function of gene LOEUF decile. The loss-of-function observed/expected upper bound fraction (LOEUF) scores were downloaded from https://gnomad.broadinstitute.org/downloads.

### UK Biobank rare variant association testing and phenotype prediction

We analyzed data from 200,593 European ancestry unrelated individuals in the UK Biobank (using field 22011 and labeled as Caucasian in field 22006), all of whom had genotypes available from exome-sequencing and microarrays as well as blood and urine measurements. A detailed list of used phenotypes can be found in suppl. table T2. For every trait, trait values were inverse rank normal transformed, as in the Genebass study^38^.

#### Identification of lead trait-associated variants

To control for common variants in the vicinity of each gene, trait-associated variants were obtained from the PanUKBB study^57^. Using plink v1.9^58,59^, variants with a p-value ≤ 0.0001 were clumped in a 250-kb window with an LD-cutoff of r^2^ < 0.5 to identify independent lead variants for every trait. The imputed genotypes were then subsetted for these lead variants in a 250-kb window around each gene.

#### Application of polygenic risk scores

Polygenic risk scores were selected from the study by Privé et al.^60^ if available and otherwise from the study by Tanigawa et al.^61^ (suppl. Table T2). Score files were obtained from the PGS catalog database^62^ and applied to the imputed genotypes using plink v2.0^58,63^.

#### Calculation of AbExp and LOFTEE scores

The UKBB whole-exome sequencing data was subsetted for variants with a minor allele frequency ≤0.001 based on gnomAD v3.1.1 and filtered for genotype quality ≥ 30 and read depth ≥ 10 reads. The remaining variants were then annotated using Ensembl VEP^54^ v108 with the LOFTEE plugin^22^ and AbExp.

#### Rare variant association testing

Gene association was tested using a likelihood ratio test between a restricted linear regression model containing only covariates and four linear regression models with additional predictor variables as described in Table 1.

**Table 1:** Rare variant association test models.

| Model | Predictor variables | Nr. of variables |
| --- | --- | --- |
| LOFTEE | Number of LOFTEE variants in the gene of interest | 1 |
| AbExp all tissues | The set of minimal AbExp scores across all rare variants in the gene (0 in absence of any rare variant) for each tissue | 49 |
| Minimum AbExp | The minimum AbExp score across all rare variants in the gene and across all tissues, 0 in absence of any rare variant | 1 |
| Median AbExp | The median AbExp score across all rare variants in the gene and across all tissues, 0 in absence of any rare variant | 1 |

The following covariates were used for all models: sex, age, age^2^, age times sex, age^2^ times sex, the 20 first genetic principal components (field 22009), lead associated variants in 250-kb window around the gene of interest, and a polygenic risk score predicting the trait. The polygenic risk scores and lead variants were based on the whole UK Biobank dataset. This mild data leakage may have led to model overfitting. However, since these features were used as covariates in both restricted and full models, the comparison between models remained valid. P- value calibration of the models was assessed by permuting the phenotype once. Identification of significantly trait-associated genes was performed on two thirds of the dataset.

#### Phenotype prediction

For the common-variant-based phenotype prediction model, the following features were used: sex, age, age^2^, age times sex, age^2^ times sex, the 20 first genetic principal components (field 22009), and a polygenic risk score predicting the trait. In contrast to the rare variant association testing, we did not include lead variants of the associating genes, as these would lead to a too large number of predictor variables.

We compared two phenotype prediction models integrating rare and common variants. Both are nonlinear models. To this end, we used gradient-boosted trees^55^ from the LightGBM^56^ framework with default parameters: boosting_type: gbdt, learning_rate: 0.1, max_depth: -1, min_child_samples: 20, min_child_weight: 0.001, min_split_gain: 0, n_estimators: 100, num_leaves: 31, reg_alpha: 0, reg_lambda: 0, subsample: 1, subsample_for_bin: 200000, subsample_freq: 0. Both models include the same features as the common-variant based prediction models. The AbExp phenotype prediction model further included the AbExp scores of significantly trait-associated genes in 49 tissues, while the LOFTEE phenotype prediction model included the number of LOFTEE pLoF variants of significantly trait-associated genes. For each trait, the two models can have a different set of significantly trait-associated genes as identified by their corresponding RVATs. The phenotype prediction models were trained with 5-fold cross- validation on the remaining third of the dataset. All evaluations were performed on the hold-out folds.

## Declarations

### Ethics approval and consent to participate

No new data was generated for this study. The different ethics approvals can be found in the corresponding publications (GTEx^64^, AnswerALS^35^, mitochondrial disease^6^). The UK Biobank was approved by the North West Multi-centre Research Ethics Committee (21/NW/0157). Our reference number approved by the UK Biobank is 25214. All UK Biobank study participants gave written informed consent.

The research conformed to the principles of the Declaration of Helsinki.

### Consent for publication

All individuals included or their legal guardians provided written consent to share pseudonymized patient data and analysis data, as described in the original publications.

## Acknowledgements

We thank Felix Brechtmann, Alexander Karollus, and Holger Prokisch for fruitful discussions and their valuable feedback on the manuscript. Figure 1 was created with BioRender.com. We thank Žiga Avsec and Alexander Karollus for advice on using the Enformer in this context.

## Authors’ contributions

J.G. conceptualized the project. F.R.H., F.P.C. and J.G. designed the methodology. F.R.H., N.W. and V.A.Y. curated the data. F.R.H., J.L., and G.T. performed investigation and developed the software. F.R.H., J.L., G.T., and V.A.Y. performed visualizations. F.R.H., V.A.Y. and J.G. wrote the original draft of the manuscript. All authors reviewed and edited the manuscript. J.G. supervised the project with the help of F.P.C. and V.A.Y.

## Availability of data and materials

## Data availability

No primary data were generated for this study. Rare variants from gnomAD v.2.1.1 are publicly available at https://gnomad.broadinstitute.org. The GTEx v8 dataset is available at (under dbGaP protection) https://gtexportal.org/home. The ALS dataset is available at http://dataportal.answerals.org after a registration and approval process. The mitochondrial dataset is described by Yépez et al.^6^ The UK Biobank dataset is available at https://biobank.ndph.ox.ac.uk/ukb/ after a registration and approval process. We provide the aberrant expression benchmark dataset, isoform proportions, and the expected gene expression in GTEx v8 as open-access in the Zenodo repository^65^ (doi: 10.5281/zenodo.13895620).

## Code availability

A Snakemake pipeline to calculate AbExp predictions can be found at https://github.com/gagneurlab/abexp.

The source code of Watershed adapted for underexpression prediction can be found at https://github.com/Hoeze/Watershed.

The source code for the UK Biobank rare-variant association study and phenotype prediction can be found in the following repositories:

- Main analysis pipeline: https://github.com/gagneurlab/abexp-ukbb-trait-analysis
- Variant clumping: https://github.com/gagneurlab/abexp-ukbb-variant-clumping
- Polygenic risk score calculation: https://github.com/gagneurlab/abexp-ukbb-prs

## References

1. Wang, L.-H., Wu, C.-F., Rajasekaran, N. & Shin, Y. K. Loss of Tumor Suppressor Gene Function in Human Cancer: An Overview. Cell. Physiol. Biochem. 51, 2647–2693 (2018).

2. Kontomanolis, E. N. et al. Role of Oncogenes and Tumor-suppressor Genes in Carcinogenesis: A Review. Anticancer Res. 40, 6009–6015 (2020).

3. Gonorazky, H. D. et al. Expanding the Boundaries of RNA Sequencing as a Diagnostic Tool for Rare Mendelian Disease. Am. J. Hum. Genet. 104, 466–483 (2019).

4. Frésard, L. et al. Identification of rare-disease genes using blood transcriptome sequencing and large control cohorts. Nat. Med. 25, 911–919 (2019).

5. Murdock, D. R. Enhancing Diagnosis Through RNA Sequencing. Clin. Lab. Med. 40, 113–119 (2020).

6. Yépez, V. A. et al. Clinical implementation of RNA sequencing for Mendelian disease diagnostics. Genome Med. 14, 38 (2022).

7. Dekker, J. et al. Web-accessible application for identifying pathogenic transcripts with RNA-seq: Increased sensitivity in diagnosis of neurodevelopmental disorders. Am. J. Hum. Genet. 110, 251–272 (2023).

8. Deshwar, A. R. et al. Trio RNA sequencing in a cohort of medically complex children. Am. J. Hum. Genet. 110, 895–900 (2023).

9. Smail, C. et al. Integration of rare expression outlier-associated variants improves polygenic risk prediction. Am. J. Hum. Genet. 109, 1055–1064 (2022).

10. Li, B. et al. Evaluation of PrediXcan for prioritizing GWAS associations and predicting gene expression. in Biocomputing 2018 448–459 (WORLD SCIENTIFIC, 2017). doi:10.1142/9789813235533_0041.

11. Brechtmann, F. et al. OUTRIDER: A Statistical Method for Detecting Aberrantly Expressed Genes in RNA Sequencing Data. Am. J. Hum. Genet. (2018) doi:10.1016/j.ajhg.2018.10.025.

12. Salkovic, E., Sadeghi, M. A., Baggag, A., Salem, A. G. R. & Bensmail, H. OutSingle: a novel method of detecting and injecting outliers in RNA-Seq count data using the optimal hard threshold for singular values. Bioinformatics 39, btad142 (2023).

13. Segers, A., Gilis, J., Heetvelde, M. V., Baere, E. D. & Clement, L. Juggling offsets unlocks RNA-seq tools for fast scalable differential usage, aberrant splicing and expression analyses. 2023.06.29.547014 Preprint at 10.1101/2023.06.29.547014 (2023).

14. Li, X. et al. The impact of rare variation on gene expression across tissues. Nature 550, 239–243 (2017).

15. Ferraro, N. M. et al. Transcriptomic signatures across human tissues identify functional rare genetic variation. Science 369, (2020).

16. Hombach, S. & Kretz, M. Non-coding RNAs: Classification, Biology and Functioning. Adv. Exp. Med. Biol. 937, 3–17 (2016).

17. Zhang, Z., Qin, Y.-W., Brewer, G. & Jing, Q. MicroRNA degradation and turnover: regulating the regulators. Wiley Interdiscip. Rev. RNA 3, 593–600 (2012).

18. Megel, C. et al. Surveillance and Cleavage of Eukaryotic tRNAs. Int. J. Mol. Sci. 16, 1873–1893 (2015).

19. Nair, L., Chung, H. & Basu, U. Regulation of long non-coding RNAs and genome dynamics by the RNA surveillance machinery. Nat. Rev. Mol. Cell Biol. 21, 123–136 (2020).

20. GTEx Consortium. The GTEx Consortium atlas of genetic regulatory effects across human tissues. Science 369, 1318–1330 (2020).

21. Karollus, A., Mauermeier, T. & Gagneur, J. Current sequence-based models capture gene expression determinants in promoters but mostly ignore distal enhancers. Genome Biol. 24, 56 (2023).

22. Karczewski, K. J. et al. The mutational constraint spectrum quantified from variation in 141,456 humans. Nature 581, 434–443 (2020).

23. Rentzsch, P., Witten, D., Cooper, G. M., Shendure, J. & Kircher, M. CADD: predicting the deleteriousness of variants throughout the human genome. Nucleic Acids Res. 47, D886– D894 (2019).

24. Cummings, B. B. et al. Transcript expression-aware annotation improves rare variant interpretation. Nature 581, 452–458 (2020).

25. Morales, J. et al. A joint NCBI and EMBL-EBI transcript set for clinical genomics and research. Nature 604, 310–315 (2022).

26. Dugan, S. L. et al. New recessive truncating mutation in LTBP3 in a family with oligodontia, short stature, and mitral valve prolapse. Am. J. Med. Genet. A. 167, 1396–1399 (2015).

27. Sharon, D., Kimchi, A. & Rivolta, C. OR2W3 sequence variants are unlikely to cause inherited retinal diseases. Ophthalmic Genet. 37, 366–368 (2016).

28. Robinson, M. D., McCarthy, D. J. & Smyth, G. K. edgeR: a Bioconductor package for differential expression analysis of digital gene expression data. Bioinformatics 26, 139–140 (2010).

29. Bader, D. M. et al. Negative feedback buffers effects of regulatory variants. Mol. Syst. Biol. 11, 785 (2015).

30. Einarsson, H. et al. Promoter sequence and architecture determine expression variability and confer robustness to genetic variants. eLife 11, e80943 (2022).

31. Teran, N. A. et al. Nonsense-mediated decay is highly stable across individuals and tissues. bioRxiv 2021.02.03.429654 (2021) doi:10.1101/2021.02.03.429654.

32. Wagner, N. et al. Aberrant splicing prediction across human tissues. Nat. Genet. 55, 861–870 (2023).

33. Avsec, Ž. et al. Effective gene expression prediction from sequence by integrating long-range interactions. Nat. Methods 18, 1196–1203 (2021).

34. Haberle, V. & Stark, A. Eukaryotic core promoters and the functional basis of transcription initiation. Nat. Rev. Mol. Cell Biol. 19, 621–637 (2018).

35. Baxi, E. G. et al. Answer ALS, a large-scale resource for sporadic and familial ALS combining clinical and multi-omics data from induced pluripotent cell lines. Nat. Neurosci. 25, 226–237 (2022).

36. Fair, B. J. et al. Gene expression variability in human and chimpanzee populations share common determinants. eLife 9, e59929 (2020).

37. Wang, Q. et al. Rare variant contribution to human disease in 281,104 UK Biobank exomes. Nature 597, 527–532 (2021).

38. Karczewski, K. J. et al. Systematic single-variant and gene-based association testing of thousands of phenotypes in 394,841 UK Biobank exomes. Cell Genomics 2, 100168 (2022).

39. Fiziev, P. P. et al. Rare penetrant mutations confer severe risk of common diseases. Science 380, eabo1131 (2023).

40. Wu, M. C. et al. Rare-Variant Association Testing for Sequencing Data with the Sequence Kernel Association Test. Am. J. Hum. Genet. 89, 82–93 (2011).

41. Lee, S., Abecasis, G. R., Boehnke, M. & Lin, X. Rare-Variant Association Analysis: Study Designs and Statistical Tests. Am. J. Hum. Genet. 95, 5–23 (2014).

42. Sudlow, C. et al. UK Biobank: An Open Access Resource for Identifying the Causes of a Wide Range of Complex Diseases of Middle and Old Age. PLOS Med. 12, e1001779 (2015).

43. Zhou, W. et al. SAIGE-GENE+ improves the efficiency and accuracy of set-based rare variant association tests. Nat. Genet. 54, 1466–1469 (2022).

44. Clarke, B. et al. Integration of variant annotations using deep set networks boosts rare variant association genetics. 2023.07.12.548506 Preprint at 10.1101/2023.07.12.548506 (2023).

45. Cummings, B. B. et al. Improving genetic diagnosis in Mendelian disease with transcriptome sequencing. Sci. Transl. Med. (2017) doi:10.1126/scitranslmed.aal5209.

46. Kremer, L. S. et al. Genetic diagnosis of Mendelian disorders via RNA sequencing. Nat. Commun. 8, (2017).

47. Lunke, S. et al. Integrated multi-omics for rapid rare disease diagnosis on a national scale. Nat. Med. 29, 1681–1691 (2023).

48. Aicher, J. K., Jewell, P., Vaquero-Garcia, J., Barash, Y. & Bhoj, E. J. Mapping RNA splicing variations in clinically-accessible and non-accessible tissues to facilitate Mendelian disease diagnosis using RNA-seq. Genet. Med. Off. J. Am. Coll. Med. Genet. 22, 1181–1190 (2020).

49. Zhou, J. et al. Deep learning sequence-based ab initio prediction of variant effects on expression and disease risk. Nat. Genet. 50, 1171–1179 (2018).

50. Linder, J., Srivastava, D., Yuan, H., Agarwal, V. & Kelley, D. R. Predicting RNA-seq coverage from DNA sequence as a unifying model of gene regulation. 2023.08.30.555582 Preprint at 10.1101/2023.08.30.555582 (2023).

51. Yépez, V. A. et al. Detection of aberrant gene expression events in RNA sequencing data. Nat. Protoc. 16, 1276–1296 (2021).

52. Frankish, A. et al. GENCODE reference annotation for the human and mouse genomes. Nucleic Acids Res. 47, D766–D773 (2019).

53. Benjamini, Y. & Yekutieli, D. The control of the false discovery rate in multiple testing under dependency. Ann. Stat. 29, 1165–1188 (2001).

54. McLaren, W. et al. The Ensembl Variant Effect Predictor. Genome Biol. 17, 122 (2016).

55. Hastie, T., Tibshirani, R. & Friedman, J. Boosting and Additive Trees. in The Elements of Statistical Learning: Data Mining, Inference, and Prediction (eds. Hastie, T., Tibshirani, R. & Friedman, J.) 337–387 (Springer, New York, NY, 2009). doi:10.1007/978-0-387-84858-7_10.

56. Ke, G. et al. LightGBM: A Highly Efficient Gradient Boosting Decision Tree. in Advances in Neural Information Processing Systems vol. 30 (Curran Associates, Inc., 2017).

57. Pan-UKB team. Pan-ancestry genetic analysis of the UK Biobank. (2020).

58. Chang, C. C. et al. Second-generation PLINK: rising to the challenge of larger and richer datasets. GigaScience 4, s13742-015-0047–8 (2015).

59. Shaun Purcell & Christopher Chang. PLINK 1.9. (2020).

60. Privé, F. et al. Portability of 245 polygenic scores when derived from the UK Biobank and applied to 9 ancestry groups from the same cohort. Am. J. Hum. Genet. 109, 12–23 (2022).

61. Tanigawa, Y. et al. Significant sparse polygenic risk scores across 813 traits in UK Biobank. PLOS Genet. 18, e1010105 (2022).

62. Lambert, S. A. et al. The Polygenic Score Catalog as an open database for reproducibility and systematic evaluation. Nat. Genet. 53, 420–425 (2021).

63. Shaun Purcell & Christopher Chang. PLINK 2.0. (2022).

64. GTEx Consortium. The Genotype-Tissue Expression (GTEx) project. Nat. Genet. 45, 580–585 (2013).

65. Hölzlwimmer, F. R. Aberrant gene expression prediction benchmark based on GTEx v8. Zenodo 10.5281/zenodo.13895620 (2024).

