## Supplementary figures for "Aberrant expression prediction across human tissues"

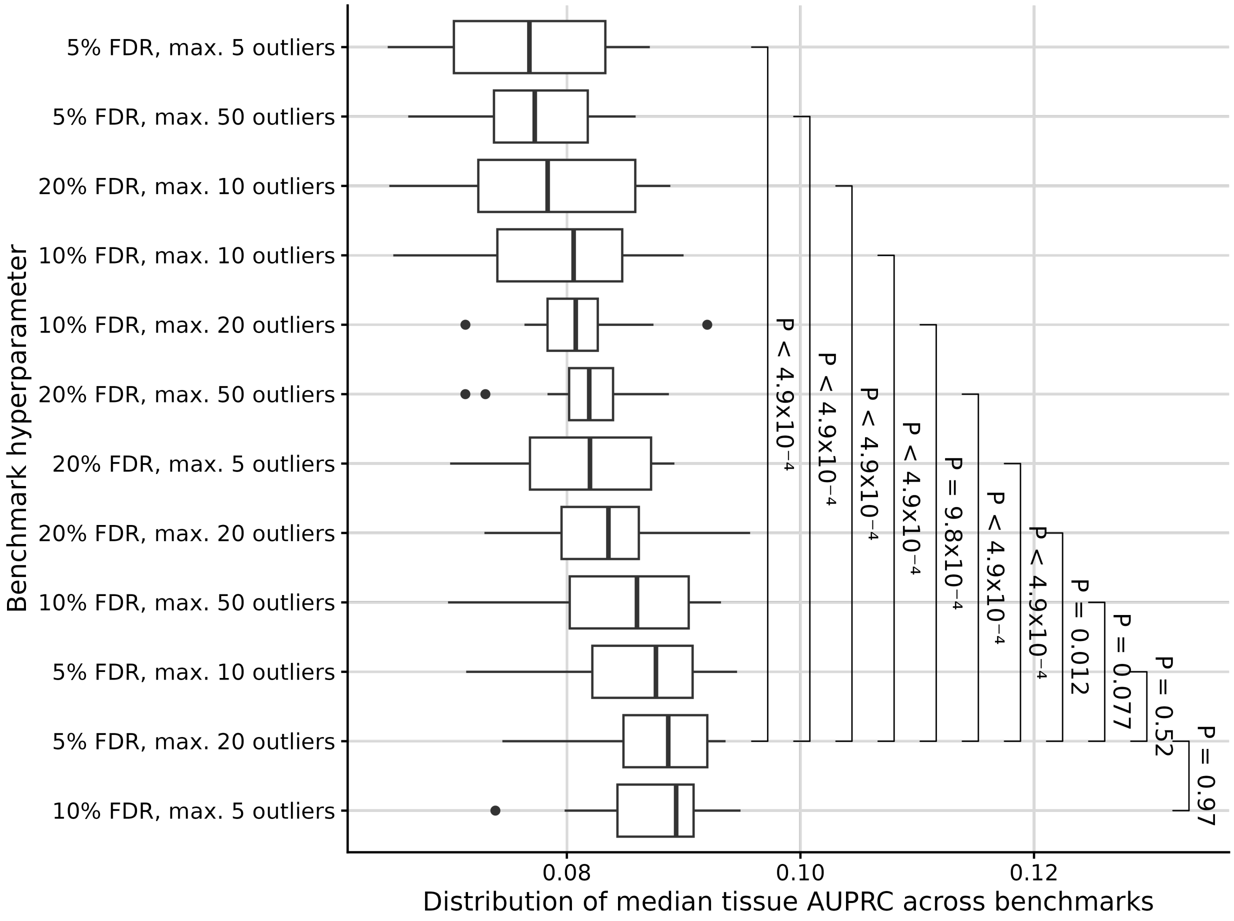


**Supplementary Fig. S1: OUTRIDER cutoff selection for the definition of the benchmark dataset.** We evaluated various FDR thresholds (0.05, 0.1, and 0.2) and cutoffs on the number of outliers per sample (5, 10, 20, 50), both for training and evaluation. To this end, we trained simple models using previously established features (LOFTEE, CADD, VEP, and structural variants) to predict underexpression outliers with a LightGBM regression on the gene expression z-score (methods), ensuring that the cross-validation folds contain the same individuals in all settings. Performance was evaluated as the median area under the precision-recall curve across tissues under all these settings (boxplots, n=12 benchmark settings). Box label, sample size; Center line, median; box limits, first and third quartiles; whiskers span all data within 1.5 interquartile ranges of the lower and upper quartiles. The settings of the bottom two boxplots led to quasi-uniformly better models across all 12 benchmark settings used for evaluation. Among these two settings, we opted for 5% FDR and max. 20 outliers as it provided twice as many outliers for training.

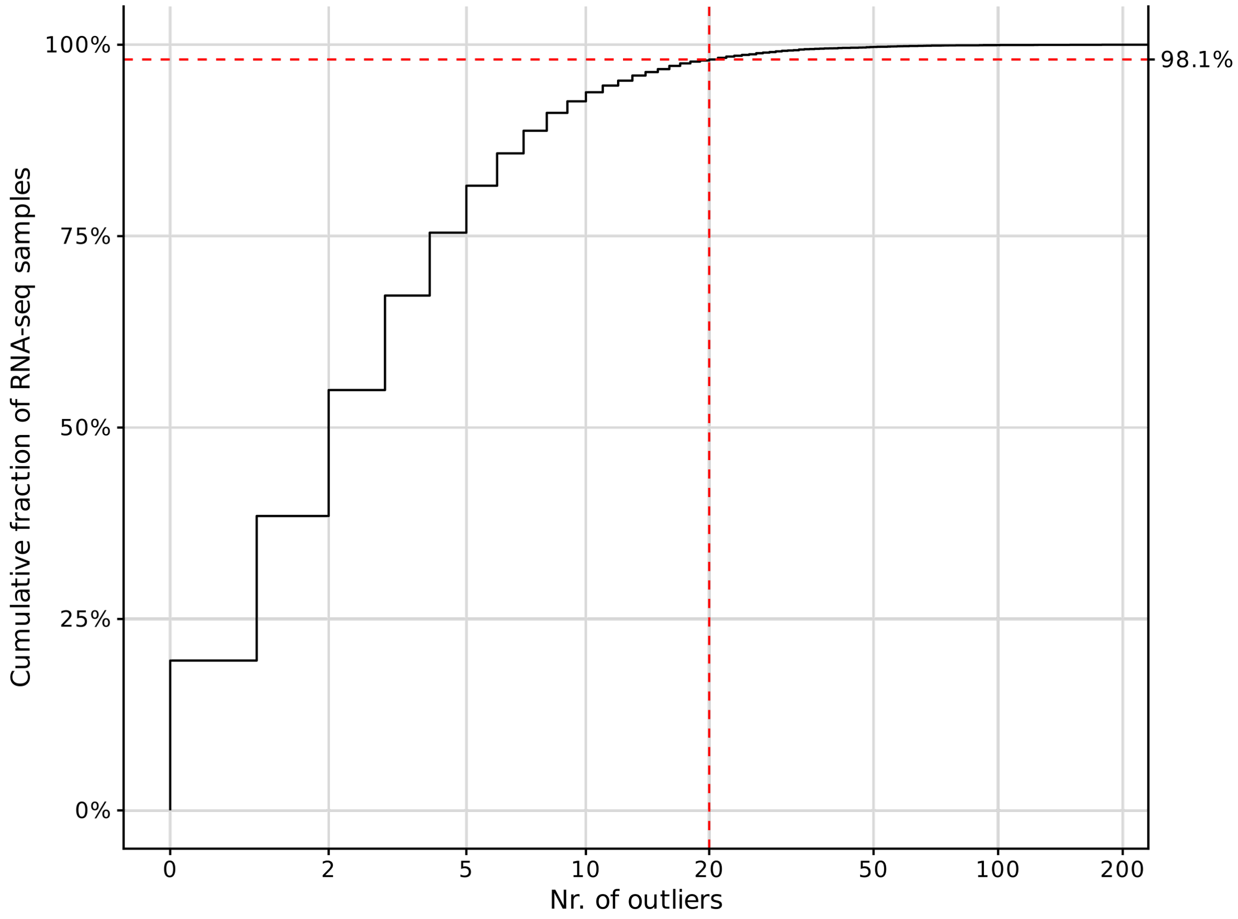


**Supplementary Fig. S2: Only 1.9% of GTEx transcriptome sequencing samples have more than 20 outliers.** Cumulative fraction of RNA-seq samples (y-axis) that have at most a given number of outliers (x-axis) in the GTEx dataset. The vertical dashed line denotes the 20 outlier cutoff, which 98.1% of samples passed (horizontal dashed line).

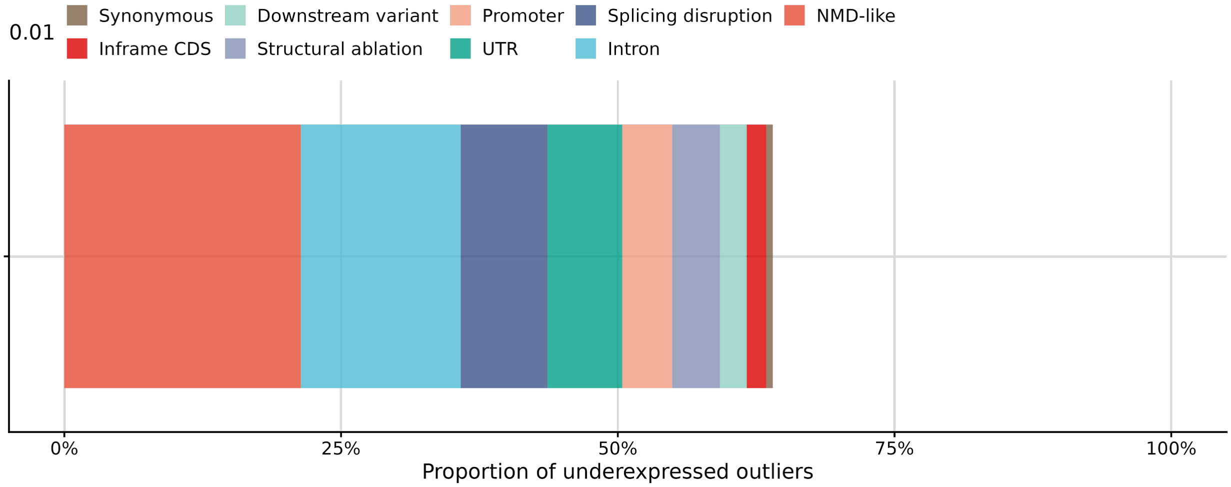


**Supplementary Fig. S3: Proportion of underexpression outliers explained by different classes of variants.** Each outlier is assigned to the most impactful consequence and therefore occurs only in one category (Methods).

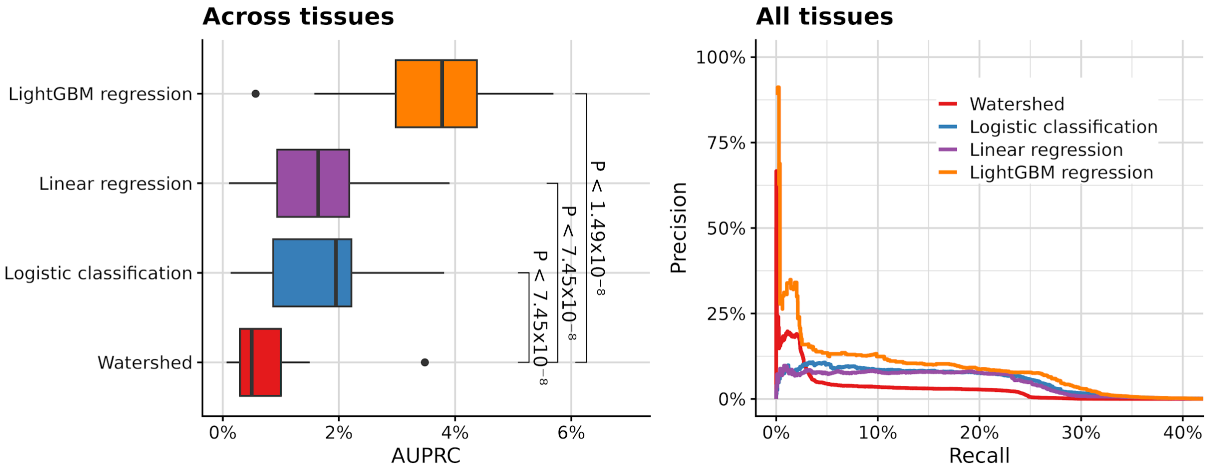


**Supplementary Fig. S4: Watershed model architecture is outperformed by classic machine learning methods.** **(a)** Distribution of average precision (AUPRC) across 27 GTEx tissue types. P-values were obtained using the paired Wilcoxon test. Box label, sample size; Center line, median; box limits, first and third quartiles; whiskers span all data within 1.5 interquartile ranges of the lower and upper quartiles. (**b**) Precision-recall curve for all tissues combined.

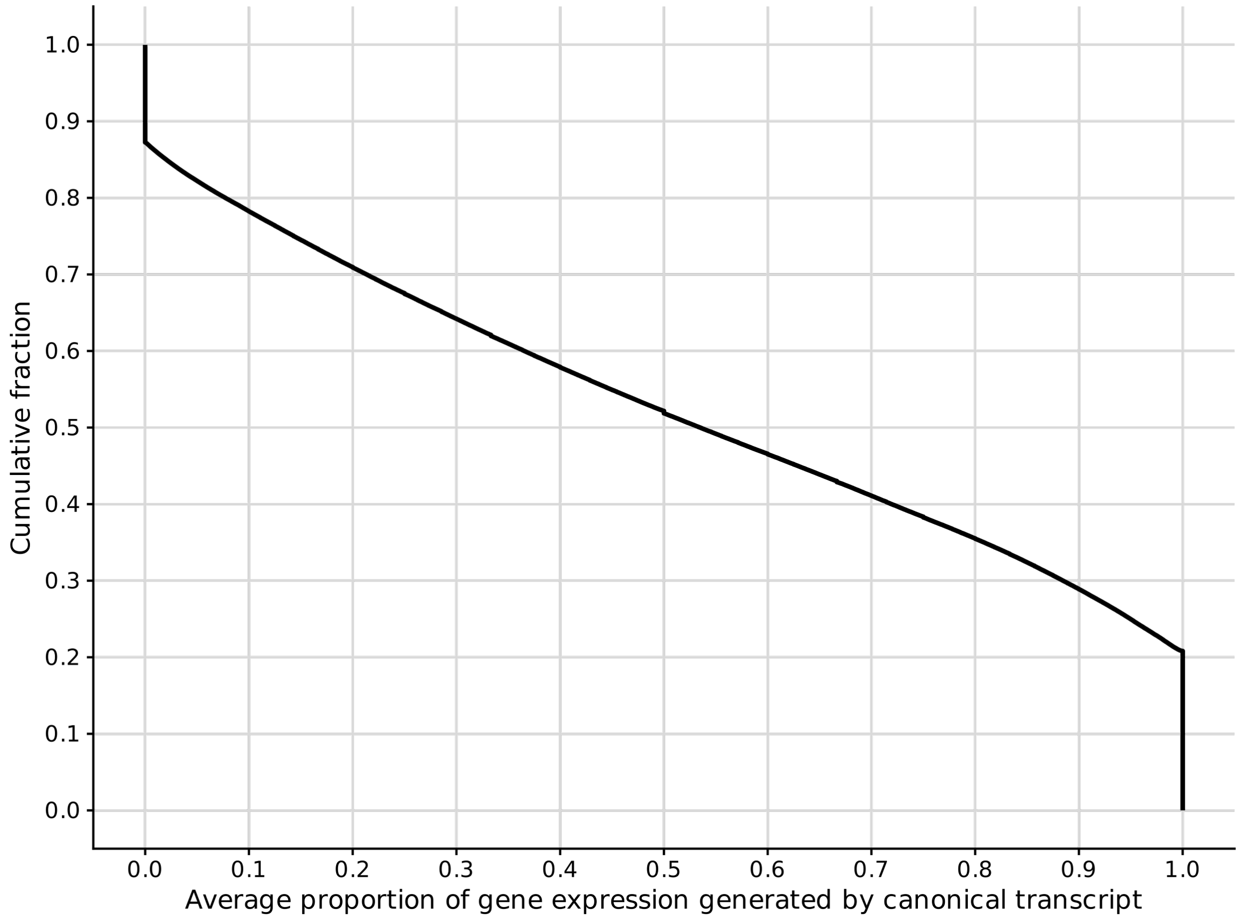


**Supplementary Fig. S5**: **Cumulative fraction of genes (y-axis) for which the canonical transcript contributes to more than a given proportion (x-axis) to the total gene expression.** The data is aggregated over all tissues and restricted to the expressed genes per tissue. Canonical according to VEP.

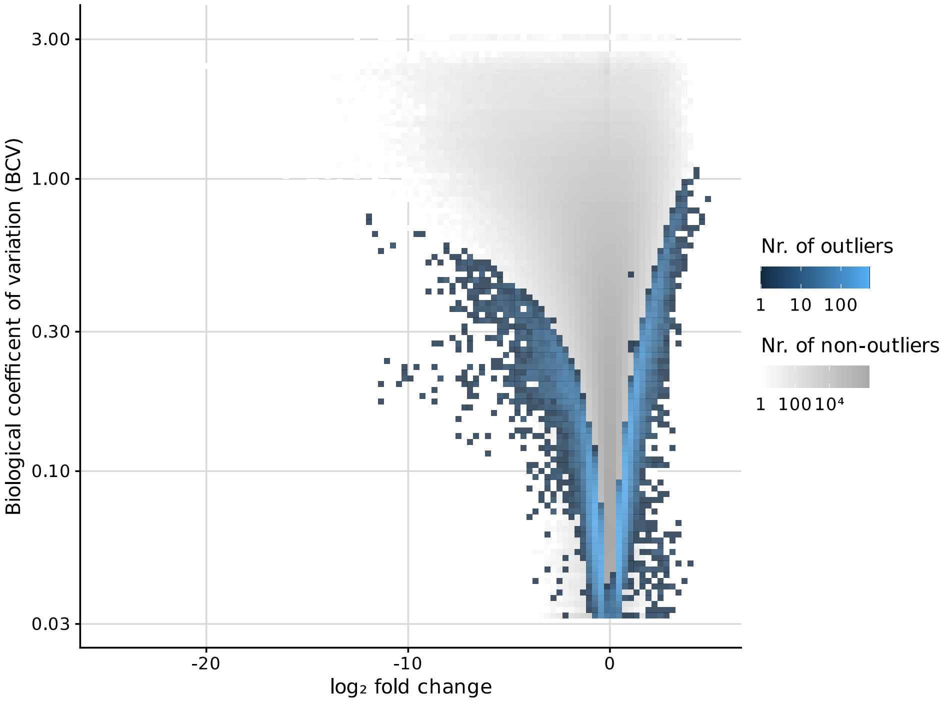


**Supplementary Fig. S6**: **Biological coefficient of variation (BCV) against expression fold change across all genes and tissues.** The BCV is an estimate of the coefficient of variation of expression of a gene measured by RNA-seq read counts controlling for sampling noise. The larger the BCV of a gene in a tissue, the more variable it is across the GTEx population. BCV is used instead of standard deviation to account for sampling noise, which particularly affects low RNA-seq read counts (Methods). Highly variable genes require a larger fold change to be called an outlier.

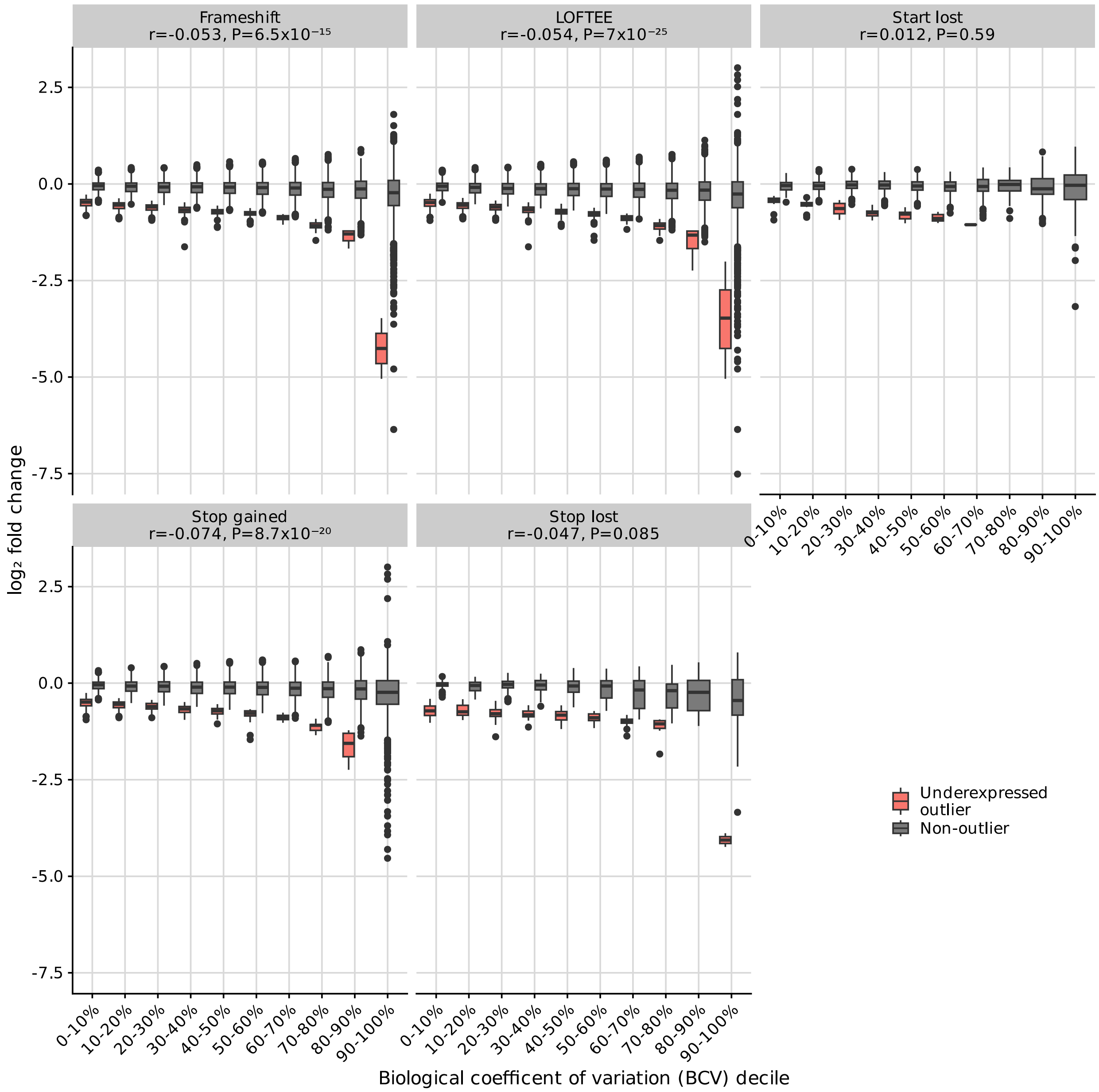


**Supplementary Fig. S7**: **Distribution of gene expression fold changes among genes in different deciles of expression variability, given that the gene is affected by some rare variant consequence (e.g. frameshift, LOFTEE).** Center line, median; box limits, first and third quartiles; whiskers span all data within 1.5 interquartile ranges of the lower and upper quartiles. The higher the coefficient of variation, the larger the gene expression impact of the variants tends to be. If a gene has rare variants with multiple consequences, they will appear in all the corresponding panels.

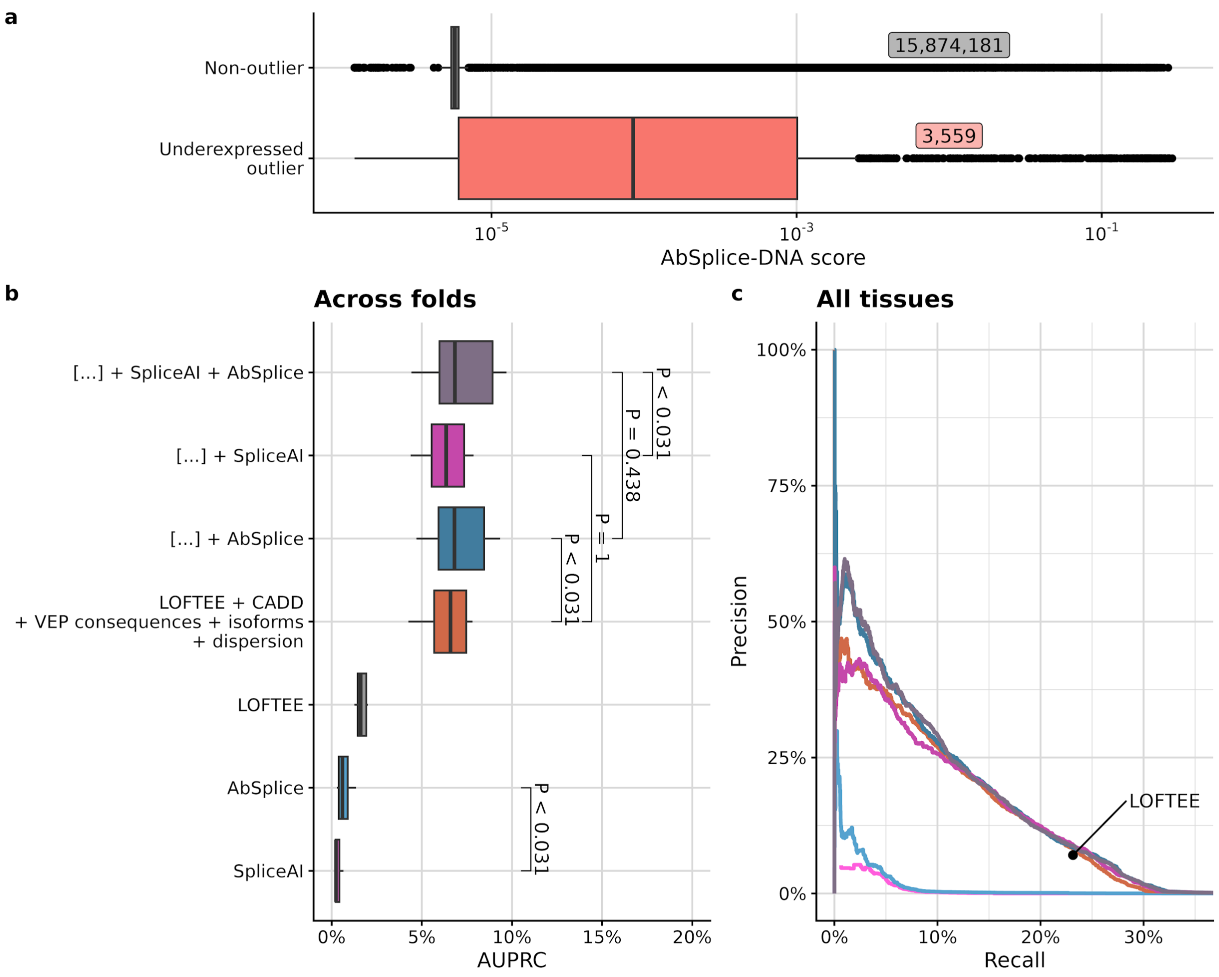


**Supplementary Fig. S8**: **AbSplice-DNA improves the outlier prediction.** **(a)** Distribution of AbSplice-DNA scores in different outlier classes. The numbers of data points are labeled for each box. (**b)** Distribution of average precision (AUPRC) across 27 GTEx tissue types. P-values were obtained using the paired Wilcoxon test. Adding AbSplice-DNA as a feature significantly improves outlier prediction, whereas adding SpliceAI does not show any significant improvement. (**c)** Precision-recall curve for all tissues combined. **All box plots:** Center line, median; box limits, first and third quartiles; whiskers span all data within 1.5 interquartile ranges of the lower and upper quartiles.

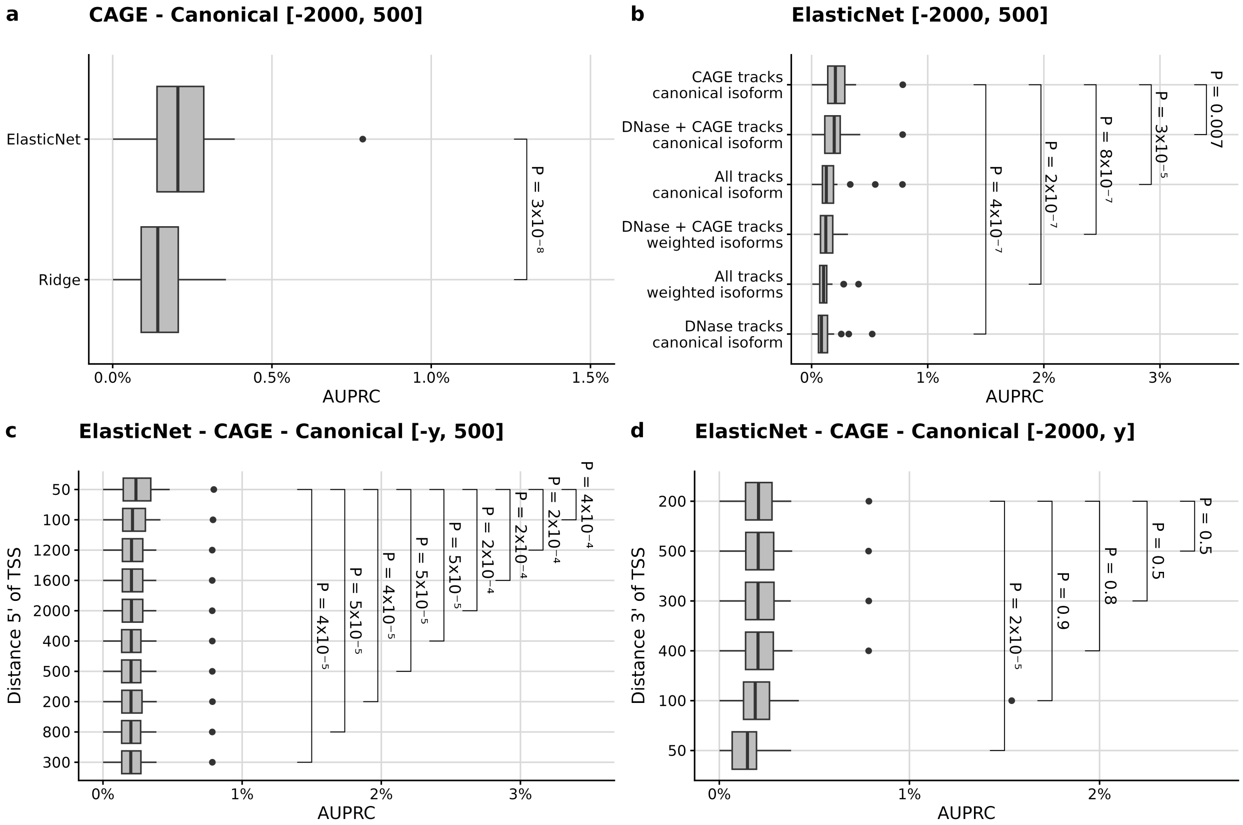


**Supplementary Fig. S9: Integration of Enformer.** We trained models predicting gene expression averaged across individuals of GTEx for each of the 49 tissues from the reference genome. The models are all regularized linear regressions from some of the Enformer-predicted tracks (CAGE, DNAse, or all tracks) applied to either the canonical isoform only or to all isoforms weighted by their tissue-specific isoform composition. **(a)** Distribution of average precision (AUPRC) on underexpression prediction across 27 GTEx tissue types for elastic net (top) or ridge (bottom) canonical-isoform models using CAGE tracks. Variant effects were considered for variants within [-2000, +500] bp of the canonical transcription start site (TSS) and set to 0 otherwise. Elastic net, which is a generalization of ridge, outperformed ridge and was further used. **(b)** As in **a** for canonical-isoform and weighted-isoform elastic net models and including various Enformer-predicted tracks. Using CAGE predictions on the canonical TSS performs significantly better than other combinations and was further used. **(c)** as in **a** setting to 0 the variant effect predictions for variants beyond a certain distance (y-axis) of the TSS. Keeping effect predictions only for variants within 50 bp 5' of the TSS and setting effects to 0 otherwise resulted in the best median AUPRC. **(d)** as in **c** but 3' of the TSS. The 200 bp threshold led to the best median AUPRC. All box plots show the distribution of average precision (AUPRC) across 27 GTEx tissue types, sorted by the respective median AUPRCs. Center line, median; box limits, first and third quartiles; whiskers span all data within 1.5 interquartile ranges of the lower and upper quartiles. P-values were obtained using the paired Wilcoxon test.

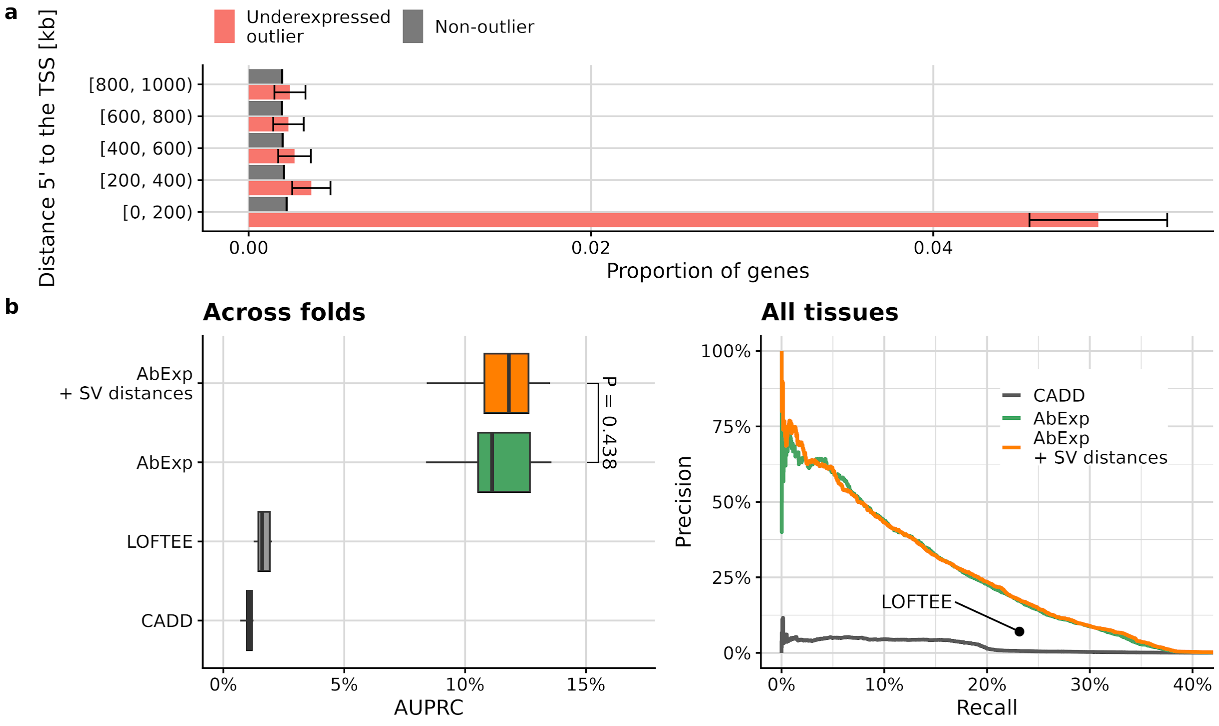


**Supplementary Fig. S10: Distal structural variants do not significantly improve underexpression outlier prediction. (a)** Proportion of underexpressed outliers (red), and non-outliers (gray) with a structural variant within a certain distance 5' of the transcription start site of the gene (rows, Methods). Error bars mark 95% binomial confidence intervals. **(b)** Left: Distribution of average precision (AUPRC) across 27 GTEx tissue types, using from bottom to top: CADD, LOFTEE, AbExp (which only has SVs within 5kb of the gene), and "AbExp + SV distances", a model integrating the AbExp features and SVs up to 1 Mb 5' of the gene stratified by distance bins as in **a)**. P-values were obtained using the paired Wilcoxon test. Center line, median; box limits, first and third quartiles; whiskers span all data within 1.5 interquartile ranges of the lower and upper quartiles. Right: Precision-recall curve for all tissues combined for the same models.

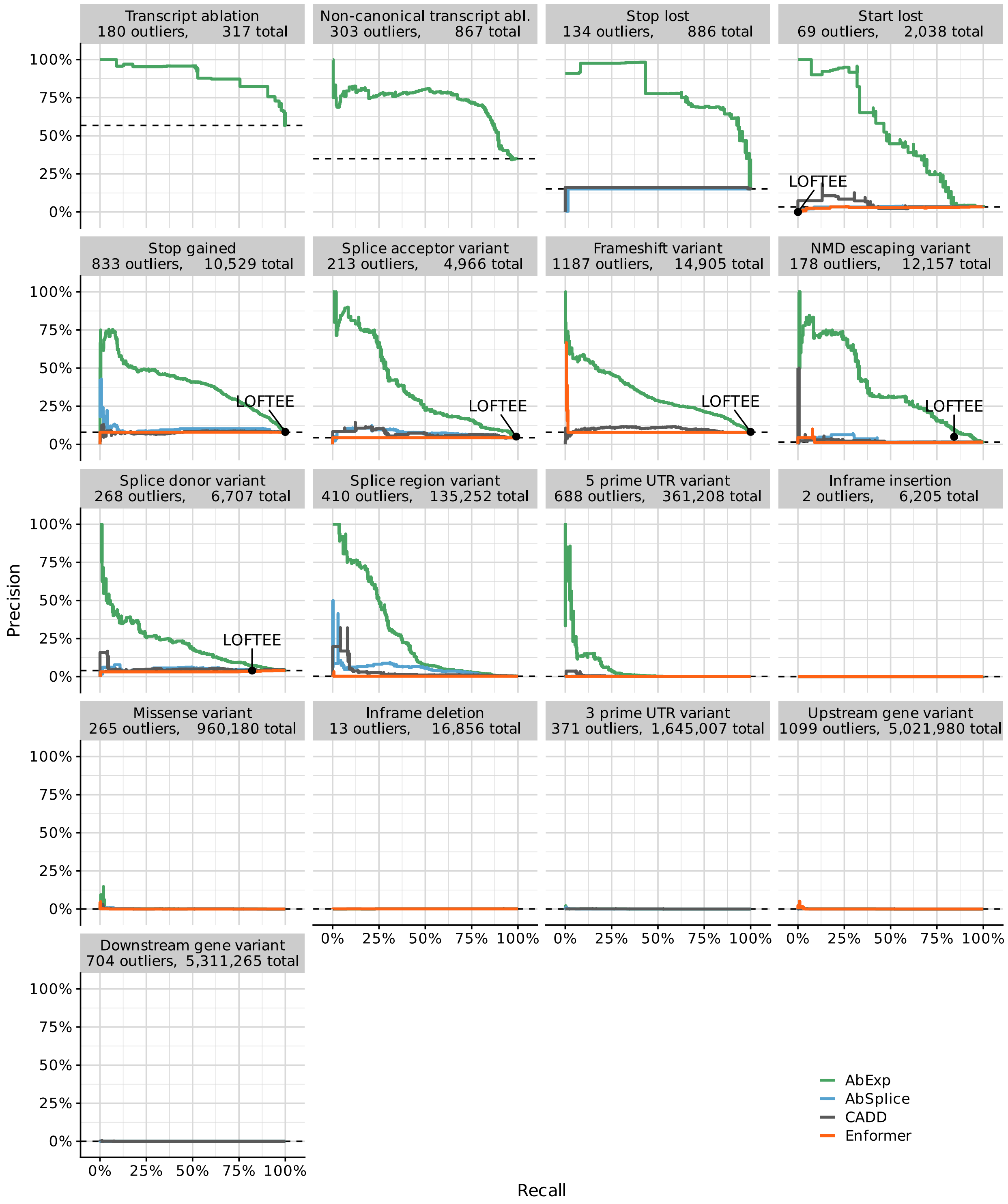


**Supplementary Fig. S11**: **Precision-recall curves stratified by variant type for different models on the GTEx benchmark dataset.** Variant effect predictions were max-aggregated per gene, individual and tissue for all models. LOFTEE as a binary predictor is displayed as a single point. Horizontal dashed lines show the baseline precision for a predictor that classifies all genes as outliers.

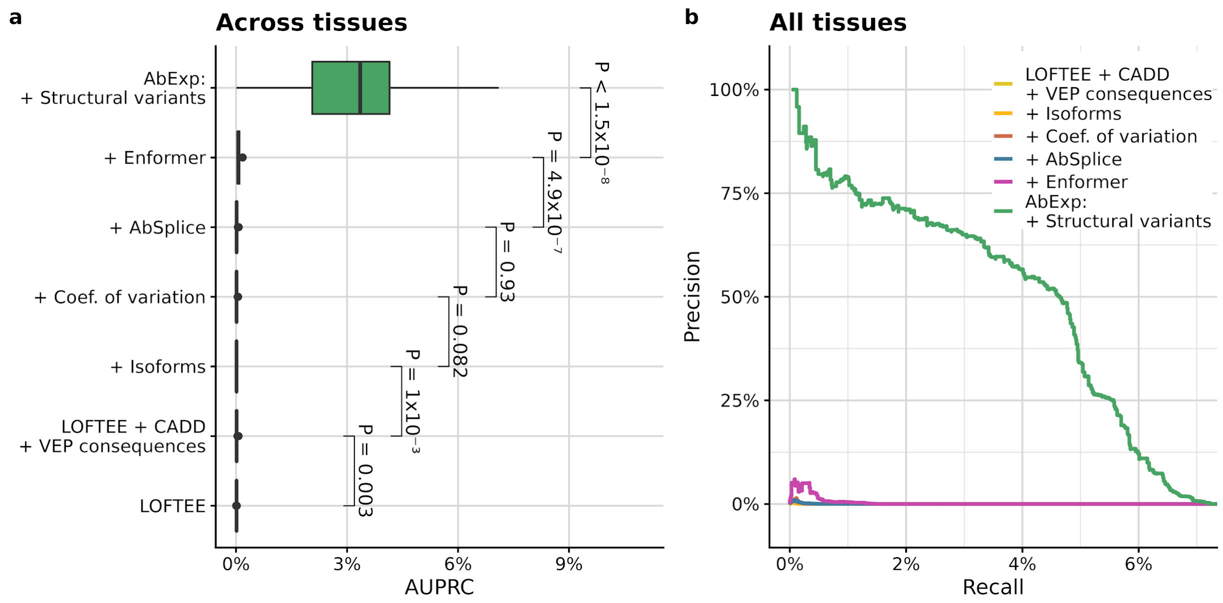


**Supplementary Fig. S12: Integrating Enformer and duplications significantly improves the overexpression outlier prediction. (a)** Distribution of average precision (AUPRC) across 27 GTEx tissue types. P-values were obtained using the paired Wilcoxon test. Center line, median; box limits, first and third quartiles; whiskers span all data within 1.5 interquartile ranges of the lower and upper quartiles. **(b)** Precision-recall curve for all tissues combined.

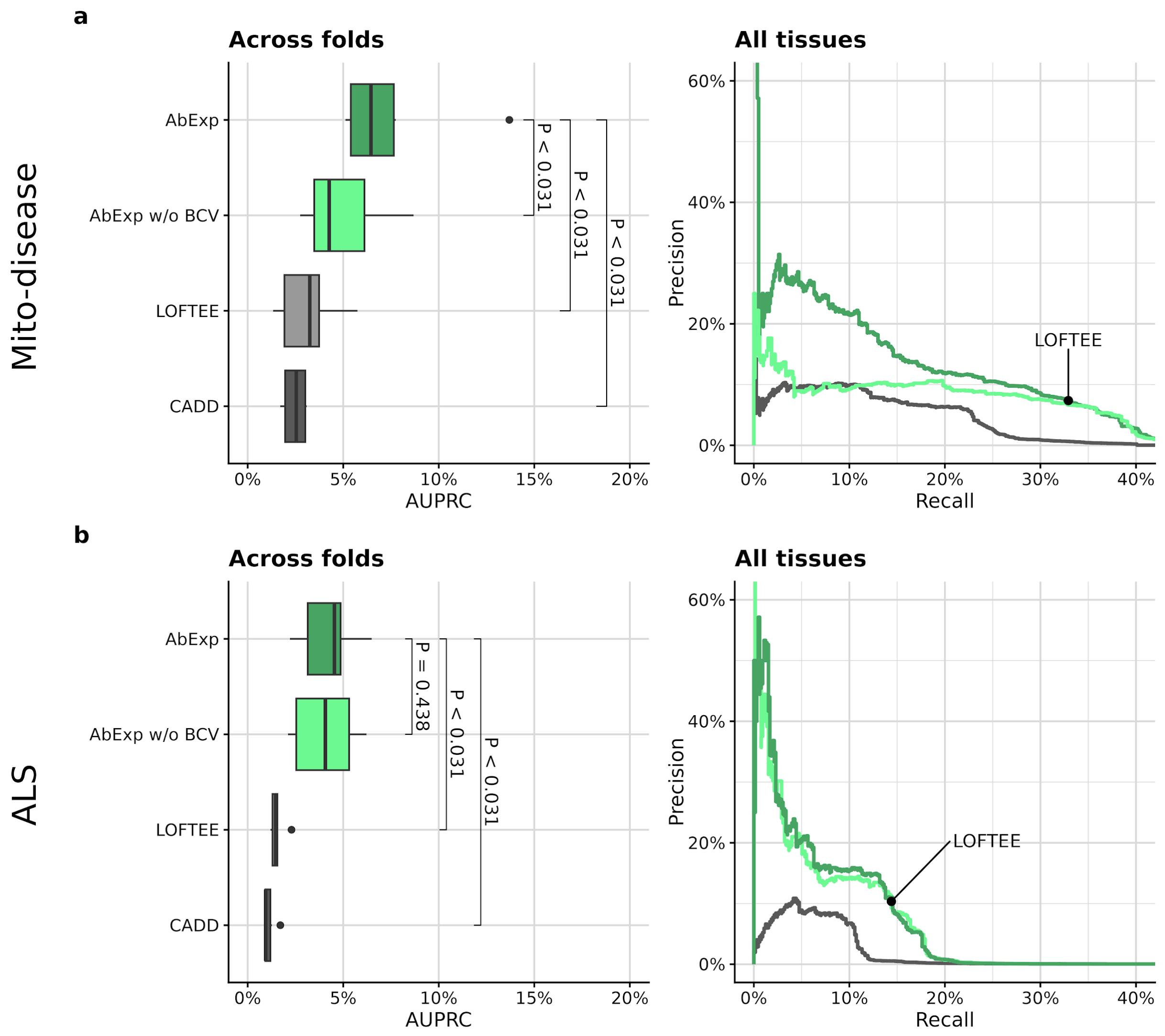


**Supplementary Fig. S13: Performance of AbExp on independent datasets. (a)** Left: Distribution of average precision (AUPRC) across five cross-validation folds in the mito-disease dataset. *P*-values were obtained using a paired Wilcoxon test. Right: Precision-recall curve on the whole mito-disease dataset. LOFTEE as a binary predictor is shown as a single point. **(b)** Left: Distribution of average precision (AUPRC) across five cross-validation folds in the ALS dataset. P-values were obtained using a paired Wilcoxon test. Right: Precision-recall curve on the whole ALS dataset. LOFTEE as a binary predictor is shown as a single point. **All box plots:** Center line, median; box limits, first and third quartiles; whiskers span all data within 1.5 interquartile ranges of the lower and upper quartiles.

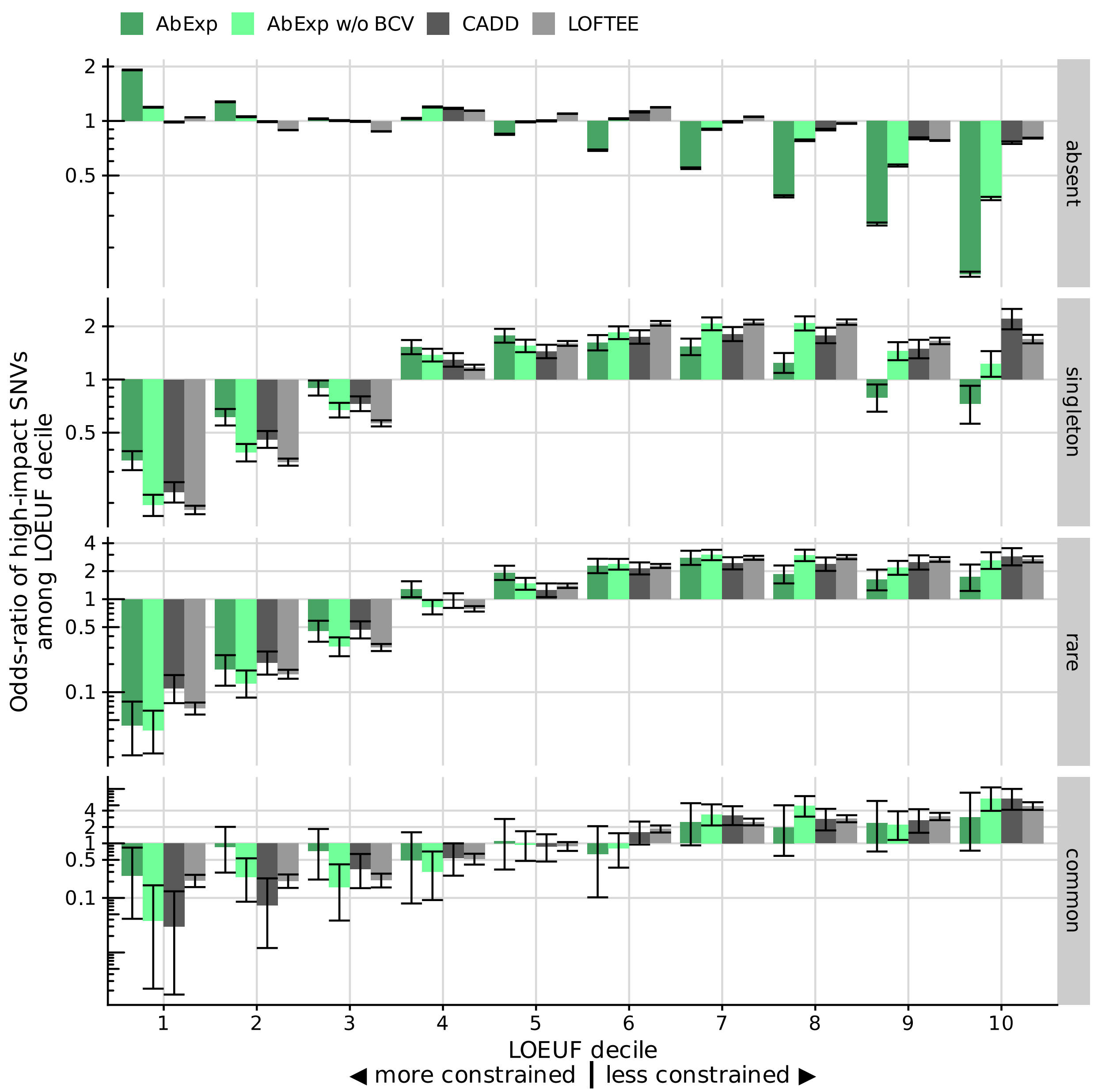


**Supplementary Fig. S14: Odds-ratio of high-impact variants among absent, singleton, rare (MAF < 0.1%) and common SNVs in gnomAD as a function of gene LOEUF decile.** Genes with a high LOEUF are more tolerant to loss of function. The analysis is restricted to variants within 5 kb of protein-coding genes. The high-impact cutoffs for CADD and AbExp without BCV were set to match the quantile of the high-impact cutoff of AbExp. Error bars show Wald 95% confidence intervals from logistic regression fits.

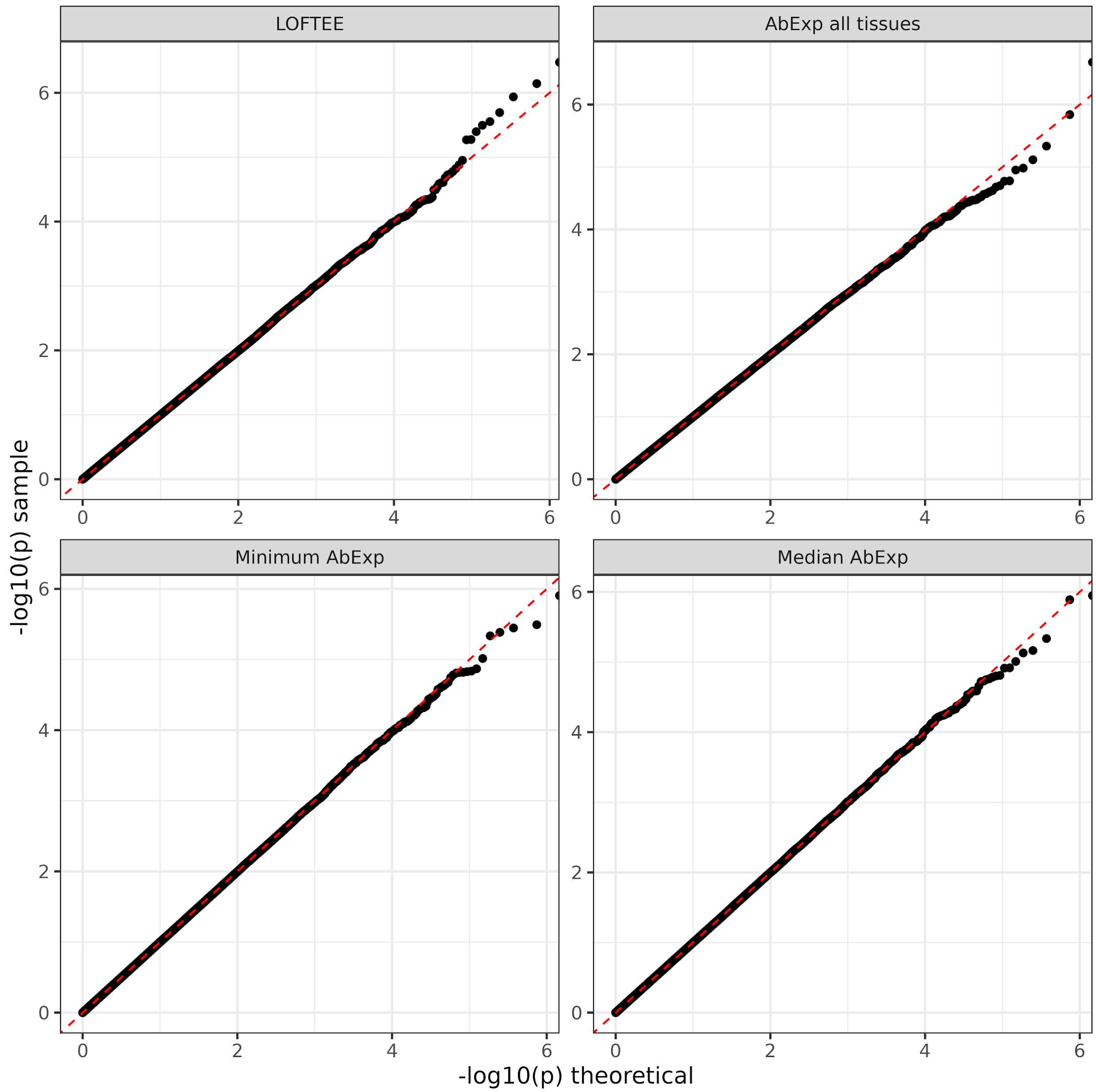


**Supplementary Fig. S15: Quantile-quantile plot of *p*-values on randomized data against random uniform distribution across all traits facetted by model.** P-values on randomized data were obtained by shuffling the phenotype randomly once and testing for associations. The data aligns on the diagonal indicating calibration of all four models. Across all the four models and 40 traits, we found only two reported associations at FDR < 0.05 on these permuted data, one for AbExp in the "Total bilirubin" trait and one for LOFTEE in the "IGF1" trait.

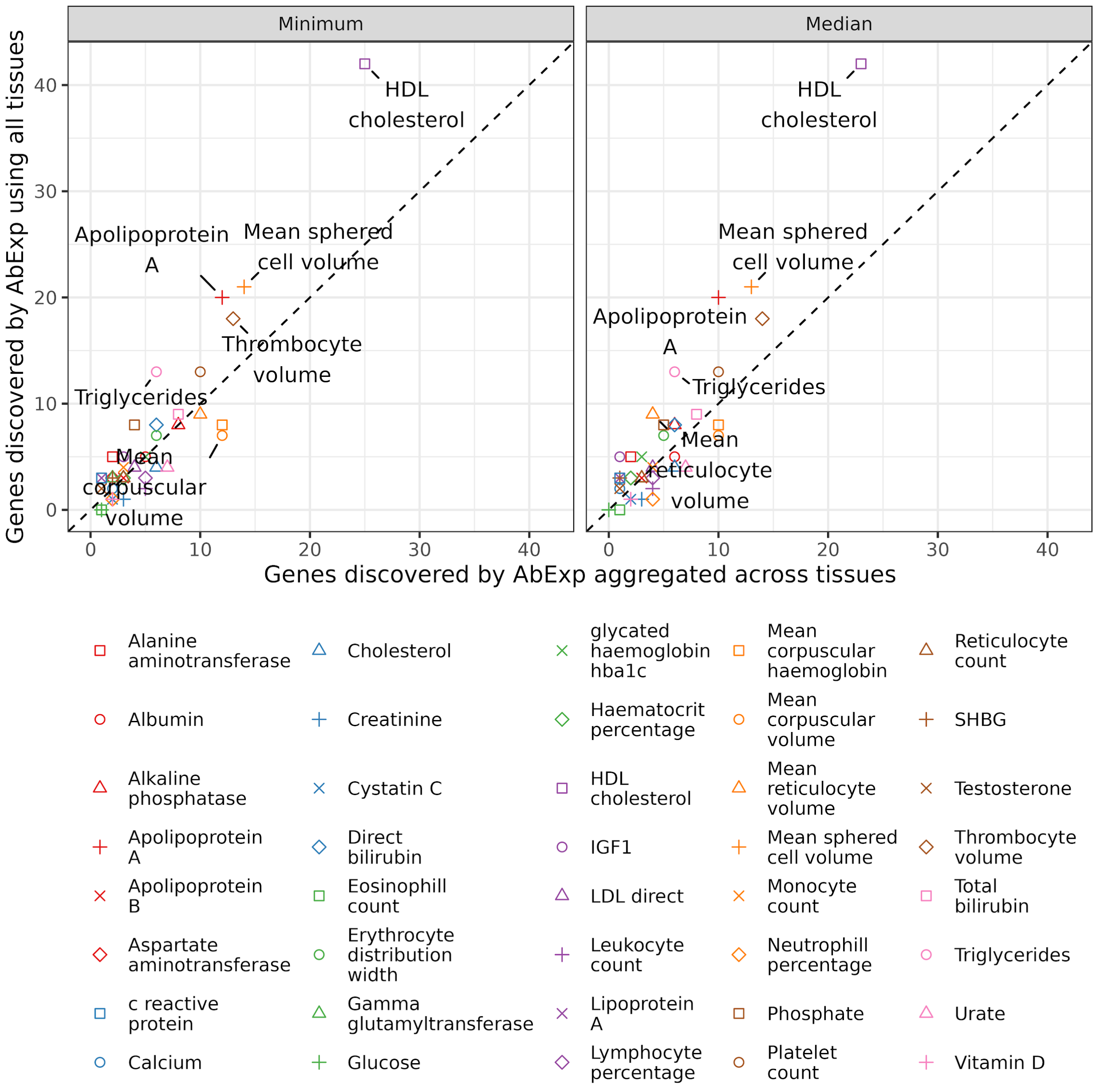


**Supplementary Fig. S16**: **Number of significant blood-trait-associated genes for different AbExp aggregations.** The y-axis shows the number of significantly associated genes when using AbExp scores of all tissues as variables in the regression test. The x-axis shows the corresponding number of significantly associated genes when using an aggregated AbExp score as variable in the regression test. ‘Minimum’ is using the minimum predicted score across tissues per gene, ‘Median’ uses the median score across tissues.

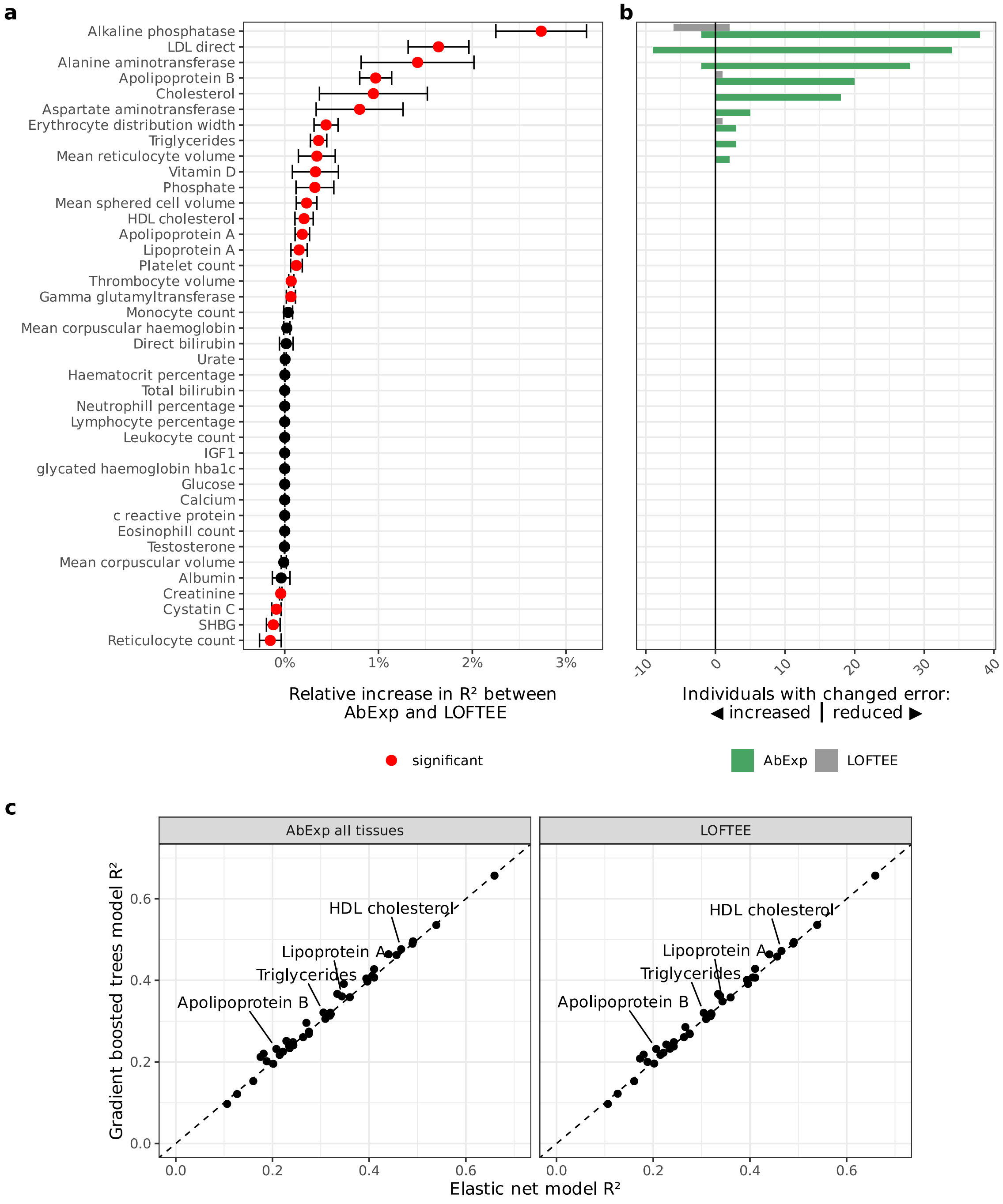


**Supplementary Fig. S17**: **AbExp improvements over LOFTEE in phenotype prediction also hold with elastic-net based predictions. (a)** Relative $R^2$ increase between AbExp-based and LOFTEE-based predictions across traits. Traits with a significant difference between both models are marked red (two-sided paired t-test, nominal $P < 0.05$). Error bars show the standard deviation among 5 cross-validation folds. **(b)** Positive bars show the number of individuals with an error reduced by at least one standard deviation in the trait scale and therefore improved prediction, negative bars show the number of individuals with an error increased by at least one standard deviation in the trait scale and therefore worse prediction of the AbExp-based model (green) and the LOFTEE-based model (grey). **(c)** $R^2$ of gradient boosted trees models against $R^2$ of elastic net models across traits, when using AbExp scores or LOFTEE. All data presented in this figure are computed on held-out folds of a 5-fold cross-validation within a third of the UKBB data.
