## Supplementary tables for "Aberrant expression prediction across human tissues"

| | Individuals | Genes | Tissues | Samples | Underexpressed outliers | Non-outliers | Overexpressed outliers |
|---|---|---|---|---|---|---|---|
| Unfiltered | 946 | 33615 | 49 | 16213 | 27399 | 285256273 | 44987 |
| Samples with whole genomes | 633 | 33615 | 49 | 11215 | 19051 | 197083281 | 31637 |
| Keep only protein-coding genes | 633 | 18563 | 49 | 11215 | 15320 | 147883919 | 26338 |
| Remove samples with many outliers | 633 | 18563 | 49 | 10999 | 13873 | 145015257 | 20047 |
| Keep only genes of samples that have sufficiently large expected number of reads ('mu' > 450) | 633 | 18171 | 49 | 10999 | 11200 | 99434253 | 14464 |

| phenotype | pgs_id | pgs_score_file_path | clumping_var_file_path | genebass_ phenocode |
|---|---|---|---|---|
| Alanine_aminotransferase | PGS001940 | /s/project/uk_biobank/processed/ risk_scores/PGS001940/PGS001940.sscore | /s/project/uk_biobank/processed/ clumping/30620/ GWAS_variants_clumped_mac.parquet | 30620 |
| Albumin | PGS001886 | /s/project/uk_biobank/processed/ risk_scores/PGS001886/PGS001886.sscore | /s/project/uk_biobank/processed/ clumping/30600/ GWAS_variants_clumped_mac.parquet | 30600 |
| Alkaline_phosphatase | PGS001939 | /s/project/uk_biobank/processed/ risk_scores/PGS001939/PGS001939.sscore | /s/project/uk_biobank/processed/ clumping/30610/ GWAS_variants_clumped_mac.parquet | 30610 |
| Apolipoprotein_A | PGS001888 | /s/project/uk_biobank/processed/ risk_scores/PGS001888/PGS001888.sscore | /s/project/uk_biobank/processed/ clumping/30630/ GWAS_variants_clumped_mac.parquet | 30630 |
| Apolipoprotein_B | PGS001889 | /s/project/uk_biobank/processed/ risk_scores/PGS001889/PGS001889.sscore | /s/project/uk_biobank/processed/ clumping/30640/ GWAS_variants_clumped_mac.parquet | 30640 |
| Aspartate_aminotransferase | PGS001941 | /s/project/uk_biobank/processed/ risk_scores/PGS001941/PGS001941.sscore | /s/project/uk_biobank/processed/ clumping/30650/ GWAS_variants_clumped_mac.parquet | 30650 |
| c_reactive_protein | PGS001946 | /s/project/uk_biobank/processed/ risk_scores/PGS001946/PGS001946.sscore | /s/project/uk_biobank/processed/ clumping/30710/ GWAS_variants_clumped_mac.parquet | 30710 |
| Calcium | PGS001893 | /s/project/uk_biobank/processed/ risk_scores/PGS001893/PGS001893.sscore | /s/project/uk_biobank/processed/ clumping/30680/ GWAS_variants_clumped_mac.parquet | 30680 |
| Cholesterol | PGS001895 | /s/project/uk_biobank/processed/ risk_scores/PGS001895/PGS001895.sscore | /s/project/uk_biobank/processed/ clumping/30690/ GWAS_variants_clumped_mac.parquet | 30690 |

| | | | | |
|---|---|---|---|---|
| Creatinine | PGS001945 | /s/project/uk_biobank/processed/ risk_scores/PGS001945/PGS001945.sscore | /s/project/uk_biobank/processed/ clumping/30700/ GWAS_variants_clumped_mac.parquet | 30700 |
| Cystatin_C | PGS001947 | /s/project/uk_biobank/processed/ risk_scores/PGS001947/PGS001947.sscore | /s/project/uk_biobank/processed/ clumping/30720/ GWAS_variants_clumped_mac.parquet | 30720 |
| Direct_bilirubin | PGS001942 | /s/project/uk_biobank/processed/ risk_scores/PGS001942/PGS001942.sscore | /s/project/uk_biobank/processed/ clumping/30660/ GWAS_variants_clumped_mac.parquet | 30660 |
| Eosinophill_count | PGS001172 | /s/project/uk_biobank/processed/ risk_scores/PGS001172/PGS001172.sscore | /s/project/uk_biobank/processed/ clumping/30150/ GWAS_variants_clumped_mac.parquet | 30150 |
| Erythrocyte_distribution_width | PGS001908 | /s/project/uk_biobank/processed/ risk_scores/PGS001908/PGS001908.sscore | /s/project/uk_biobank/processed/ clumping/30070/ GWAS_variants_clumped_mac.parquet | 30070 |
| Gamma_glutamyltransferase | PGS001964 | /s/project/uk_biobank/processed/ risk_scores/PGS001964/PGS001964.sscore | /s/project/uk_biobank/processed/ clumping/30730/ GWAS_variants_clumped_mac.parquet | 30730 |
| Glucose | PGS001952 | /s/project/uk_biobank/processed/ risk_scores/PGS001952/PGS001952.sscore | /s/project/uk_biobank/processed/ clumping/30740/ GWAS_variants_clumped_mac.parquet | 30740 |
| glycated_haemoglobin_hba1c | PGS001953 | /s/project/uk_biobank/processed/ risk_scores/PGS001953/PGS001953.sscore | /s/project/uk_biobank/processed/ clumping/30750/ GWAS_variants_clumped_mac.parquet | 30750 |
| Haematocrit_percentage | PGS001925 | /s/project/uk_biobank/processed/ risk_scores/PGS001925/PGS001925.sscore | /s/project/uk_biobank/processed/ clumping/30030/ GWAS_variants_clumped_mac.parquet | 30030 |
| HDL_cholesterol | PGS001954 | /s/project/uk_biobank/processed/ risk_scores/PGS001954/PGS001954.sscore | /s/project/uk_biobank/processed/ clumping/30760/ GWAS_variants_clumped_mac.parquet | 30760 |

| | | | | |
|---|---|---|---|---|
| IGF1 | PGS001960 | /s/project/uk_biobank/processed/ risk_scores/PGS001960/PGS001960.sscore | /s/project/uk_biobank/processed/ clumping/30770/ GWAS_variants_clumped_mac.parquet | 30770 |
| LDL_direct | PGS001933 | /s/project/uk_biobank/processed/ risk_scores/PGS001933/PGS001933.sscore | /s/project/uk_biobank/processed/ clumping/30780/ GWAS_variants_clumped_mac.parquet | 30780 |
| Leukocyte_count | PGS001962 | /s/project/uk_biobank/processed/ risk_scores/PGS001962/PGS001962.sscore | /s/project/uk_biobank/processed/ clumping/30000/ GWAS_variants_clumped_mac.parquet | 30000 |
| Lipoprotein_A | PGS001963 | /s/project/uk_biobank/processed/ risk_scores/PGS001963/PGS001963.sscore | /s/project/uk_biobank/processed/ clumping/30790/ GWAS_variants_clumped_mac.parquet | 30790 |
| Lymphocyte_percentage | PGS001986 | /s/project/uk_biobank/processed/ risk_scores/PGS001986/PGS001986.sscore | /s/project/uk_biobank/processed/ clumping/30180/ GWAS_variants_clumped_mac.parquet | 30180 |
| Mean_corpuscular_haemoglobin | PGS001989 | /s/project/uk_biobank/processed/ risk_scores/PGS001989/PGS001989.sscore | /s/project/uk_biobank/processed/ clumping/30050/ GWAS_variants_clumped_mac.parquet | 30050 |
| Mean_corpuscular_volume | PGS001990 | /s/project/uk_biobank/processed/ risk_scores/PGS001990/PGS001990.sscore | /s/project/uk_biobank/processed/ clumping/30040/ GWAS_variants_clumped_mac.parquet | 30040 |
| Mean_reticulocyte_volume | PGS000987 | /s/project/uk_biobank/processed/ risk_scores/PGS000987/PGS000987.sscore | /s/project/uk_biobank/processed/ clumping/30260/ GWAS_variants_clumped_mac.parquet | 30260 |
| Mean_sphered_cell_volume | PGS002008 | /s/project/uk_biobank/processed/ risk_scores/PGS002008/PGS002008.sscore | /s/project/uk_biobank/processed/ clumping/30270/ GWAS_variants_clumped_mac.parquet | 30270 |
| Monocyte_count | PGS001968 | /s/project/uk_biobank/processed/ risk_scores/PGS001968/PGS001968.sscore | /s/project/uk_biobank/processed/ clumping/30130/ GWAS_variants_clumped_mac.parquet | 30130 |

| | | | | |
|---|---|---|---|---|
| Neutrophill_percentage | PGS001997 | /s/project/uk_biobank/processed/ risk_scores/PGS001997/PGS001997.sscore | /s/project/uk_biobank/processed/ clumping/30200/ GWAS_variants_clumped_mac.parquet | 30200 |
| Phosphate | PGS001998 | /s/project/uk_biobank/processed/ risk_scores/PGS001998/PGS001998.sscore | /s/project/uk_biobank/processed/ clumping/30810/ GWAS_variants_clumped_mac.parquet | 30810 |
| Platelet_count | PGS001973 | /s/project/uk_biobank/processed/ risk_scores/PGS001973/PGS001973.sscore | /s/project/uk_biobank/processed/ clumping/30080/ GWAS_variants_clumped_mac.parquet | 30080 |
| Reticulocyte_count | PGS001528 | /s/project/uk_biobank/processed/ risk_scores/PGS001528/PGS001528.sscore | /s/project/uk_biobank/processed/ clumping/30250/ GWAS_variants_clumped_mac.parquet | 30250 |
| SHBG | PGS001977 | /s/project/uk_biobank/processed/ risk_scores/PGS001977/PGS001977.sscore | /s/project/uk_biobank/processed/ clumping/30830/ GWAS_variants_clumped_mac.parquet | 30830 |
| Testosterone | PGS001988 | /s/project/uk_biobank/processed/ risk_scores/PGS001988/PGS001988.sscore | /s/project/uk_biobank/processed/ clumping/30850/ GWAS_variants_clumped_mac.parquet | 30850 |
| Testosterone | PGS001914 | /s/project/uk_biobank/processed/ risk_scores/PGS001914/PGS001914.sscore | /s/project/uk_biobank/processed/ clumping/30850/ GWAS_variants_clumped_mac.parquet | 30850 |
| Thrombocyte_volume | PGS001971 | /s/project/uk_biobank/processed/ risk_scores/PGS001971/PGS001971.sscore | /s/project/uk_biobank/processed/ clumping/30100/ GWAS_variants_clumped_mac.parquet | 30100 |
| Total_bilirubin | PGS001942 | /s/project/uk_biobank/processed/ risk_scores/PGS001942/PGS001942.sscore | /s/project/uk_biobank/processed/ clumping/30840/ GWAS_variants_clumped_mac.parquet | 30840 |
| Triglycerides | PGS001979 | /s/project/uk_biobank/processed/ risk_scores/PGS001979/PGS001979.sscore | /s/project/uk_biobank/processed/ clumping/30870/ GWAS_variants_clumped_mac.parquet | 30870 |

| | | | | |
|---|---|---|---|---|
| Urate | PGS002010 | /s/project/uk_biobank/processed/risk_scores/PGS002010/PGS002010.sscore | /s/project/uk_biobank/processed/clumping/30880/GWAS_variants_clumped_mac.parquet | 30880 |
| Vitamin_D | PGS001982 | /s/project/uk_biobank/processed/risk_scores/PGS001982/PGS001982.sscore | /s/project/uk_biobank/processed/clumping/30890/GWAS_variants_clumped_mac.parquet | 30890 |

| phenotype | feature set | gene | gene name | LR test statistic | degrees of freedom | nominal p-value | adjusted p-value | number of observations | tissue with the largest effect |
|---|---|---|---|---|---|---|---|---|---|
| Alanine_aminotransferase | AbExp_all_tissues | ENSG00000167701 | GPT | 1379.81566 | 44 | 1.9793E-260 | 3.67811E-256 | 102827 | Stomach |
| Alanine_aminotransferase | AbExp_all_tissues | ENSG00000150753 | CCT5 | 133.461924 | 48 | 5.55666E-10 | 1.032595E-05 | 102827 | Brain - Amygdala |
| Alanine_aminotransferase | AbExp_all_tissues | ENSG00000167700 | MFSD3 | 128.570765 | 48 | 2.76809E-09 | 5.143939E-05 | 102827 | Brain - Caudate (basal ganglia) |
| Alanine_aminotransferase | AbExp_all_tissues | ENSG00000146151 | HMGCLL1 | 64.5122374 | 18 | 3.74274E-07 | 0.0069551269 | 102827 | Breast - Mammary Tissue |
| Alanine_aminotransferase | AbExp_all_tissues | ENSG00000079385 | CEACAM1 | 76.2496021 | 28 | 2.38319E-06 | 0.0442867577 | 102827 | Uterus |
| Albumin | AbExp_all_tissues | ENSG00000163631 | ALB | 377.640249 | 43 | 4.57514E-55 | 8.501982E-51 | 94914 | Testis |
| Albumin | AbExp_all_tissues | ENSG00000104870 | FCGRT | 163.532934 | 48 | 1.60953E-14 | 2.99099E-10 | 94914 | Muscle - Skeletal |
| Albumin | AbExp_all_tissues | ENSG00000145703 | IQGAP2 | 108.770917 | 33 | 5.09402E-10 | 9.466213E-06 | 94914 | Minor Salivary Gland |
| Albumin | AbExp_all_tissues | ENSG00000127377 | CRYGN | 29.3726665 | 2 | 4.18607E-07 | 0.0077789741 | 94914 | Artery - Tibial |
| Albumin | AbExp_all_tissues | ENSG00000181830 | SLC35C1 | 108.812099 | 48 | 1.28434E-06 | 0.0238669001 | 94914 | Minor Salivary Gland |
| Alkaline_phosphatase | AbExp_all_tissues | ENSG00000162551 | ALPL | 3304.4726 | 46 | 0 | 0 | 102858 | Brain - Cortex |
| Alkaline_phosphatase | AbExp_all_tissues | ENSG00000112293 | GPLD1 | 325.690332 | 35 | 7.67263E-49 | 1.425805E-44 | 102858 | Muscle - Skeletal |
| Alkaline_phosphatase | AbExp_all_tissues | ENSG00000141505 | ASGR1 | 265.83316 | 18 | 4.84078E-46 | 8.995626E-42 | 102858 | Lung |
| Alkaline_phosphatase | AbExp_all_tissues | ENSG00000244038 | DDOST | 217.037396 | 48 | 2.38318E-23 | 4.428669E-19 | 102858 | Pancreas |
| Alkaline_phosphatase | AbExp_all_tissues | ENSG00000072195 | SPEG | 113.802147 | 44 | 4.27455E-08 | 0.0007943389 | 102858 | Brain - Cerebellar Hemisphere |
| Alkaline_phosphatase | AbExp_all_tissues | ENSG00000139044 | B4GALNT3 | 97.2102216 | 34 | 5.30613E-08 | 0.0009860376 | 102858 | Small Intestine - Terminal Ileum |
| Alkaline_phosphatase | AbExp_all_tissues | ENSG00000181577 | C6orf223 | 45.3074822 | 7 | 1.19193E-07 | 0.0022149679 | 102858 | Artery - Aorta |
| Alkaline_phosphatase | AbExp_all_tissues | ENSG00000073734 | ABCB11 | 39.670587 | 6 | 5.28701E-07 | 0.009824848 | 102858 | Testis |
| Apolipoprotein_A | AbExp_all_tissues | ENSG00000165029 | ABCA1 | 485.543137 | 47 | 3.51049E-74 | 6.52355E-70 | 94312 | Artery - Coronary |
| Apolipoprotein_A | AbExp_all_tissues | ENSG00000213398 | LCAT | 270.997705 | 48 | 7.17193E-33 | 1.33276E-28 | 94312 | Whole Blood |
| Apolipoprotein_A | AbExp_all_tissues | ENSG00000132855 | ANGPTL3 | 97.3735982 | 2 | 7.1712E-22 | 1.332624E-17 | 94312 | Adipose - Visceral (Omentum) |
| Apolipoprotein_A | AbExp_all_tissues | ENSG00000166819 | PLIN1 | 167.511239 | 41 | 3.19453E-17 | 5.936399E-13 | 94312 | Ovary |
| Apolipoprotein_A | AbExp_all_tissues | ENSG00000118137 | APOA1 | 121.063142 | 24 | 6.27139E-15 | 1.165413E-10 | 94312 | Liver |
| Apolipoprotein_A | AbExp_all_tissues | ENSG00000164597 | COG5 | 161.639968 | 48 | 3.18683E-14 | 5.922095E-10 | 94312 | Brain - Putamen (basal ganglia) |
| Apolipoprotein_A | AbExp_all_tissues | ENSG00000087237 | CETP | 87.2034177 | 11 | 5.87524E-14 | 1.091796E-09 | 94312 | Brain - Hippocampus |
| Apolipoprotein_A | AbExp_all_tissues | ENSG00000101670 | LIPG | 97.4305869 | 16 | 1.04751E-13 | 1.946594E-09 | 94312 | Lung |
| Apolipoprotein_A | AbExp_all_tissues | ENSG00000126337 | KRT36 | 57.2327325 | 2 | 3.7331E-13 | 6.937227E-09 | 94312 | Artery - Coronary |
| Apolipoprotein_A | AbExp_all_tissues | ENSG00000076555 | ACACB | 137.027119 | 46 | 5.563E-11 | 1.033772E-06 | 94312 | Brain - Frontal Cortex (BA9) |
| Apolipoprotein_A | AbExp_all_tissues | ENSG00000153094 | BCL2L11 | 128.237413 | 45 | 6.29908E-10 | 1.170559E-05 | 94312 | Small Intestine - Terminal Ileum |
| Apolipoprotein_A | AbExp_all_tissues | ENSG00000123739 | PLA2G12A | 129.107936 | 47 | 1.37893E-09 | 2.562466E-05 | 94312 | Brain - Cortex |
| Apolipoprotein_A | AbExp_all_tissues | ENSG00000166035 | LIPC | 38.6536926 | 2 | 4.04071E-09 | 7.508845E-05 | 94312 | Muscle - Skeletal |
| Apolipoprotein_A | AbExp_all_tissues | ENSG00000171227 | TMEM37 | 93.2625938 | 31 | 3.76215E-08 | 0.0006991207 | 94312 | Lung |
| Apolipoprotein_A | AbExp_all_tissues | ENSG00000206549 | AC109583.1 | 74.173071 | 22 | 1.43553E-07 | 0.0026676373 | 94312 | Brain - Cerebellar Hemisphere |
| Apolipoprotein_A | AbExp_all_tissues | ENSG00000153558 | FBXL2 | 99.1512501 | 38 | 2.32737E-07 | 0.004324946 | 94312 | Brain - Cerebellum |
| Apolipoprotein_A | AbExp_all_tissues | ENSG00000136160 | EDNRB | 109.289876 | 46 | 4.55995E-07 | 0.0084737464 | 94312 | Brain - Cortex |
| Apolipoprotein_A | AbExp_all_tissues | ENSG00000172869 | DMXL1 | 110.07107 | 47 | 5.66914E-07 | 0.0105349628 | 94312 | Brain - Substantia nigra |
| Apolipoprotein_A | AbExp_all_tissues | ENSG00000105717 | PBX4 | 78.7673396 | 28 | 1.01723E-06 | 0.0189032488 | 94312 | Muscle - Skeletal |
| Apolipoprotein_A | AbExp_all_tissues | ENSG00000110245 | APOC3 | 49.6964664 | 12 | 1.57948E-06 | 0.0293514983 | 94312 | Testis |
| Apolipoprotein_B | AbExp_all_tissues | ENSG00000084674 | APOB | 464.523981 | 8 | 2.85301E-95 | 5.301755E-91 | 102371 | Liver |
| Apolipoprotein_B | AbExp_all_tissues | ENSG00000169174 | PCSK9 | 302.612428 | 16 | 7.33738E-55 | 1.363505E-50 | 102371 | Liver |
| Apolipoprotein_B | AbExp_all_tissues | ENSG00000130164 | LDLR | 181.658914 | 48 | 2.01702E-17 | 3.748233E-13 | 102371 | Adrenal Gland |
| Aspartate_aminotransferase | AbExp_all_tissues | ENSG00000120053 | GOT1 | 876.138808 | 48 | 1.3039E-152 | 2.42297E-148 | 102516 | Colon - Sigmoid |
| Aspartate_aminotransferase | AbExp_all_tissues | ENSG00000119946 | CNNM1 | 124.904631 | 24 | 1.28708E-15 | 2.391772E-11 | 102516 | Cells - EBV-transformed lymphocytes |
| Aspartate_aminotransferase | AbExp_all_tissues | ENSG00000171714 | ANO5 | 84.1118798 | 33 | 2.40238E-06 | 0.0446434591 | 102516 | Minor Salivary Gland |
| Calcium | AbExp_all_tissues | ENSG00000163631 | ALB | 146.62511 | 43 | 3.10229E-13 | 5.764985E-09 | 94857 | Liver |
| Calcium | AbExp_all_tissues | ENSG00000036828 | CASR | 35.039477 | 1 | 3.23088E-09 | 6.003949E-05 | 94857 | Prostate |

| | | | | | | | | | |
|---|---|---|---|---|---|---|---|---|---|
| Cholesterol | AbExp_all_tissues | ENSG00000084674 | APOB | 425.786273 | 8 | 5.67745E-87 | 1.055041E-82 | 102851 | Liver |
| Cholesterol | AbExp_all_tissues | ENSG00000169174 | PCSK9 | 222.916505 | 16 | 1.77643E-38 | 3.301135E-34 | 102851 | Liver |
| Cholesterol | AbExp_all_tissues | ENSG00000132855 | ANGPTL3 | 60.1689875 | 2 | 8.59944E-14 | 1.598035E-09 | 102851 | Brain - Putamen (basal ganglia) |
| Cholesterol | AbExp_all_tissues | ENSG00000130164 | LDLR | 125.642672 | 48 | 7.13149E-09 | 0.0001325244 | 102851 | Adrenal Gland |
| Creatinine | AbExp_all_tissues | ENSG00000171049 | FPR2 | 37.3028908 | 5 | 5.20776E-07 | 0.0096775737 | 102797 | Spleen |
| Cystatin_C | AbExp_all_tissues | ENSG00000171049 | FPR2 | 37.9810884 | 5 | 3.80639E-07 | 0.0070734137 | 102797 | Spleen |
| Direct_bilirubin | AbExp_all_tissues | ENSG00000244474 | UGT1A4 | 92.9948074 | 15 | 2.73097E-13 | 5.074967E-09 | 87754 | Minor Salivary Gland |
| Direct_bilirubin | AbExp_all_tissues | ENSG00000241635 | UGT1A1 | 82.7923812 | 14 | 8.55747E-12 | 1.590235E-07 | 87754 | Testis |
| Direct_bilirubin | AbExp_all_tissues | ENSG00000167165 | UGT1A6 | 73.592619 | 12 | 6.77377E-11 | 1.25877E-06 | 87754 | Brain - Caudate (basal ganglia) |
| Direct_bilirubin | AbExp_all_tissues | ENSG00000244122 | UGT1A7 | 68.6314702 | 12 | 5.77035E-10 | 1.072304E-05 | 87754 | Brain - Cortex |
| Direct_bilirubin | AbExp_all_tissues | ENSG00000084674 | APOB | 53.6587953 | 8 | 8.03933E-09 | 0.0001493949 | 87754 | Liver |
| Direct_bilirubin | AbExp_all_tissues | ENSG00000242366 | UGT1A8 | 64.1299684 | 15 | 4.85147E-08 | 0.0009015486 | 87754 | Prostate |
| Direct_bilirubin | AbExp_all_tissues | ENSG00000004939 | SLC4A1 | 38.5505449 | 5 | 2.92455E-07 | 0.0054346951 | 87754 | Brain - Amygdala |
| Direct_bilirubin | AbExp_all_tissues | ENSG00000241119 | UGT1A9 | 56.492451 | 15 | 1.00039E-06 | 0.0185901817 | 87754 | Prostate |
| Erythrocyte_distribution_width | AbExp_all_tissues | ENSG00000105610 | KLF1 | 104.557535 | 2 | 1.97524E-23 | 3.670587E-19 | 104959 | Skin - Sun Exposed (Lower leg) |
| Erythrocyte_distribution_width | AbExp_all_tissues | ENSG00000113013 | HSPA9 | 134.779725 | 48 | 3.58684E-10 | 6.665428E-06 | 104959 | Vagina |
| Erythrocyte_distribution_width | AbExp_all_tissues | ENSG00000101310 | SEC23B | 129.902784 | 48 | 1.79299E-09 | 3.331917E-05 | 104959 | Ovary |
| Erythrocyte_distribution_width | AbExp_all_tissues | ENSG00000075651 | PLD1 | 120.61993 | 44 | 4.65549E-09 | 8.651306E-05 | 104959 | Minor Salivary Gland |
| Erythrocyte_distribution_width | AbExp_all_tissues | ENSG00000176273 | SLC35G1 | 67.3473681 | 17 | 6.14167E-08 | 0.001141306 | 104959 | Cells - Cultured fibroblasts |
| Erythrocyte_distribution_width | AbExp_all_tissues | ENSG00000170276 | HSPB2 | 107.43809 | 46 | 7.98766E-07 | 0.0148434733 | 104959 | Prostate |
| Erythrocyte_distribution_width | AbExp_all_tissues | ENSG00000187045 | TMPRSS6 | 44.1356993 | 9 | 1.33244E-06 | 0.0247607827 | 104959 | Liver |
| Gamma_glutamyltransferase | AbExp_all_tissues | ENSG00000100031 | GGT1 | 464.016854 | 45 | 5.79815E-71 | 1.077469E-66 | 102807 | Small Intestine - Terminal Ileum |
| Gamma_glutamyltransferase | AbExp_all_tissues | ENSG00000286070 | GGT1 | 216.077842 | 48 | 3.4815E-23 | 6.469672E-19 | 102807 | Brain - Putamen (basal ganglia) |
| Gamma_glutamyltransferase | AbExp_all_tissues | ENSG00000144182 | LIPT1 | 101.471064 | 37 | 6.46447E-08 | 0.0012012928 | 102807 | Testis |
| HDL_cholesterol | AbExp_all_tissues | ENSG00000171316 | CHD7 | 425.237803 | 44 | 7.53352E-64 | 1.399953E-59 | 94865 | Brain - Caudate (basal ganglia) |
| HDL_cholesterol | AbExp_all_tissues | ENSG00000165029 | ABCA1 | 401.506956 | 47 | 8.8276E-58 | 1.640433E-53 | 94865 | Esophagus - Mucosa |
| HDL_cholesterol | AbExp_all_tissues | ENSG00000172594 | SMPDL3A | 364.956308 | 47 | 9.02595E-51 | 1.677292E-46 | 94865 | Testis |
| HDL_cholesterol | AbExp_all_tissues | ENSG00000129250 | KIF1C | 291.724004 | 48 | 1.21615E-36 | 2.259979E-32 | 94865 | Brain - Nucleus accumbens (basal ganglia) |
| HDL_cholesterol | AbExp_all_tissues | ENSG00000213398 | LCAT | 263.936 | 48 | 1.34171E-31 | 2.493292E-27 | 94865 | Cells - EBV-transformed lymphocytes |
| HDL_cholesterol | AbExp_all_tissues | ENSG00000180626 | ZNF594 | 195.767772 | 20 | 7.7199E-31 | 1.434589E-26 | 94865 | Spleen |
| HDL_cholesterol | AbExp_all_tissues | ENSG00000085415 | SEH1L | 254.89886 | 47 | 2.3645E-30 | 4.393952E-26 | 94865 | Esophagus - Mucosa |
| HDL_cholesterol | AbExp_all_tissues | ENSG00000178297 | TMPRSS9 | 147.327601 | 6 | 2.8411E-29 | 5.279609E-25 | 94865 | Brain - Cerebellum |
| HDL_cholesterol | AbExp_all_tissues | ENSG00000087237 | CETP | 150.308883 | 11 | 1.28889E-26 | 2.39514E-22 | 94865 | Artery - Aorta |
| HDL_cholesterol | AbExp_all_tissues | ENSG00000255346 | NOX5 | 119.857698 | 3 | 8.28102E-26 | 1.538862E-21 | 94865 | Testis |
| HDL_cholesterol | AbExp_all_tissues | ENSG00000110245 | APOC3 | 124.058573 | 12 | 9.56492E-21 | 1.77745E-16 | 94865 | Pituitary |
| HDL_cholesterol | AbExp_all_tissues | ENSG00000150753 | CCT5 | 197.101663 | 48 | 5.693E-20 | 1.057931E-15 | 94865 | Colon - Transverse |
| HDL_cholesterol | AbExp_all_tissues | ENSG00000118137 | APOA1 | 135.994642 | 24 | 1.26072E-17 | 2.342801E-13 | 94865 | Adrenal Gland |
| HDL_cholesterol | AbExp_all_tissues | ENSG00000062282 | DGAT2 | 163.97044 | 38 | 1.38354E-17 | 2.571037E-13 | 94865 | Testis |
| HDL_cholesterol | AbExp_all_tissues | ENSG00000162928 | PEX13 | 162.331246 | 47 | 1.31857E-14 | 2.450291E-10 | 94865 | Adipose - Visceral (Omentum) |
| HDL_cholesterol | AbExp_all_tissues | ENSG00000133731 | IMPA1 | 161.800995 | 48 | 3.00729E-14 | 5.58844E-10 | 94865 | Esophagus - Gastroesophageal Junction |
| HDL_cholesterol | AbExp_all_tissues | ENSG00000101670 | LIPG | 92.716935 | 16 | 7.87927E-13 | 1.464205E-08 | 94865 | Brain - Cerebellar Hemisphere |
| HDL_cholesterol | AbExp_all_tissues | ENSG00000075826 | SEC31B | 146.258098 | 48 | 7.28254E-12 | 1.353315E-07 | 94865 | Brain - Hypothalamus |
| HDL_cholesterol | AbExp_all_tissues | ENSG00000101639 | CEP192 | 142.784991 | 47 | 1.35865E-11 | 2.524772E-07 | 94865 | Brain - Putamen (basal ganglia) |
| HDL_cholesterol | AbExp_all_tissues | ENSG00000175874 | CREG2 | 77.0580467 | 12 | 1.49753E-11 | 2.782859E-07 | 94865 | Vagina |
| HDL_cholesterol | AbExp_all_tissues | ENSG00000105717 | PBX4 | 107.244145 | 28 | 3.28692E-11 | 6.108079E-07 | 94865 | Nerve - Tibial |
| HDL_cholesterol | AbExp_all_tissues | ENSG00000110243 | APOA5 | 50.7243657 | 3 | 5.60034E-11 | 1.040711E-06 | 94865 | Liver |
| HDL_cholesterol | AbExp_all_tissues | ENSG00000198231 | DDX42 | 135.091348 | 48 | 3.23315E-10 | 6.008156E-06 | 94865 | Testis |

| | | | | | | | | | |
|---|---|---|---|---|---|---|---|---|---|
| HDL_cholesterol | AbExp_all_tissues | ENSG00000092820 | EZR | 133.295413 | 48 | 5.87177E-10 | 1.091151E-05 | 94865 | Esophagus - Mucosa |
| HDL_cholesterol | AbExp_all_tissues | ENSG00000136160 | EDNRB | 129.958034 | 46 | 6.09409E-10 | 1.132465E-05 | 94865 | Brain - Cerebellar Hemisphere |
| HDL_cholesterol | AbExp_all_tissues | ENSG00000119681 | LTBP2 | 113.828667 | 37 | 9.50881E-10 | 1.767022E-05 | 94865 | Heart - Atrial Appendage |
| HDL_cholesterol | AbExp_all_tissues | ENSG00000171227 | TMEM37 | 102.005316 | 31 | 1.68257E-09 | 3.126711E-05 | 94865 | Ovary |
| HDL_cholesterol | AbExp_all_tissues | ENSG00000137824 | RMDN3 | 128.193757 | 48 | 3.1288E-09 | 5.814247E-05 | 94865 | Adrenal Gland |
| HDL_cholesterol | AbExp_all_tissues | ENSG00000166035 | LIPC | 38.7743956 | 2 | 3.80406E-09 | 7.069079E-05 | 94865 | Liver |
| HDL_cholesterol | AbExp_all_tissues | ENSG00000099804 | CDC34 | 126.042758 | 48 | 6.27074E-09 | 0.0001165292 | 94865 | Esophagus - Mucosa |
| HDL_cholesterol | AbExp_all_tissues | ENSG00000126337 | KRT36 | 37.6395978 | 2 | 6.70911E-09 | 0.0001246755 | 94865 | Adrenal Gland |
| HDL_cholesterol | AbExp_all_tissues | ENSG00000166819 | PLIN1 | 114.132382 | 41 | 8.10966E-09 | 0.0001507017 | 94865 | Artery - Aorta |
| HDL_cholesterol | AbExp_all_tissues | ENSG00000142530 | FAM71E1 | 96.5111175 | 32 | 2.16701E-08 | 0.000402696 | 94865 | Brain - Hippocampus |
| HDL_cholesterol | AbExp_all_tissues | ENSG00000227124 | ZNF717 | 70.9890514 | 18 | 3.0719E-08 | 0.0005708519 | 94865 | Brain - Amygdala |
| HDL_cholesterol | AbExp_all_tissues | ENSG00000076555 | ACACB | 116.862023 | 46 | 4.34205E-08 | 0.0008068829 | 94865 | Brain - Cerebellum |
| HDL_cholesterol | AbExp_all_tissues | ENSG00000028310 | BRD9 | 116.600656 | 48 | 1.22747E-07 | 0.0022809984 | 94865 | Minor Salivary Gland |
| HDL_cholesterol | AbExp_all_tissues | ENSG00000165972 | CCDC38 | 27.6466692 | 1 | 1.45623E-07 | 0.0027061048 | 94865 | Adipose - Visceral (Omentum) |
| HDL_cholesterol | AbExp_all_tissues | ENSG00000100591 | AHSA1 | 114.037801 | 48 | 2.68786E-07 | 0.0049948564 | 94865 | Prostate |
| HDL_cholesterol | AbExp_all_tissues | ENSG00000111707 | SUDS3 | 113.771847 | 48 | 2.91383E-07 | 0.0054147792 | 94865 | Thyroid |
| HDL_cholesterol | AbExp_all_tissues | ENSG00000132855 | ANGPTL3 | 29.5901892 | 2 | 3.75467E-07 | 0.0069773077 | 94865 | Adipose - Subcutaneous |
| HDL_cholesterol | AbExp_all_tissues | ENSG00000206549 | AC109583.1 | 67.5333116 | 22 | 1.6073E-06 | 0.0298684809 | 94865 | Artery - Aorta |
| HDL_cholesterol | AbExp_all_tissues | ENSG00000065413 | ANKRD44 | 100.618611 | 44 | 2.52567E-06 | 0.0469345767 | 94865 | Stomach |
| Haematocrit_percentage | AbExp_all_tissues | ENSG00000129167 | TPH1 | 46.4138969 | 10 | 1.20664E-06 | 0.0224229646 | 104960 | Artery - Coronary |
| Haematocrit_percentage | AbExp_all_tissues | ENSG00000078725 | BRINP1 | 68.4513364 | 23 | 2.10408E-06 | 0.0391000628 | 104960 | Esophagus - Mucosa |
| Haematocrit_percentage | AbExp_all_tissues | ENSG00000162775 | RBM15 | 88.9988233 | 36 | 2.21521E-06 | 0.041165209 | 104960 | Colon - Sigmoid |
| IGF1 | AbExp_all_tissues | ENSG00000099769 | IGFALS | 81.6811246 | 8 | 2.24164E-14 | 4.16564E-10 | 102326 | Lung |
| IGF1 | AbExp_all_tissues | ENSG00000143379 | SETDB1 | 146.061753 | 48 | 7.79389E-12 | 1.448339E-07 | 102326 | Ovary |
| IGF1 | AbExp_all_tissues | ENSG00000107164 | FUBP3 | 132.501353 | 47 | 4.47353E-10 | 8.313163E-06 | 102326 | Liver |
| IGF1 | AbExp_all_tissues | ENSG00000139324 | TMTC3 | 120.168217 | 43 | 3.17467E-09 | 5.899498E-05 | 102326 | Esophagus - Muscularis |
| IGF1 | AbExp_all_tissues | ENSG00000146674 | IGFBP3 | 113.200394 | 47 | 2.18979E-07 | 0.00406929 | 102326 | Cells - Cultured fibroblasts |
| LDL_direct | AbExp_all_tissues | ENSG00000084674 | APOB | 505.563644 | 8 | 4.503E-104 | 8.36789E-100 | 102696 | Liver |
| LDL_direct | AbExp_all_tissues | ENSG00000169174 | PCSK9 | 284.226405 | 16 | 4.66448E-51 | 8.668006E-47 | 102696 | Liver |
| LDL_direct | AbExp_all_tissues | ENSG00000130164 | LDLR | 164.890514 | 48 | 9.84347E-15 | 1.829212E-10 | 102696 | Adrenal Gland |
| LDL_direct | AbExp_all_tissues | ENSG00000132855 | ANGPTL3 | 53.2580008 | 2 | 2.72378E-12 | 5.061601E-08 | 102696 | Muscle - Skeletal |
| Leukocyte_count | AbExp_all_tissues | ENSG00000254709 | IGLL5 | 124.254856 | 34 | 3.28517E-12 | 6.104836E-08 | 104960 | Skin - Not Sun Exposed (Suprapubic) |
| Leukocyte_count | AbExp_all_tissues | ENSG00000125910 | S1PR4 | 28.153491 | 2 | 7.701E-07 | 0.0143107677 | 104960 | Brain - Substantia nigra |
| Lipoprotein_A | AbExp_all_tissues | ENSG00000198670 | LPA | 153.238913 | 6 | 1.598E-30 | 2.969559E-26 | 82169 | Stomach |
| Lipoprotein_A | AbExp_all_tissues | ENSG00000026652 | AGPAT4 | 173.065133 | 42 | 7.74658E-18 | 1.439547E-13 | 82169 | Colon - Transverse |
| Lipoprotein_A | AbExp_all_tissues | ENSG00000146477 | SLC22A3 | 114.580778 | 43 | 2.00182E-08 | 0.0003719973 | 82169 | Skin - Not Sun Exposed (Suprapubic) |
| Lymphocyte_percentage | AbExp_all_tissues | ENSG00000254709 | IGLL5 | 122.210251 | 34 | 7.04106E-12 | 1.308441E-07 | 104807 | Breast - Mammary Tissue |
| Lymphocyte_percentage | AbExp_all_tissues | ENSG00000198216 | CACNA1E | 65.5043388 | 15 | 2.78724E-08 | 0.0005179522 | 104807 | Brain - Spinal cord (cervical c-1) |
| Lymphocyte_percentage | AbExp_all_tissues | ENSG00000215301 | DDX3X | 111.211733 | 48 | 6.29941E-07 | 0.0117061949 | 104807 | Skin - Sun Exposed (Lower leg) |
| Mean_corpuscular_haemoglobin | AbExp_all_tissues | ENSG00000244734 | HBB | 212.920376 | 44 | 5.26898E-24 | 9.79135E-20 | 104960 | Cells - Cultured fibroblasts |
| Mean_corpuscular_haemoglobin | AbExp_all_tissues | ENSG00000105610 | KLF1 | 95.5897502 | 2 | 1.74964E-21 | 3.251362E-17 | 104960 | Brain - Hypothalamus |
| Mean_corpuscular_haemoglobin | AbExp_all_tissues | ENSG00000188536 | HBA2 | 107.727743 | 25 | 3.03805E-12 | 5.645611E-08 | 104960 | Whole Blood |
| Mean_corpuscular_haemoglobin | AbExp_all_tissues | ENSG00000072274 | TFRC | 138.415588 | 48 | 1.06063E-10 | 1.970968E-06 | 104960 | Prostate |
| Mean_corpuscular_haemoglobin | AbExp_all_tissues | ENSG00000106327 | TFR2 | 76.7391742 | 14 | 1.1335E-10 | 2.106389E-06 | 104960 | Breast - Mammary Tissue |
| Mean_corpuscular_haemoglobin | AbExp_all_tissues | ENSG00000187045 | TMPRSS6 | 63.3055534 | 9 | 3.07811E-10 | 5.720045E-06 | 104960 | Liver |
| Mean_corpuscular_haemoglobin | AbExp_all_tissues | ENSG00000130193 | THEM6 | 113.436003 | 48 | 3.22599E-07 | 0.0059948544 | 104960 | Brain - Cerebellar Hemisphere |
| Mean_corpuscular_haemoglobin | AbExp_all_tissues | ENSG00000129173 | E2F8 | 54.1631949 | 15 | 2.46683E-06 | 0.0458411036 | 104960 | Muscle - Skeletal |

| | | | | | | | | | |
|---|---|---|---|---|---|---|---|---|---|
| Mean_corpuscular_volume | AbExp_all_tissues | ENSG00000244734 | HBB | 216.619476 | 43 | 5.22054E-25 | 9.701331E-21 | 104959 | Cells - Cultured fibroblasts |
| Mean_corpuscular_volume | AbExp_all_tissues | ENSG00000105610 | KLF1 | 75.8573314 | 2 | 3.37124E-17 | 6.264777E-13 | 104959 | Esophagus - Muscularis |
| Mean_corpuscular_volume | AbExp_all_tissues | ENSG00000188536 | HBA2 | 103.010599 | 27 | 8.20234E-11 | 1.524241E-06 | 104959 | Skin - Not Sun Exposed (Suprapubic) |
| Mean_corpuscular_volume | AbExp_all_tissues | ENSG00000129173 | E2F8 | 77.3785307 | 15 | 2.09901E-10 | 3.900595E-06 | 104959 | Muscle - Skeletal |
| Mean_corpuscular_volume | AbExp_all_tissues | ENSG00000072274 | TFRC | 128.375253 | 48 | 2.9497E-09 | 5.481427E-05 | 104959 | Artery - Tibial |
| Mean_corpuscular_volume | AbExp_all_tissues | ENSG00000187045 | TMPRSS6 | 51.8871688 | 9 | 4.7479E-08 | 0.0008823031 | 104959 | Testis |
| Mean_corpuscular_volume | AbExp_all_tissues | ENSG00000106327 | TFR2 | 60.1946336 | 14 | 1.08449E-07 | 0.0020153063 | 104959 | Brain - Cortex |
| Mean_reticulocyte_volume | AbExp_all_tissues | ENSG00000004939 | SLC4A1 | 82.0566168 | 5 | 3.11459E-16 | 5.787834E-12 | 102867 | Whole Blood |
| Mean_reticulocyte_volume | AbExp_all_tissues | ENSG00000204147 | ASAH2B | 131.723329 | 35 | 3.92543E-13 | 7.294632E-09 | 102867 | Brain - Cortex |
| Mean_reticulocyte_volume | AbExp_all_tissues | ENSG00000070182 | SPTB | 110.326694 | 31 | 7.95872E-11 | 1.478969E-06 | 102867 | Brain - Hippocampus |
| Mean_reticulocyte_volume | AbExp_all_tissues | ENSG00000188611 | ASAH2 | 60.9761015 | 7 | 9.63441E-11 | 1.790362E-06 | 102867 | Testis |
| Mean_reticulocyte_volume | AbExp_all_tissues | ENSG00000084774 | CAD | 122.616414 | 48 | 1.87318E-08 | 0.0003480924 | 102867 | Cells - EBV-transformed lymphocytes |
| Mean_reticulocyte_volume | AbExp_all_tissues | ENSG00000139597 | N4BP2L1 | 114.436407 | 48 | 2.38105E-07 | 0.0044247044 | 102867 | Brain - Cortex |
| Mean_reticulocyte_volume | AbExp_all_tissues | ENSG00000243772 | KIR2DL3 | 30.5289224 | 3 | 1.06809E-06 | 0.0198483769 | 102867 | Adrenal Gland |
| Mean_reticulocyte_volume | AbExp_all_tissues | ENSG00000112077 | RHAG | 35.001264 | 5 | 1.50378E-06 | 0.0279446875 | 102867 | Spleen |
| Mean_reticulocyte_volume | AbExp_all_tissues | ENSG00000105146 | AURKC | 72.9299174 | 26 | 2.46158E-06 | 0.0457435901 | 102867 | Adipose - Subcutaneous |
| Mean_sphered_cell_volume | AbExp_all_tissues | ENSG00000204147 | ASAH2B | 218.378389 | 37 | 1.40337E-27 | 2.607876E-23 | 102817 | Brain - Anterior cingulate cortex (BA24) |
| Mean_sphered_cell_volume | AbExp_all_tissues | ENSG00000004939 | SLC4A1 | 93.4333088 | 5 | 1.27542E-18 | 2.37012E-14 | 102817 | Whole Blood |
| Mean_sphered_cell_volume | AbExp_all_tissues | ENSG00000139597 | N4BP2L1 | 183.927258 | 48 | 8.59718E-18 | 1.597614E-13 | 102817 | Brain - Cortex |
| Mean_sphered_cell_volume | AbExp_all_tissues | ENSG00000188611 | ASAH2 | 95.1148636 | 7 | 1.09769E-17 | 2.039841E-13 | 102817 | Testis |
| Mean_sphered_cell_volume | AbExp_all_tissues | ENSG00000196531 | NACA | 168.956662 | 48 | 2.23703E-15 | 4.15708E-11 | 102817 | Adipose - Subcutaneous |
| Mean_sphered_cell_volume | AbExp_all_tissues | ENSG00000105146 | AURKC | 118.472103 | 26 | 9.13908E-14 | 1.698316E-09 | 102817 | Brain - Substantia nigra |
| Mean_sphered_cell_volume | AbExp_all_tissues | ENSG00000133612 | AGAP3 | 156.62599 | 48 | 1.91729E-13 | 3.562907E-09 | 102817 | Ovary |
| Mean_sphered_cell_volume | AbExp_all_tissues | ENSG00000285625 | AC117378.1 | 154.846011 | 48 | 3.60596E-13 | 6.700951E-09 | 102817 | Brain - Cerebellar Hemisphere |
| Mean_sphered_cell_volume | AbExp_all_tissues | ENSG00000070182 | SPTB | 110.386373 | 31 | 7.78414E-11 | 1.446528E-06 | 102817 | Brain - Hippocampus |
| Mean_sphered_cell_volume | AbExp_all_tissues | ENSG00000112077 | RHAG | 54.9471049 | 5 | 1.33848E-10 | 2.487302E-06 | 102817 | Spleen |
| Mean_sphered_cell_volume | AbExp_all_tissues | ENSG00000129173 | E2F8 | 69.0294272 | 15 | 6.64747E-09 | 0.0001235299 | 102817 | Muscle - Skeletal |
| Mean_sphered_cell_volume | AbExp_all_tissues | ENSG00000180228 | PRKRA | 122.939913 | 48 | 1.6905E-08 | 0.0003141454 | 102817 | Brain - Nucleus accumbens (basal ganglia) |
| Mean_sphered_cell_volume | AbExp_all_tissues | ENSG00000004809 | SLC22A16 | 33.2386579 | 2 | 6.05783E-08 | 0.0011257267 | 102817 | Adipose - Visceral (Omentum) |
| Mean_sphered_cell_volume | AbExp_all_tissues | ENSG00000105610 | KLF1 | 33.1746633 | 2 | 6.2548E-08 | 0.0011623294 | 102817 | Spleen |
| Mean_sphered_cell_volume | AbExp_all_tissues | ENSG00000265303 | AC099850.3 | 113.819031 | 47 | 1.81091E-07 | 0.0033652187 | 102817 | Minor Salivary Gland |
| Mean_sphered_cell_volume | AbExp_all_tissues | ENSG00000243772 | KIR2DL3 | 32.4814954 | 3 | 4.14279E-07 | 0.0076985387 | 102817 | Whole Blood |
| Mean_sphered_cell_volume | AbExp_all_tissues | ENSG00000128340 | RAC2 | 107.174249 | 45 | 5.49104E-07 | 0.0102040037 | 102817 | Stomach |
| Mean_sphered_cell_volume | AbExp_all_tissues | ENSG00000179588 | ZFPM1 | 106.983896 | 45 | 5.8192E-07 | 0.010813816 | 102817 | Breast - Mammary Tissue |
| Mean_sphered_cell_volume | AbExp_all_tissues | ENSG00000244734 | HBB | 101.63699 | 43 | 1.18636E-06 | 0.0220461942 | 102817 | Cells - Cultured fibroblasts |
| Mean_sphered_cell_volume | AbExp_all_tissues | ENSG00000084774 | CAD | 107.999243 | 48 | 1.63105E-06 | 0.0303098609 | 102817 | Heart - Atrial Appendage |
| Mean_sphered_cell_volume | AbExp_all_tissues | ENSG00000271605 | MILR1 | 88.1840073 | 35 | 1.76005E-06 | 0.0327070914 | 102817 | Minor Salivary Gland |
| Monocyte_count | AbExp_all_tissues | ENSG00000177663 | IL17RA | 256.551711 | 47 | 1.19509E-30 | 2.220829E-26 | 104807 | Thyroid |
| Monocyte_count | AbExp_all_tissues | ENSG00000122025 | FLT3 | 62.5222443 | 11 | 3.14137E-09 | 5.837608E-05 | 104807 | Heart - Atrial Appendage |
| Monocyte_count | AbExp_all_tissues | ENSG00000106328 | FSCN3 | 82.5015615 | 29 | 4.95642E-07 | 0.0092105181 | 104807 | Esophagus - Muscularis |
| Monocyte_count | AbExp_all_tissues | ENSG00000151490 | PTPRO | 52.528709 | 12 | 4.99566E-07 | 0.0092834355 | 104807 | Brain - Putamen (basal ganglia) |
| Neutrophill_percentage | AbExp_all_tissues | ENSG00000254709 | IGLL5 | 113.804505 | 34 | 1.54389E-10 | 2.869014E-06 | 104807 | Breast - Mammary Tissue |
| Phosphate | AbExp_all_tissues | ENSG00000162551 | ALPL | 324.465621 | 46 | 1.50758E-43 | 2.801543E-39 | 94729 | Brain - Cortex |
| Phosphate | AbExp_all_tissues | ENSG00000198569 | SLC34A3 | 52.4900772 | 4 | 1.08946E-10 | 2.024547E-06 | 94729 | Brain - Hypothalamus |
| Phosphate | AbExp_all_tissues | ENSG00000285330 | AC126283.2 | 131.067491 | 47 | 7.21086E-10 | 1.339995E-05 | 94729 | Stomach |
| Phosphate | AbExp_all_tissues | ENSG00000123739 | PLA2G12A | 129.896226 | 47 | 1.06299E-09 | 1.975363E-05 | 94729 | Lung |
| Phosphate | AbExp_all_tissues | ENSG00000139160 | ETFBKMT | 112.946232 | 41 | 1.2031E-08 | 0.000223572 | 94729 | Esophagus - Gastroesophageal Junction |

| | | | | | | | | | |
|---|---|---|---|---|---|---|---|---|---|
| Phosphate | AbExp_all_tissues | ENSG00000196652 | ZKSCAN5 | 118.002439 | 46 | 3.02348E-08 | 0.000561853 | 94729 | Brain - Cerebellar Hemisphere |
| Phosphate | AbExp_all_tissues | ENSG00000109118 | PHF12 | 112.904992 | 48 | 3.78772E-07 | 0.0070387288 | 94729 | Brain - Caudate (basal ganglia) |
| Phosphate | AbExp_all_tissues | ENSG00000117505 | DR1 | 64.6128449 | 21 | 2.49793E-06 | 0.0464190303 | 94729 | Brain - Cerebellar Hemisphere |
| Platelet_count | AbExp_all_tissues | ENSG00000096968 | JAK2 | 252.206669 | 48 | 1.67792E-29 | 3.118077E-25 | 104959 | Brain - Cerebellar Hemisphere |
| Platelet_count | AbExp_all_tissues | ENSG00000124172 | ATP5F1E | 188.727488 | 44 | 7.73748E-20 | 1.437856E-15 | 104959 | Uterus |
| Platelet_count | AbExp_all_tissues | ENSG00000101162 | TUBB1 | 91.29985 | 8 | 2.5322E-16 | 4.705586E-12 | 104959 | Whole Blood |
| Platelet_count | AbExp_all_tissues | ENSG00000111252 | SH2B3 | 163.236696 | 48 | 1.79147E-14 | 3.329096E-10 | 104959 | Testis |
| Platelet_count | AbExp_all_tissues | ENSG00000185245 | GP1BA | 64.8004245 | 7 | 1.6493E-11 | 3.064896E-07 | 104959 | Spleen |
| Platelet_count | AbExp_all_tissues | ENSG00000169704 | GP9 | 55.7862313 | 4 | 2.22315E-11 | 4.131274E-07 | 104959 | Brain - Caudate (basal ganglia) |
| Platelet_count | AbExp_all_tissues | ENSG00000203618 | GP1BB | 77.1944186 | 14 | 9.34409E-11 | 1.736412E-06 | 104959 | Artery - Aorta |
| Platelet_count | AbExp_all_tissues | ENSG00000005961 | ITGA2B | 68.8749904 | 13 | 1.29233E-09 | 2.401538E-05 | 104959 | Ovary |
| Platelet_count | AbExp_all_tissues | ENSG00000145703 | IQGAP2 | 105.995874 | 33 | 1.3802E-09 | 2.564825E-05 | 104959 | Skin - Sun Exposed (Lower leg) |
| Platelet_count | AbExp_all_tissues | ENSG00000090534 | THPO | 71.4680977 | 19 | 5.235E-08 | 0.0009728196 | 104959 | Brain - Hypothalamus |
| Platelet_count | AbExp_all_tissues | ENSG00000179144 | GIMAP7 | 45.6156871 | 7 | 1.03838E-07 | 0.0019296255 | 104959 | Artery - Aorta |
| Platelet_count | AbExp_all_tissues | ENSG00000124089 | MC3R | 24.6220423 | 1 | 6.97493E-07 | 0.0129615194 | 104959 | Brain - Anterior cingulate cortex (BA24) |
| Platelet_count | AbExp_all_tissues | ENSG00000117400 | MPL | 28.1827368 | 2 | 7.58921E-07 | 0.0141030251 | 104959 | Brain - Substantia nigra |
| Reticulocyte_count | AbExp_all_tissues | ENSG00000004939 | SLC4A1 | 54.1528437 | 5 | 1.94958E-10 | 3.622896E-06 | 102867 | Whole Blood |
| Reticulocyte_count | AbExp_all_tissues | ENSG00000112077 | RHAG | 48.4454833 | 5 | 2.88073E-09 | 5.353253E-05 | 102867 | Artery - Aorta |
| Reticulocyte_count | AbExp_all_tissues | ENSG00000214706 | IFRD2 | 113.621165 | 48 | 3.05004E-07 | 0.0056678831 | 102867 | Pancreas |
| SHBG | AbExp_all_tissues | ENSG00000129214 | SHBG | 352.088964 | 3 | 5.26438E-76 | 9.782801E-72 | 94020 | Liver |
| SHBG | AbExp_all_tissues | ENSG00000134627 | PIWIL4 | 101.417799 | 33 | 6.98977E-09 | 0.0001298909 | 94020 | Thyroid |
| SHBG | AbExp_all_tissues | ENSG00000154165 | GPR15 | 31.9386783 | 4 | 1.96911E-06 | 0.0365918941 | 94020 | Adipose - Visceral (Omentum) |
| Testosterone | AbExp_all_tissues | ENSG00000086065 | CHMP5 | 122.279372 | 48 | 2.0842E-08 | 0.0003873064 | 93177 | Breast - Mammary Tissue |
| Testosterone | AbExp_all_tissues | ENSG00000129214 | SHBG | 34.5172037 | 3 | 1.5407E-07 | 0.0028630833 | 93177 | Liver |
| Thrombocyte_volume | AbExp_all_tissues | ENSG00000145703 | IQGAP2 | 421.072567 | 33 | 7.84677E-69 | 1.458165E-64 | 104958 | Minor Salivary Gland |
| Thrombocyte_volume | AbExp_all_tissues | ENSG00000169704 | GP9 | 104.435668 | 4 | 1.11722E-21 | 2.076136E-17 | 104958 | Brain - Caudate (basal ganglia) |
| Thrombocyte_volume | AbExp_all_tissues | ENSG00000124172 | ATP5F1E | 199.393644 | 46 | 5.36031E-21 | 9.961071E-17 | 104958 | Skin - Not Sun Exposed (Suprapubic) |
| Thrombocyte_volume | AbExp_all_tissues | ENSG00000185245 | GP1BA | 106.535844 | 7 | 4.79839E-20 | 8.916841E-16 | 104958 | Spleen |
| Thrombocyte_volume | AbExp_all_tissues | ENSG00000077044 | DGKD | 188.607198 | 48 | 1.46436E-18 | 2.72122E-14 | 104958 | Adipose - Subcutaneous |
| Thrombocyte_volume | AbExp_all_tissues | ENSG00000205436 | EXOC3L4 | 109.009912 | 14 | 8.70309E-17 | 1.617295E-12 | 104958 | Spleen |
| Thrombocyte_volume | AbExp_all_tissues | ENSG00000101162 | TUBB1 | 92.101872 | 8 | 1.73973E-16 | 3.232946E-12 | 104958 | Lung |
| Thrombocyte_volume | AbExp_all_tissues | ENSG00000165702 | GFI1B | 52.7733086 | 2 | 3.47074E-12 | 6.449669E-08 | 104958 | Whole Blood |
| Thrombocyte_volume | AbExp_all_tissues | ENSG00000071205 | ARHGAP10 | 106.142459 | 34 | 2.39729E-09 | 4.454887E-05 | 104958 | Breast - Mammary Tissue |
| Thrombocyte_volume | AbExp_all_tissues | ENSG00000203618 | GP1BB | 66.185029 | 14 | 9.3923E-09 | 0.0001745372 | 104958 | Lung |
| Thrombocyte_volume | AbExp_all_tissues | ENSG00000162738 | VANGL2 | 111.873692 | 40 | 1.00638E-08 | 0.0001870162 | 104958 | Artery - Aorta |
| Thrombocyte_volume | AbExp_all_tissues | ENSG00000139725 | RHOF | 121.668916 | 48 | 2.5277E-08 | 0.000469722 | 104958 | Pancreas |
| Thrombocyte_volume | AbExp_all_tissues | ENSG00000187800 | PEAR1 | 117.44402 | 46 | 3.61062E-08 | 0.0006709612 | 104958 | Testis |
| Thrombocyte_volume | AbExp_all_tissues | ENSG00000169607 | CKAP2L | 48.7351207 | 8 | 7.14596E-08 | 0.0013279343 | 104958 | Esophagus - Mucosa |
| Thrombocyte_volume | AbExp_all_tissues | ENSG00000119285 | HEATR1 | 116.457256 | 48 | 1.28285E-07 | 0.0023839236 | 104958 | Prostate |
| Thrombocyte_volume | AbExp_all_tissues | ENSG00000111206 | FOXM1 | 80.3991166 | 27 | 3.285E-07 | 0.0061045229 | 104958 | Small Intestine - Terminal Ileum |
| Thrombocyte_volume | AbExp_all_tissues | ENSG00000275052 | PPP4R3B | 109.302663 | 48 | 1.1112E-06 | 0.0206494641 | 104958 | Brain - Cortex |
| Thrombocyte_volume | AbExp_all_tissues | ENSG00000134690 | CDCA8 | 66.3929973 | 21 | 1.31468E-06 | 0.0244307161 | 104958 | Artery - Tibial |
| Total_bilirubin | AbExp_all_tissues | ENSG00000167165 | UGT1A6 | 163.619756 | 12 | 9.60062E-29 | 1.784084E-24 | 102463 | Stomach |
| Total_bilirubin | AbExp_all_tissues | ENSG00000241635 | UGT1A1 | 163.935937 | 14 | 1.14517E-27 | 2.128073E-23 | 102463 | Prostate |
| Total_bilirubin | AbExp_all_tissues | ENSG00000244474 | UGT1A4 | 156.939763 | 15 | 1.00328E-25 | 1.864401E-21 | 102463 | Whole Blood |
| Total_bilirubin | AbExp_all_tissues | ENSG00000242366 | UGT1A8 | 155.770725 | 15 | 1.71576E-25 | 3.188394E-21 | 102463 | Prostate |
| Total_bilirubin | AbExp_all_tissues | ENSG00000241119 | UGT1A9 | 133.142516 | 15 | 5.14715E-21 | 9.564949E-17 | 102463 | Artery - Coronary |

| | | | | | | | | | |
|---|---|---|---|---|---|---|---|---|---|
| Total_bilirubin | AbExp_all_tissues | ENSG00000244122 | UGT1A7 | 119.045849 | 12 | 9.57558E-20 | 1.77943E-15 | 102463 | Skin - Not Sun Exposed (Suprapubic) |
| Total_bilirubin | AbExp_all_tissues | ENSG00000242515 | UGT1A10 | 120.738087 | 15 | 1.36069E-18 | 2.528569E-14 | 102463 | Prostate |
| Total_bilirubin | AbExp_all_tissues | ENSG00000134538 | SLCO1B1 | 30.9581132 | 1 | 2.63658E-08 | 0.0004899549 | 102463 | Liver |
| Total_bilirubin | AbExp_all_tissues | ENSG00000004939 | SLC4A1 | 36.2709046 | 5 | 8.38364E-07 | 0.0155793124 | 102463 | Artery - Tibial |
| Triglycerides | AbExp_all_tissues | ENSG00000084674 | APOB | 283.215308 | 8 | 1.5308E-56 | 2.844689E-52 | 102769 | Liver |
| Triglycerides | AbExp_all_tissues | ENSG00000132855 | ANGPTL3 | 147.919799 | 2 | 7.57921E-33 | 1.408444E-28 | 102769 | Pancreas |
| Triglycerides | AbExp_all_tissues | ENSG00000110243 | APOA5 | 98.5894162 | 3 | 3.12446E-21 | 5.80619E-17 | 102769 | Whole Blood |
| Triglycerides | AbExp_all_tissues | ENSG00000110245 | APOC3 | 126.419565 | 12 | 3.22293E-21 | 5.989172E-17 | 102769 | Colon - Transverse |
| Triglycerides | AbExp_all_tissues | ENSG00000197614 | MFAP5 | 133.494427 | 32 | 2.34575E-14 | 4.359115E-10 | 102769 | Pancreas |
| Triglycerides | AbExp_all_tissues | ENSG00000153560 | UBP1 | 150.550113 | 48 | 1.63621E-12 | 3.04056E-08 | 102769 | Whole Blood |
| Triglycerides | AbExp_all_tissues | ENSG00000065615 | CYB5R4 | 127.653385 | 42 | 1.42355E-10 | 2.645388E-06 | 102769 | Vagina |
| Triglycerides | AbExp_all_tissues | ENSG00000170049 | KCNAB3 | 105.053034 | 30 | 2.9088E-10 | 5.405426E-06 | 102769 | Nerve - Tibial |
| Triglycerides | AbExp_all_tissues | ENSG00000132561 | MATN2 | 125.388419 | 46 | 2.77225E-09 | 5.151663E-05 | 102769 | Brain - Anterior cingulate cortex (BA24) |
| Triglycerides | AbExp_all_tissues | ENSG00000081320 | STK17B | 102.349306 | 41 | 3.71112E-07 | 0.0068963834 | 102769 | Spleen |
| Triglycerides | AbExp_all_tissues | ENSG00000140368 | PSTPIP1 | 90.1602439 | 34 | 5.59846E-07 | 0.0104036197 | 102769 | Pancreas |
| Triglycerides | AbExp_all_tissues | ENSG00000088179 | PTPN4 | 104.004795 | 45 | 1.43134E-06 | 0.0265986553 | 102769 | Esophagus - Muscularis |
| Triglycerides | AbExp_all_tissues | ENSG00000170004 | CHD3 | 106.690435 | 48 | 2.39043E-06 | 0.0444213925 | 102769 | Brain - Nucleus accumbens (basal ganglia) |
| Urate | AbExp_all_tissues | ENSG00000068831 | RASGRP2 | 157.752682 | 46 | 3.6788E-14 | 6.83631E-10 | 102742 | Cells - EBV-transformed lymphocytes |
| Urate | AbExp_all_tissues | ENSG00000109667 | SLC2A9 | 90.9019822 | 16 | 1.70592E-12 | 3.170102E-08 | 102742 | Lung |
| Urate | AbExp_all_tissues | ENSG00000174827 | PDZK1 | 51.8731194 | 12 | 6.52898E-07 | 0.0121328029 | 102742 | Pancreas |
| Urate | AbExp_all_tissues | ENSG00000171405 | XAGE5 | 24.5453602 | 1 | 7.25809E-07 | 0.0134877091 | 102742 | Testis |
| Vitamin_D | AbExp_all_tissues | ENSG00000084674 | APOB | 48.8429551 | 8 | 6.814E-08 | 0.0012662458 | 98461 | Liver |
| c_reactive_protein | AbExp_all_tissues | ENSG00000132470 | ITGB4 | 120.171482 | 45 | 9.08128E-09 | 0.0001687575 | 102635 | Brain - Putamen (basal ganglia) |
| c_reactive_protein | AbExp_all_tissues | ENSG00000144182 | LIPT1 | 108.08866 | 38 | 1.20886E-08 | 0.0002246426 | 102635 | Brain - Cerebellar Hemisphere |
| c_reactive_protein | AbExp_all_tissues | ENSG00000106636 | YKT6 | 111.76253 | 48 | 5.34158E-07 | 0.0099262657 | 102635 | Artery - Tibial |
| glycated_haemoglobin_hba1c | AbExp_all_tissues | ENSG00000112077 | RHAG | 99.7585546 | 5 | 5.94214E-20 | 1.104228E-15 | 103043 | Nerve - Tibial |
| glycated_haemoglobin_hba1c | AbExp_all_tissues | ENSG00000103335 | PIEZO1 | 161.594101 | 48 | 3.2399E-14 | 6.020706E-10 | 103043 | Brain - Caudate (basal ganglia) |
| glycated_haemoglobin_hba1c | AbExp_all_tissues | ENSG00000106633 | GCK | 88.4478437 | 13 | 2.76607E-13 | 5.140196E-09 | 103043 | Brain - Putamen (basal ganglia) |
| glycated_haemoglobin_hba1c | AbExp_all_tissues | ENSG00000159023 | EPB41 | 124.889788 | 44 | 1.12384E-09 | 2.088438E-05 | 103043 | Liver |
| glycated_haemoglobin_hba1c | AbExp_all_tissues | ENSG00000161547 | SRSF2 | 86.8672923 | 35 | 2.67431E-06 | 0.0496966829 | 103043 | Skin - Sun Exposed (Lower leg) |
| Alanine_aminotransferase | LOFTEE | ENSG00000167701 | GPT | 350.107006 | 1 | 4.02E-78 | 7.54E-74 | 102827 | not applicable |
| Alanine_aminotransferase | LOFTEE | ENSG00000079385 | CEACAM1 | 27.405765 | 1 | 1.65E-07 | 3.10E-03 | 102827 | not applicable |
| Albumin | LOFTEE | ENSG00000163631 | ALB | 292.086866 | 1 | 1.75E-65 | 3.28E-61 | 94914 | not applicable |
| Albumin | LOFTEE | ENSG00000104870 | FCGRT | 36.1543566 | 1 | 1.82E-09 | 3.42E-05 | 94914 | not applicable |
| Albumin | LOFTEE | ENSG00000223547 | ZNF844 | 32.811606 | 1 | 1.02E-08 | 0.0001906455 | 94914 | not applicable |
| Albumin | LOFTEE | ENSG00000127377 | CRYGN | 25.9638869 | 1 | 3.48E-07 | 0.0065314956 | 94914 | not applicable |
| Albumin | LOFTEE | ENSG00000145703 | IQGAP2 | 25.0432911 | 1 | 5.61E-07 | 1.05E-02 | 94914 | not applicable |
| Alkaline_phosphatase | LOFTEE | ENSG00000112293 | GPLD1 | 135.695468 | 1 | 2.33E-31 | 4.37E-27 | 102858 | not applicable |
| Alkaline_phosphatase | LOFTEE | ENSG00000162551 | ALPL | 131.854936 | 1 | 1.61E-30 | 3.02E-26 | 102858 | not applicable |
| Alkaline_phosphatase | LOFTEE | ENSG00000141505 | ASGR1 | 49.2338179 | 1 | 2.27E-12 | 4.265905E-08 | 102858 | not applicable |
| Alkaline_phosphatase | LOFTEE | ENSG00000073734 | ABCB11 | 29.7303684 | 1 | 4.96509E-08 | 0.0009322451 | 102858 | not applicable |
| Alkaline_phosphatase | LOFTEE | ENSG00000134215 | VAV3 | 23.3476237 | 1 | 1.35E-06 | 2.54E-02 | 102858 | not applicable |
| Apolipoprotein_A | LOFTEE | ENSG00000165029 | ABCA1 | 225.956113 | 1 | 4.54E-51 | 8.53E-47 | 94312 | not applicable |
| Apolipoprotein_A | LOFTEE | ENSG00000132855 | ANGPTL3 | 89.1513111 | 1 | 3.66E-21 | 6.87E-17 | 94312 | not applicable |
| Apolipoprotein_A | LOFTEE | ENSG00000087237 | CETP | 74.3375268 | 1 | 6.58E-18 | 1.236211E-13 | 94312 | not applicable |
| Apolipoprotein_A | LOFTEE | ENSG00000126337 | KRT36 | 55.2402623 | 1 | 1.07E-13 | 2.002675E-09 | 94312 | not applicable |
| Apolipoprotein_A | LOFTEE | ENSG00000166819 | PLIN1 | 41.944252 | 1 | 9.39E-11 | 1.763311E-06 | 94312 | not applicable |

| | | | | | | | | | |
|---|---|---|---|---|---|---|---|---|---|
| Apolipoprotein_A | LOFTEE | ENSG00000101670 | LIPG | 40.0204236 | 1 | 2.51E-10 | 4.72E-06 | 94312 | not applicable |
| Apolipoprotein_A | LOFTEE | ENSG00000118137 | APOA1 | 33.8236568 | 1 | 6.03E-09 | 1.13E-04 | 94312 | not applicable |
| Apolipoprotein_A | LOFTEE | ENSG00000213398 | LCAT | 24.7739314 | 1 | 6.45E-07 | 1.21E-02 | 94312 | not applicable |
| Apolipoprotein_A | LOFTEE | ENSG00000110245 | APOC3 | 23.141407 | 1 | 1.51E-06 | 2.83E-02 | 94312 | not applicable |
| Apolipoprotein_A | LOFTEE | ENSG00000142530 | FAM71E1 | 22.5233487 | 1 | 2.08E-06 | 3.90E-02 | 94312 | not applicable |
| Apolipoprotein_B | LOFTEE | ENSG00000169174 | PCSK9 | 147.016252 | 1 | 7.78E-34 | 1.46E-29 | 102371 | not applicable |
| Apolipoprotein_B | LOFTEE | ENSG00000084674 | APOB | 131.199797 | 1 | 2.24E-30 | 4.20E-26 | 102371 | not applicable |
| Apolipoprotein_B | LOFTEE | ENSG00000146592 | CREB5 | 23.8413087 | 1 | 1.05E-06 | 1.96E-02 | 102371 | not applicable |
| Apolipoprotein_B | LOFTEE | ENSG00000125844 | RRBP1 | 22.3488469 | 1 | 2.27E-06 | 4.27E-02 | 102371 | not applicable |
| Aspartate_aminotransferase | LOFTEE | ENSG00000120053 | GOT1 | 202.711342 | 1 | 5.35E-46 | 1.00E-41 | 102516 | not applicable |
| Aspartate_aminotransferase | LOFTEE | ENSG00000171714 | ANO5 | 23.2862451 | 1 | 1.40E-06 | 0.0262103125 | 102516 | not applicable |
| Calcium | LOFTEE | ENSG00000163631 | ALB | 68.7355311 | 1 | 1.13E-16 | 2.114064E-12 | 94857 | not applicable |
| Calcium | LOFTEE | ENSG00000036828 | CASR | 39.8661517 | 1 | 2.72E-10 | 5.106618E-06 | 94857 | not applicable |
| Cholesterol | LOFTEE | ENSG00000084674 | APOB | 174.547805 | 1 | 7.52E-40 | 1.411101E-35 | 102851 | not applicable |
| Cholesterol | LOFTEE | ENSG00000169174 | PCSK9 | 130.213964 | 1 | 3.68E-30 | 6.91E-26 | 102851 | not applicable |
| Cholesterol | LOFTEE | ENSG00000132855 | ANGPTL3 | 56.448345 | 1 | 5.77E-14 | 1.08E-09 | 102851 | not applicable |
| Cholesterol | LOFTEE | ENSG00000146830 | GIGYF1 | 32.4604532 | 1 | 1.22E-08 | 0.0002283967 | 102851 | not applicable |
| Cholesterol | LOFTEE | ENSG00000165029 | ABCA1 | 31.1589446 | 1 | 2.38E-08 | 4.46E-04 | 102851 | not applicable |
| Creatinine | LOFTEE | ENSG00000112499 | SLC22A2 | 44.5669479 | 1 | 2.46E-11 | 4.62E-07 | 102797 | not applicable |
| Creatinine | LOFTEE | ENSG00000008710 | PKD1 | 33.9026237 | 1 | 5.79E-09 | 1.09E-04 | 102797 | not applicable |
| Cystatin_C | LOFTEE | ENSG00000112499 | SLC22A2 | 70.1752191 | 1 | 5.43E-17 | 1.02E-12 | 102797 | not applicable |
| Cystatin_C | LOFTEE | ENSG00000008710 | PKD1 | 40.331222 | 1 | 2.14E-10 | 4.02E-06 | 102797 | not applicable |
| Direct_bilirubin | LOFTEE | ENSG00000004939 | SLC4A1 | 39.0085952 | 1 | 4.22E-10 | 7.92E-06 | 87754 | not applicable |
| Direct_bilirubin | LOFTEE | ENSG00000163554 | SPTA1 | 30.2614014 | 1 | 3.78E-08 | 7.09E-04 | 87754 | not applicable |
| Direct_bilirubin | LOFTEE | ENSG00000084674 | APOB | 26.3700503 | 1 | 2.82E-07 | 0.0052925323 | 87754 | not applicable |
| Eosinophill_count | LOFTEE | ENSG00000168769 | TET2 | 29.3487981 | 1 | 6.05E-08 | 1.14E-03 | 104807 | not applicable |
| Erythrocyte_distribution_width | LOFTEE | ENSG00000105610 | KLF1 | 126.039445 | 1 | 3.01E-29 | 5.659685E-25 | 104959 | not applicable |
| Erythrocyte_distribution_width | LOFTEE | ENSG00000113013 | HSPA9 | 66.5113723 | 1 | 3.48E-16 | 6.53181E-12 | 104959 | not applicable |
| Erythrocyte_distribution_width | LOFTEE | ENSG00000244734 | HBB | 62.5991657 | 1 | 2.53E-15 | 4.76E-11 | 104959 | not applicable |
| Erythrocyte_distribution_width | LOFTEE | ENSG00000101310 | SEC23B | 57.493089 | 1 | 3.39E-14 | 6.368236E-10 | 104959 | not applicable |
| Erythrocyte_distribution_width | LOFTEE | ENSG00000187045 | TMPRSS6 | 31.223985 | 1 | 2.30E-08 | 4.32E-04 | 104959 | not applicable |
| Erythrocyte_distribution_width | LOFTEE | ENSG00000151474 | FRMD4A | 28.1141096 | 1 | 1.14E-07 | 2.15E-03 | 104959 | not applicable |
| Erythrocyte_distribution_width | LOFTEE | ENSG00000084774 | CAD | 26.4936806 | 1 | 2.64E-07 | 4.96E-03 | 104959 | not applicable |
| Erythrocyte_distribution_width | LOFTEE | ENSG00000072274 | TFRC | 26.4349639 | 1 | 2.73E-07 | 0.0051176221 | 104959 | not applicable |
| Erythrocyte_distribution_width | LOFTEE | ENSG00000004939 | SLC4A1 | 23.2918521 | 1 | 1.39E-06 | 0.0261340141 | 104959 | not applicable |
| Erythrocyte_distribution_width | LOFTEE | ENSG00000155903 | RASA2 | 23.1979627 | 1 | 1.46E-06 | 2.74E-02 | 104959 | not applicable |
| Gamma_glutamyltransferase | LOFTEE | ENSG00000100031 | GGT1 | 89.6767835 | 1 | 2.80E-21 | 5.27E-17 | 102807 | not applicable |
| Gamma_glutamyltransferase | LOFTEE | ENSG00000170927 | PKHD1 | 37.4490231 | 1 | 9.38E-10 | 1.76E-05 | 102807 | not applicable |
| Glucose | LOFTEE | ENSG00000106633 | GCK | 28.327707 | 1 | 1.02E-07 | 1.92E-03 | 94795 | not applicable |
| Glucose | LOFTEE | ENSG00000255150 | EID3 | 22.4852704 | 1 | 2.12E-06 | 0.0397602793 | 94795 | not applicable |
| HDL_cholesterol | LOFTEE | ENSG00000165029 | ABCA1 | 168.702134 | 1 | 1.42E-38 | 2.67E-34 | 94865 | not applicable |
| HDL_cholesterol | LOFTEE | ENSG00000087237 | CETP | 119.070678 | 1 | 1.01E-27 | 1.90E-23 | 94865 | not applicable |
| HDL_cholesterol | LOFTEE | ENSG00000062282 | DGAT2 | 98.4962711 | 1 | 3.26E-23 | 6.11E-19 | 94865 | not applicable |
| HDL_cholesterol | LOFTEE | ENSG00000110245 | APOC3 | 57.2885222 | 1 | 3.76E-14 | 7.066256E-10 | 94865 | not applicable |
| HDL_cholesterol | LOFTEE | ENSG00000166819 | PLIN1 | 49.6722025 | 1 | 1.82E-12 | 3.41E-08 | 94865 | not applicable |
| HDL_cholesterol | LOFTEE | ENSG00000142530 | FAM71E1 | 47.5857423 | 1 | 5.27E-12 | 9.89E-08 | 94865 | not applicable |
| HDL_cholesterol | LOFTEE | ENSG00000110243 | APOA5 | 40.5936257 | 1 | 1.87E-10 | 3.52E-06 | 94865 | not applicable |

| | | | | | | | | | |
|---|---|---|---|---|---|---|---|---|---|
| HDL_cholesterol | LOFTEE | ENSG00000126337 | KRT36 | 33.7932395 | 1 | 6.13E-09 | 1.15E-04 | 94865 | not applicable |
| HDL_cholesterol | LOFTEE | ENSG00000132855 | ANGPTL3 | 30.5509255 | 1 | 3.25E-08 | 6.11E-04 | 94865 | not applicable |
| HDL_cholesterol | LOFTEE | ENSG00000101670 | LIPG | 29.6859116 | 1 | 5.08E-08 | 9.54E-04 | 94865 | not applicable |
| HDL_cholesterol | LOFTEE | ENSG00000152270 | PDE3B | 27.4334748 | 1 | 1.63E-07 | 3.05E-03 | 94865 | not applicable |
| HDL_cholesterol | LOFTEE | ENSG00000175445 | LPL | 27.4048313 | 1 | 1.65E-07 | 3.10E-03 | 94865 | not applicable |
| HDL_cholesterol | LOFTEE | ENSG00000165972 | CCDC38 | 25.9127998 | 1 | 3.57E-07 | 6.71E-03 | 94865 | not applicable |
| HDL_cholesterol | LOFTEE | ENSG00000118137 | APOA1 | 25.4927789 | 1 | 4.44E-07 | 8.34E-03 | 94865 | not applicable |
| HDL_cholesterol | LOFTEE | ENSG00000164978 | NUDT2 | 25.0085597 | 1 | 5.71E-07 | 1.07E-02 | 94865 | not applicable |
| HDL_cholesterol | LOFTEE | ENSG00000187855 | ASCL4 | 23.8888904 | 1 | 1.02E-06 | 1.92E-02 | 94865 | not applicable |
| HDL_cholesterol | LOFTEE | ENSG00000213398 | LCAT | 23.5904011 | 1 | 1.19E-06 | 2.24E-02 | 94865 | not applicable |
| Haematocrit_percentage | LOFTEE | ENSG00000244734 | HBB | 50.424642 | 1 | 1.24E-12 | 2.33E-08 | 104960 | not applicable |
| Haematocrit_percentage | LOFTEE | ENSG00000004939 | SLC4A1 | 29.0108375 | 1 | 7.20E-08 | 1.35E-03 | 104960 | not applicable |
| Haematocrit_percentage | LOFTEE | ENSG00000122729 | ACO1 | 24.4789417 | 1 | 7.51E-07 | 1.41E-02 | 104960 | not applicable |
| IGF1 | LOFTEE | ENSG00000140443 | IGF1R | 22.2449287 | 1 | 2.40E-06 | 4.51E-02 | 102326 | not applicable |
| LDL_direct | LOFTEE | ENSG00000084674 | APOB | 194.046317 | 1 | 4.16E-44 | 7.81E-40 | 102696 | not applicable |
| LDL_direct | LOFTEE | ENSG00000169174 | PCSK9 | 178.961637 | 1 | 8.17E-41 | 1.53E-36 | 102696 | not applicable |
| LDL_direct | LOFTEE | ENSG00000132855 | ANGPTL3 | 48.6140964 | 1 | 3.12E-12 | 5.85E-08 | 102696 | not applicable |
| Leukocyte_count | LOFTEE | ENSG00000196712 | NF1 | 35.4018432 | 1 | 2.68E-09 | 5.04E-05 | 104960 | not applicable |
| Leukocyte_count | LOFTEE | ENSG00000173575 | CHD2 | 30.1883439 | 1 | 3.92E-08 | 7.36E-04 | 104960 | not applicable |
| Leukocyte_count | LOFTEE | ENSG00000064932 | SBNO2 | 24.5768424 | 1 | 7.14E-07 | 1.34E-02 | 104960 | not applicable |
| Leukocyte_count | LOFTEE | ENSG00000185187 | SIGIRR | 23.2530945 | 1 | 1.42E-06 | 2.67E-02 | 104960 | not applicable |
| Lipoprotein_A | LOFTEE | ENSG00000198670 | LPA | 66.7703993 | 1 | 3.05E-16 | 5.727513E-12 | 82169 | not applicable |
| Lymphocyte_percentage | LOFTEE | ENSG00000173575 | CHD2 | 54.2035214 | 1 | 1.81E-13 | 3.393999E-09 | 104807 | not applicable |
| Lymphocyte_percentage | LOFTEE | ENSG00000215301 | DDX3X | 48.906911 | 1 | 2.68E-12 | 5.039541E-08 | 104807 | not applicable |
| Lymphocyte_percentage | LOFTEE | ENSG00000101745 | ANKRD12 | 35.5566277 | 1 | 2.48E-09 | 4.651617E-05 | 104807 | not applicable |
| Lymphocyte_percentage | LOFTEE | ENSG00000121966 | CXCR4 | 33.3521645 | 1 | 7.69E-09 | 0.0001443728 | 104807 | not applicable |
| Lymphocyte_percentage | LOFTEE | ENSG00000196712 | NF1 | 30.0413249 | 1 | 4.23E-08 | 0.0007941053 | 104807 | not applicable |
| Lymphocyte_percentage | LOFTEE | ENSG00000185989 | RASA3 | 22.4110857 | 1 | 2.20E-06 | 0.0413259643 | 104807 | not applicable |
| Mean_corpuscular_haemoglobin | LOFTEE | ENSG00000244734 | HBB | 143.284315 | 1 | 5.09E-33 | 9.564677E-29 | 104960 | not applicable |
| Mean_corpuscular_haemoglobin | LOFTEE | ENSG00000105610 | KLF1 | 104.376301 | 1 | 1.67E-24 | 3.141599E-20 | 104960 | not applicable |
| Mean_corpuscular_haemoglobin | LOFTEE | ENSG00000072274 | TFRC | 76.9488941 | 1 | 1.75E-18 | 3.294053E-14 | 104960 | not applicable |
| Mean_corpuscular_haemoglobin | LOFTEE | ENSG00000187045 | TMPRSS6 | 46.7231904 | 1 | 8.18E-12 | 1.535047E-07 | 104960 | not applicable |
| Mean_corpuscular_haemoglobin | LOFTEE | ENSG00000188536 | HBA2 | 36.2710603 | 1 | 1.72E-09 | 3.223737E-05 | 104960 | not applicable |
| Mean_corpuscular_haemoglobin | LOFTEE | ENSG00000127586 | CHTF18 | 30.6402506 | 1 | 3.11E-08 | 0.0005831512 | 104960 | not applicable |
| Mean_corpuscular_haemoglobin | LOFTEE | ENSG00000258366 | RTEL1 | 26.8088114 | 1 | 2.25E-07 | 0.0042172936 | 104960 | not applicable |
| Mean_corpuscular_haemoglobin | LOFTEE | ENSG00000129173 | E2F8 | 25.0423522 | 1 | 5.61E-07 | 0.0105304752 | 104960 | not applicable |
| Mean_corpuscular_haemoglobin | LOFTEE | ENSG00000113013 | HSPA9 | 24.2246864 | 1 | 8.57E-07 | 0.0160959885 | 104960 | not applicable |
| Mean_corpuscular_haemoglobin | LOFTEE | ENSG00000205250 | E2F4 | 24.0852191 | 1 | 9.22E-07 | 0.0173049068 | 104960 | not applicable |
| Mean_corpuscular_haemoglobin | LOFTEE | ENSG00000042429 | MED17 | 24.0091113 | 1 | 9.59E-07 | 1.80E-02 | 104960 | not applicable |
| Mean_corpuscular_volume | LOFTEE | ENSG00000244734 | HBB | 145.200375 | 1 | 1.94E-33 | 3.645501E-29 | 104959 | not applicable |
| Mean_corpuscular_volume | LOFTEE | ENSG00000105610 | KLF1 | 87.2864631 | 1 | 9.39E-21 | 1.762847E-16 | 104959 | not applicable |
| Mean_corpuscular_volume | LOFTEE | ENSG00000072274 | TFRC | 75.9594433 | 1 | 2.90E-18 | 5.436611E-14 | 104959 | not applicable |
| Mean_corpuscular_volume | LOFTEE | ENSG00000129173 | E2F8 | 45.9967483 | 1 | 1.18E-11 | 2.22E-07 | 104959 | not applicable |
| Mean_corpuscular_volume | LOFTEE | ENSG00000084774 | CAD | 39.7975107 | 1 | 2.82E-10 | 5.29E-06 | 104959 | not applicable |
| Mean_corpuscular_volume | LOFTEE | ENSG00000187045 | TMPRSS6 | 35.9299436 | 1 | 2.05E-09 | 3.84E-05 | 104959 | not applicable |
| Mean_corpuscular_volume | LOFTEE | ENSG00000127586 | CHTF18 | 34.7031812 | 1 | 3.84E-09 | 7.210051E-05 | 104959 | not applicable |
| Mean_corpuscular_volume | LOFTEE | ENSG00000188536 | HBA2 | 33.1164523 | 1 | 8.68E-09 | 0.0001629778 | 104959 | not applicable |

| | | | | | | | | | |
|---|---|---|---|---|---|---|---|---|---|
| Mean_corpuscular_volume | LOFTEE | ENSG00000042429 | MED17 | 29.7021415 | 1 | 5.04E-08 | 0.0009459187 | 104959 | not applicable |
| Mean_corpuscular_volume | LOFTEE | ENSG00000258366 | RTEL1 | 28.9233248 | 1 | 7.53E-08 | 1.41E-03 | 104959 | not applicable |
| Mean_corpuscular_volume | LOFTEE | ENSG00000205250 | E2F4 | 23.3238187 | 1 | 1.37E-06 | 0.0257032581 | 104959 | not applicable |
| Mean_reticulocyte_volume | LOFTEE | ENSG00000084774 | CAD | 72.0550316 | 1 | 2.09E-17 | 3.93E-13 | 102867 | not applicable |
| Mean_reticulocyte_volume | LOFTEE | ENSG00000163554 | SPTA1 | 54.0331475 | 1 | 1.97E-13 | 3.70142E-09 | 102867 | not applicable |
| Mean_reticulocyte_volume | LOFTEE | ENSG00000004939 | SLC4A1 | 39.1330383 | 1 | 3.96E-10 | 7.43E-06 | 102867 | not applicable |
| Mean_reticulocyte_volume | LOFTEE | ENSG00000177084 | POLE | 29.8410151 | 1 | 4.69E-08 | 8.81E-04 | 102867 | not applicable |
| Mean_reticulocyte_volume | LOFTEE | ENSG00000244734 | HBB | 26.8552377 | 1 | 2.19E-07 | 4.12E-03 | 102867 | not applicable |
| Mean_reticulocyte_volume | LOFTEE | ENSG00000112077 | RHAG | 26.1758234 | 1 | 3.12E-07 | 5.85E-03 | 102867 | not applicable |
| Mean_reticulocyte_volume | LOFTEE | ENSG00000127586 | CHTF18 | 25.9064687 | 1 | 3.58E-07 | 6.73E-03 | 102867 | not applicable |
| Mean_reticulocyte_volume | LOFTEE | ENSG00000070182 | SPTB | 24.2753495 | 1 | 8.35E-07 | 1.57E-02 | 102867 | not applicable |
| Mean_reticulocyte_volume | LOFTEE | ENSG00000101161 | PRPF6 | 22.9431479 | 1 | 1.67E-06 | 3.13E-02 | 102867 | not applicable |
| Mean_sphered_cell_volume | LOFTEE | ENSG00000004939 | SLC4A1 | 66.0043518 | 1 | 4.50E-16 | 8.447857E-12 | 102817 | not applicable |
| Mean_sphered_cell_volume | LOFTEE | ENSG00000244734 | HBB | 63.3627047 | 1 | 1.72E-15 | 3.228399E-11 | 102817 | not applicable |
| Mean_sphered_cell_volume | LOFTEE | ENSG00000163554 | SPTA1 | 53.8034176 | 1 | 2.22E-13 | 4.16E-09 | 102817 | not applicable |
| Mean_sphered_cell_volume | LOFTEE | ENSG00000112077 | RHAG | 40.8031815 | 1 | 1.68E-10 | 3.16E-06 | 102817 | not applicable |
| Mean_sphered_cell_volume | LOFTEE | ENSG00000084774 | CAD | 40.0654011 | 1 | 2.46E-10 | 4.61E-06 | 102817 | not applicable |
| Mean_sphered_cell_volume | LOFTEE | ENSG00000105610 | KLF1 | 34.204301 | 1 | 4.96E-09 | 9.32E-05 | 102817 | not applicable |
| Mean_sphered_cell_volume | LOFTEE | ENSG00000159023 | EPB41 | 33.4704442 | 1 | 7.24E-09 | 1.36E-04 | 102817 | not applicable |
| Mean_sphered_cell_volume | LOFTEE | ENSG00000072274 | TFRC | 31.7515503 | 1 | 1.75E-08 | 3.29E-04 | 102817 | not applicable |
| Mean_sphered_cell_volume | LOFTEE | ENSG00000070182 | SPTB | 31.0750084 | 1 | 2.48E-08 | 0.0004661089 | 102817 | not applicable |
| Mean_sphered_cell_volume | LOFTEE | ENSG00000127586 | CHTF18 | 30.2584329 | 1 | 3.78E-08 | 0.0007100024 | 102817 | not applicable |
| Mean_sphered_cell_volume | LOFTEE | ENSG00000123700 | KCNJ2 | 25.7836448 | 1 | 3.82E-07 | 0.0071707616 | 102817 | not applicable |
| Mean_sphered_cell_volume | LOFTEE | ENSG00000101161 | PRPF6 | 24.9134902 | 1 | 6.00E-07 | 0.0112583465 | 102817 | not applicable |
| Mean_sphered_cell_volume | LOFTEE | ENSG00000165424 | ZCCHC24 | 23.9319226 | 1 | 9.98E-07 | 1.87E-02 | 102817 | not applicable |
| Neutrophill_percentage | LOFTEE | ENSG00000173575 | CHD2 | 35.0963309 | 1 | 3.14E-09 | 5.89E-05 | 104807 | not applicable |
| Neutrophill_percentage | LOFTEE | ENSG00000121966 | CXCR4 | 30.3845302 | 1 | 3.54E-08 | 6.65E-04 | 104807 | not applicable |
| Neutrophill_percentage | LOFTEE | ENSG00000196712 | NF1 | 28.4104204 | 1 | 9.81E-08 | 1.84E-03 | 104807 | not applicable |
| Neutrophill_percentage | LOFTEE | ENSG00000215301 | DDX3X | 27.8965696 | 1 | 1.28E-07 | 2.40E-03 | 104807 | not applicable |
| Neutrophill_percentage | LOFTEE | ENSG00000101745 | ANKRD12 | 26.1568303 | 1 | 3.15E-07 | 5.91E-03 | 104807 | not applicable |
| Phosphate | LOFTEE | ENSG00000118785 | SPP1 | 35.6517972 | 1 | 2.36E-09 | 4.429824E-05 | 94729 | not applicable |
| Phosphate | LOFTEE | ENSG00000198569 | SLC34A3 | 31.6803032 | 1 | 1.82E-08 | 0.0003412656 | 94729 | not applicable |
| Platelet_count | LOFTEE | ENSG00000101162 | TUBB1 | 67.1918398 | 1 | 2.46E-16 | 4.63E-12 | 104959 | not applicable |
| Platelet_count | LOFTEE | ENSG00000096968 | JAK2 | 47.2062805 | 1 | 6.39E-12 | 1.20E-07 | 104959 | not applicable |
| Platelet_count | LOFTEE | ENSG00000145703 | IQGAP2 | 41.9604152 | 1 | 9.31E-11 | 1.75E-06 | 104959 | not applicable |
| Platelet_count | LOFTEE | ENSG00000215301 | DDX3X | 40.2219794 | 1 | 2.27E-10 | 4.26E-06 | 104959 | not applicable |
| Platelet_count | LOFTEE | ENSG00000197183 | NOL4L | 32.3219325 | 1 | 1.31E-08 | 2.45E-04 | 104959 | not applicable |
| Platelet_count | LOFTEE | ENSG00000175792 | RUVBL1 | 30.9334654 | 1 | 2.67E-08 | 5.01E-04 | 104959 | not applicable |
| Platelet_count | LOFTEE | ENSG00000090534 | THPO | 28.4970991 | 1 | 9.38E-08 | 1.76E-03 | 104959 | not applicable |
| Platelet_count | LOFTEE | ENSG00000168769 | TET2 | 23.2428277 | 1 | 1.43E-06 | 2.68E-02 | 104959 | not applicable |
| Reticulocyte_count | LOFTEE | ENSG00000163554 | SPTA1 | 51.5215717 | 1 | 7.08E-13 | 1.33E-08 | 102867 | not applicable |
| Reticulocyte_count | LOFTEE | ENSG00000004939 | SLC4A1 | 51.4526544 | 1 | 7.33E-13 | 1.38E-08 | 102867 | not applicable |
| Reticulocyte_count | LOFTEE | ENSG00000112077 | RHAG | 44.3268156 | 1 | 2.78E-11 | 5.22E-07 | 102867 | not applicable |
| Reticulocyte_count | LOFTEE | ENSG00000159023 | EPB41 | 29.3829287 | 1 | 5.94E-08 | 0.0011152679 | 102867 | not applicable |
| SHBG | LOFTEE | ENSG00000129214 | SHBG | 320.380039 | 1 | 1.20E-71 | 2.247625E-67 | 94020 | not applicable |
| Testosterone | LOFTEE | ENSG00000129214 | SHBG | 35.5886105 | 1 | 2.44E-09 | 4.575867E-05 | 93177 | not applicable |
| Thrombocyte_volume | LOFTEE | ENSG00000145703 | IQGAP2 | 261.53494 | 1 | 7.94E-59 | 1.491224E-54 | 104958 | not applicable |

| | | | | | | | | | |
|---|---|---|---|---|---|---|---|---|---|
| Thrombocyte_volume | LOFTEE | ENSG00000205436 | EXOC3L4 | 66.9060671 | 1 | 2.85E-16 | 5.346601E-12 | 104958 | not applicable |
| Thrombocyte_volume | LOFTEE | ENSG00000101162 | TUBB1 | 64.2633158 | 1 | 1.09E-15 | 2.043838E-11 | 104958 | not applicable |
| Thrombocyte_volume | LOFTEE | ENSG00000185245 | GP1BA | 54.732222 | 1 | 1.38E-13 | 2.59E-09 | 104958 | not applicable |
| Thrombocyte_volume | LOFTEE | ENSG00000149177 | PTPRJ | 48.0381011 | 1 | 4.18E-12 | 7.85E-08 | 104958 | not applicable |
| Thrombocyte_volume | LOFTEE | ENSG00000131504 | DIAPH1 | 30.3764423 | 1 | 3.56E-08 | 0.0006680938 | 104958 | not applicable |
| Thrombocyte_volume | LOFTEE | ENSG00000163808 | KIF15 | 28.2474878 | 1 | 1.07E-07 | 0.0020043961 | 104958 | not applicable |
| Thrombocyte_volume | LOFTEE | ENSG00000169607 | CKAP2L | 28.1305916 | 1 | 1.13E-07 | 0.0021291826 | 104958 | not applicable |
| Thrombocyte_volume | LOFTEE | ENSG00000071205 | ARHGAP10 | 26.6511155 | 1 | 2.44E-07 | 4.58E-03 | 104958 | not applicable |
| Total_bilirubin | LOFTEE | ENSG00000241635 | UGT1A1 | 82.0486279 | 1 | 1.33E-19 | 2.49E-15 | 102463 | not applicable |
| Total_bilirubin | LOFTEE | ENSG00000167165 | UGT1A6 | 75.6947621 | 1 | 3.31E-18 | 6.22E-14 | 102463 | not applicable |
| Total_bilirubin | LOFTEE | ENSG00000242366 | UGT1A8 | 73.336139 | 1 | 1.09E-17 | 2.053106E-13 | 102463 | not applicable |
| Total_bilirubin | LOFTEE | ENSG00000244122 | UGT1A7 | 52.2553527 | 1 | 4.87E-13 | 9.150094E-09 | 102463 | not applicable |
| Total_bilirubin | LOFTEE | ENSG00000242515 | UGT1A10 | 48.3945993 | 1 | 3.49E-12 | 6.54E-08 | 102463 | not applicable |
| Total_bilirubin | LOFTEE | ENSG00000241119 | UGT1A9 | 37.154545 | 1 | 1.09E-09 | 2.05E-05 | 102463 | not applicable |
| Total_bilirubin | LOFTEE | ENSG00000004939 | SLC4A1 | 35.2597751 | 1 | 2.89E-09 | 5.42E-05 | 102463 | not applicable |
| Total_bilirubin | LOFTEE | ENSG00000244474 | UGT1A4 | 28.2765858 | 1 | 1.05E-07 | 1.97E-03 | 102463 | not applicable |
| Total_bilirubin | LOFTEE | ENSG00000244734 | HBB | 23.2250236 | 1 | 1.44E-06 | 2.71E-02 | 102463 | not applicable |
| Triglycerides | LOFTEE | ENSG00000084674 | APOB | 134.500528 | 1 | 4.25E-31 | 7.97E-27 | 102769 | not applicable |
| Triglycerides | LOFTEE | ENSG00000132855 | ANGPTL3 | 133.215096 | 1 | 8.11E-31 | 1.52E-26 | 102769 | not applicable |
| Triglycerides | LOFTEE | ENSG00000110245 | APOC3 | 96.5336202 | 1 | 8.77E-23 | 1.647429E-18 | 102769 | not applicable |
| Triglycerides | LOFTEE | ENSG00000110243 | APOA5 | 93.6922938 | 1 | 3.69E-22 | 6.920706E-18 | 102769 | not applicable |
| Urate | LOFTEE | ENSG00000197891 | SLC22A12 | 62.1524976 | 1 | 3.18E-15 | 5.968208E-11 | 102742 | not applicable |
| Urate | LOFTEE | ENSG00000174827 | PDZK1 | 34.6038396 | 1 | 4.04E-09 | 7.58754E-05 | 102742 | not applicable |
| Urate | LOFTEE | ENSG00000161542 | PRPSAP1 | 29.4264142 | 1 | 5.81E-08 | 0.0010905213 | 102742 | not applicable |
| Urate | LOFTEE | ENSG00000118777 | ABCG2 | 24.4547585 | 1 | 7.61E-07 | 1.43E-02 | 102742 | not applicable |
| Vitamin_D | LOFTEE | ENSG00000084674 | APOB | 25.8527521 | 1 | 3.68E-07 | 6.92E-03 | 98461 | not applicable |
| Vitamin_D | LOFTEE | ENSG00000186104 | CYP2R1 | 25.2977088 | 1 | 4.91E-07 | 9.22E-03 | 98461 | not applicable |
| c_reactive_protein | LOFTEE | ENSG00000144182 | LIPT1 | 24.1064464 | 1 | 9.12E-07 | 0.0171151944 | 102635 | not applicable |
| glycated_haemoglobin_hba1c | LOFTEE | ENSG00000163554 | SPTA1 | 89.0118438 | 1 | 3.92E-21 | 7.368751E-17 | 103043 | not applicable |
| glycated_haemoglobin_hba1c | LOFTEE | ENSG00000112077 | RHAG | 76.9327303 | 1 | 1.77E-18 | 3.321123E-14 | 103043 | not applicable |
| glycated_haemoglobin_hba1c | LOFTEE | ENSG00000159023 | EPB41 | 56.0100626 | 1 | 7.21E-14 | 1.353769E-09 | 103043 | not applicable |
| glycated_haemoglobin_hba1c | LOFTEE | ENSG00000004939 | SLC4A1 | 37.2231624 | 1 | 1.05E-09 | 1.98E-05 | 103043 | not applicable |
| glycated_haemoglobin_hba1c | LOFTEE | ENSG00000106633 | GCK | 36.4659362 | 1 | 1.55E-09 | 2.92E-05 | 103043 | not applicable |
| glycated_haemoglobin_hba1c | LOFTEE | ENSG00000146830 | GIGYF1 | 34.2396586 | 1 | 4.87E-09 | 9.15E-05 | 103043 | not applicable |

| | Individuals | Genes | Tissues | Samples | Underexpressed outliers | Non-outliers | Overexpressed outliers |
|---|---|---|---|---|---|---|---|
| Unfiltered | 325 | 14740 | 1 | 325 | 1681 | 4787910 | 909 |
| Samples with whole genomes | 311 | 14740 | 1 | 311 | 1626 | 4581633 | 881 |
| Keep only protein-coding genes | 311 | 11849 | 1 | 311 | 1471 | 3682880 | 688 |
| Remove samples with many outliers | 295 | 11849 | 1 | 295 | 983 | 3493975 | 497 |
| Keep only genes of samples that have sufficiently large expected number of reads ('mu' > 450) | 295 | 11314 | 1 | 295 | 808 | 2586111 | 384 |

|  | Individuals | Genes | Tissues | Samples | Underexpressed outliers | Non-outliers | Overexpressed outliers |
|---|---|---|---|---|---|---|---|
| Unfiltered | 253 | 16381 | 1 | 253 | 1442 | 4142056 | 895 |
| Keep only protein-coding genes | 253 | 12771 | 1 | 253 | 1231 | 3229015 | 817 |
| Remove samples with many outliers | 233 | 12771 | 1 | 233 | 811 | 2974445 | 387 |
| Keep only genes of samples that have sufficiently large expected number of reads ('mu' > 450) | 233 | 11748 | 1 | 233 | 653 | 2182453 | 302 |
